# Experimental umbrella sampling of a rotary nanomachine

**DOI:** 10.64898/2026.10.15.758702

**Authors:** Lennart J. K. Weiß, Laetitia Daubner, Niccolò Tedeschi, Petr Šulc, Friedrich C. Simmel

## Abstract

The free-energy landscape of a molecular system, which describes its free energy as a function of a few collective coordinates, governs its dynamics and function, but is challenging to determine experimentally. In molecular simulations, enhanced-sampling methods such as umbrella sampling use bias potentials to access regions of this landscape that are rarely visited by equilibrium fluctuations. Here, we transfer this concept to a single-molecule experiment: we combine electric-field biasing, *in situ* calibration of the bias potentials, and statistical reweighting to reconstruct the angular free-energy landscapes of rotary DNA nanomachines. The method resolves barriers exceeding **10 *k***_B_***T*** and detects changes in local landscape shape caused by small modifications of the stator design. A systematic analysis of bias calibration, sampling, and finite-exposure averaging establishes the conditions required for accurate reconstruction. Experimental umbrella sampling thus relates nanoscale structure to the underlying free-energy landscape and supports the physics-informed design of artificial molecular machinery.

## Main

The dynamics of molecular systems are naturally described in terms of free-energy landscapes. The relative free energies of a system’s states determine their equilibrium populations, while the barriers separating them govern the rates of transitions between them. With the development of single-molecule manipulation techniques, such landscapes have become experimentally accessible [1]. Atomic force microscopy as well as optical and magnetic tweezers have been used to probe free-energy landscapes along defined reaction coordinates for processes including protein and DNA hairpin folding, DNA supercoiling, and strand displacement [2–6].

Advances in nucleic-acid nanotechnology and protein design enable the bottom-up construction of dynamic molecular devices such as switches, rotary motors, and walkers [7–12]. As the field progresses from designing molecular structure toward engineering function, it becomes increasingly important to understand how structure shapes the underlying free-energy landscape. In theoretical models of molecular machines, the shape of this landscape – in particular its barriers and asymmetry – determines whether and how fast a machine performs directed motion under nonequilibrium driving [13–17].

For rotary nanomachines, the landscape is described by the free energy *G*(*φ*) as a function of the angular position *φ* of the rotor relative to the stator. In thermal equilibrium, rotor orientations are distributed according to *p*(*φ*) ∝ exp[−*G*(*φ*)*/k*_B_*T*], where *k*_B_*T* is the thermal energy. Regions of high free energy are therefore exponentially less populated and remain poorly resolved in passive trajectories [18, 19]. This is a rare-event sampling problem familiar from statistical physics. In molecular simulations, it is addressed by enhanced-sampling methods such as umbrella sampling, which introduce bias potentials to populate otherwise rarely sampled regions of configuration space and recover the unbiased free-energy landscape by reweighting the resulting biased distributions [20–22].

In this work, we transfer this strategy to a single-molecule experiment. Electric fields applied along different directions bias the rotor of a DNA nanomachine toward angular regions that are rarely visited in thermal equilibrium, while *in situ* calibration determines the experimentally realized bias potentials. The rotor moves over a surface-tethered stator whose design shapes the free-energy landscape *G*(*φ*), and its angular position is tracked by total internal reflection fluorescence microscopy (TIRFM). The resulting biased angular distributions are combined using the multistate Bennett acceptance ratio (MBAR) to reconstruct *G*(*φ*) across the full 2*π* range. We further establish the experimental conditions required for accurate reconstruction and use the method to resolve barrier regions and design-dependent landscape features that remain poorly sampled during passive diffusion.

### Experimental umbrella sampling

To realize experimental umbrella sampling, we designed a rotary DNA origami system in which the rotor can be biased by an external electric field and its angular position tracked at the single-molecule level (Fig. 1a and Fig. S1). The system comprises a two-layer stator plate with a flexibly connected symmetric rotor arm and a second six-helix-bundle structure that extends the rotor and carries a fluorescently labeled tip for angular tracking. The stator is rendered asymmetric by omitting selected crossovers between the upper and lower stator layers, which creates flexible flaps that remain connected to the lower layer through single-stranded DNA tethers. Transmission electron microscopy confirms the assembly of the rotor–stator structure together with the extended rotor arm (Fig. 1b).

**Fig. 1.**
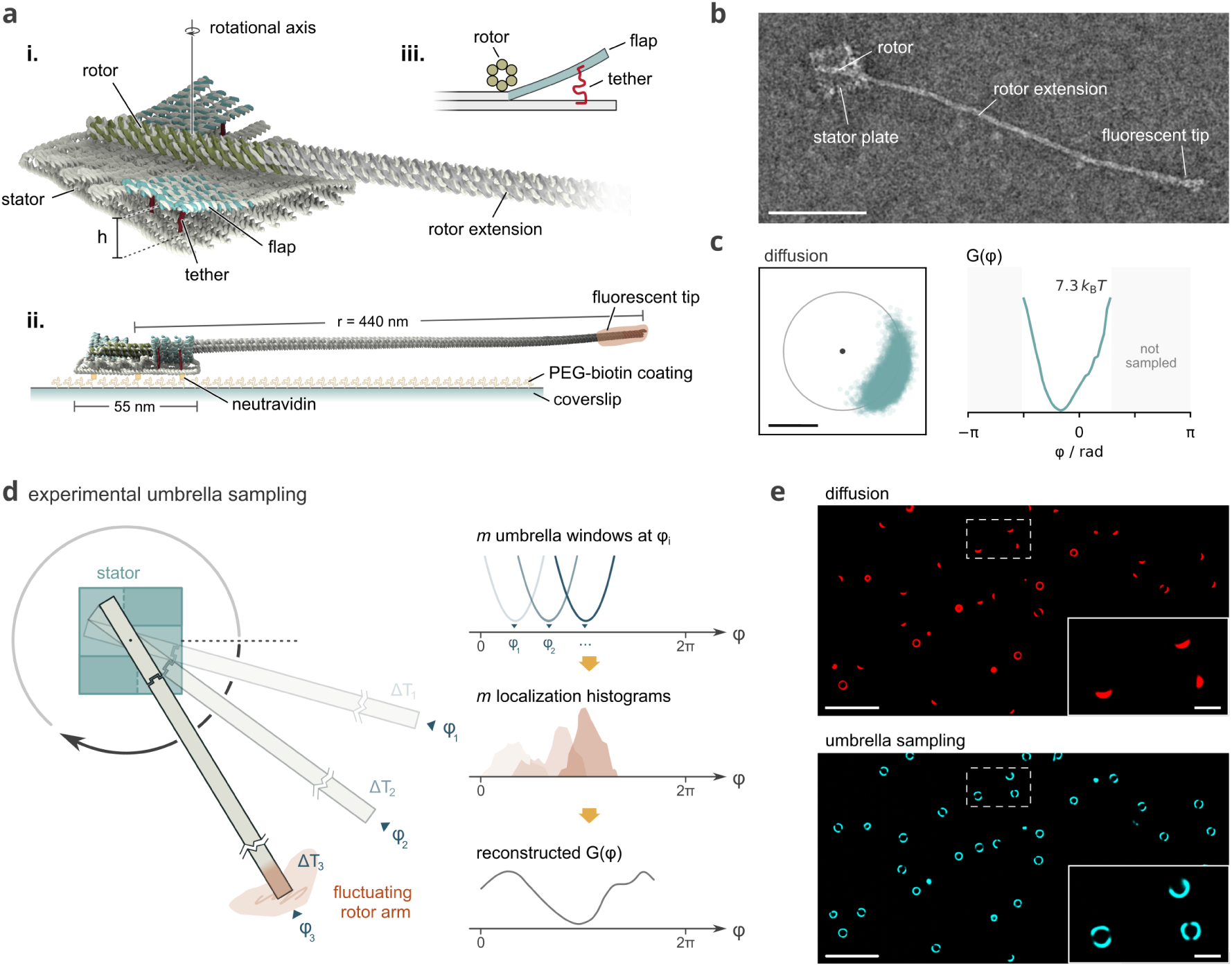
Experimental umbrella sampling of a rotary DNA origami system. **a** Design of the rotor–stator system. **i**, oxDNA representation of the two-layer stator plate, flexibly connected rotor, and elongated rotor arm (see Fig. S1). **ii**, Side view of the structure, anchored to a PEG–biotin-coated coverslip via biotin–neutravidin binding, with a fluorescently labeled tip for optical tracking. **iii**, Schematic cross-section of the asymmetric stator topology, in which the flexible flaps remain connected to the lower stator layer by a single-stranded tether. **b** TEM micrograph of an assembled rotor–stator structure with extended rotor arm. Scale bar, 100 nm. **c** Free diffusion of an individual rotor recorded for 180 s, shown as (*x, y*) localizations (left) and the corresponding free-energy profile *G*(*φ*) (right). The trajectory remains confined to the basin of a single minimum, while states *>* 7.3 *k*_B_*T* remain unsampled. Scale bar, 300 nm. **d** Principle of experimental umbrella sampling. Successively shifted electric-field biases generate *m* overlapping umbrella windows centered at *φ_i_*, yielding overlapping angular localization distributions *p_i_*(*φ*) that collectively span the rotational coordinate and can be combined to reconstruct *G*(*φ*). **e** Representative fluorescence localizations of a rotor ensemble during 3 min of free diffusion and during an umbrella-sampling protocol with a steering voltage of 300 V. Scale bars, 5 µm and 1 µm, respectively.

The asymmetric stator geometry gives rise to a non-uniform free-energy landscape *G*(*φ*). During passive diffusion, however, individual rotors explore only part of this landscape within the available observation time (Fig. 1c). Boltzmann inversion of the measured angular distribution, *G*(*φ*) = −*k*_B_*T* ln *p*(*φ*)+*C*, shows that the rotor remains confined to the basin of a single minimum, so that substantial parts of the landscape, including the barrier regions, are not sampled by thermal fluctuations alone.

Umbrella sampling overcomes this limitation by applying a bias potential that shifts the system toward states of higher free energy (Fig. 1d). In umbrella window *i* (*i* = 1*,…,m*), we apply an external bias potential *U_i_*(*φ*), with its minimum at angle *φ_i_*. The angular distribution *p_i_*(*φ*) measured in this window is determined by the sum of the intrinsic rotor–stator free-energy landscape *G*(*φ*) and the bias potential,

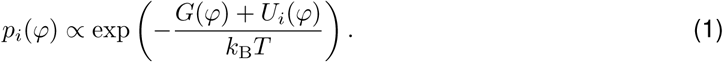

By successively shifting the bias center through *φ*_1_*,…,φ_m_*, we obtain *m* umbrella windows that together cover the full 2*π* range. Provided that the distributions of neighboring windows overlap and the bias potentials *U_i_*(*φ*) are known, the biased distributions can be reweighted to recover *G*(*φ*). Here, we combine the distributions using MBAR [22, 23].

In molecular-dynamics simulations, umbrella sampling is commonly implemented with harmonic restraint potentials. In our experiments, the bias is instead generated by an externally applied electric field acting on the rotor [8, 24], which results in a cosine-shaped potential, *U_i_*(*φ*) = *κ_i_* (1 − cos (*φ* − *φ_i_*)), that approaches a harmonic restraint close to its minimum, *U_i_*(*φ*) ≈ 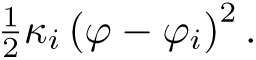 Here, *κ_i_* denotes the bias strength in window *i*, which equals the angular trap stiffness in the harmonic limit. Whereas the restraint strength can be chosen freely in simulations, the experimental bias strength is limited by the attainable electric-field-induced torque [24]. We therefore first tested whether the bias attainable in our setup is strong enough to drive the rotors out of their free-energy minima. As shown in Fig. 1e, at an applied voltage of 300 V the majority of rotors covered the full angular range, whereas during passive diffusion most rotors remained confined.

### *In situ* bias inference enables free-energy reconstruction

In addition to a sufficiently strong bias, quantitative reconstruction requires accurate knowledge of the bias potential experienced by the rotor. This is a central difference between computational and experimental umbrella sampling. In simulations, the bias potential is defined by the user and is therefore known exactly, whereas in the experiment only the applied voltage and the nominal field direction are set. The local electric field at the rotor can vary from experiment to experiment owing to differences in buffer conditions, electrode geometry, and the position of the field of view. As a result, the center of the realized bias potential can deviate from its nominal position, and its strength is not known *a priori*. The bias potential therefore cannot be inferred from the applied voltage alone but must be determined separately for the conditions of each experiment.

We therefore developed an *in situ* calibration strategy in which the asymmetric rotor–stator structures, hereafter referred to as *target rotors*, are measured alongside structurally distinct *calibration rotors* within the same field of view (Fig. 2a). The calibration rotors consist of six-helix-bundles directly immobilized on the PEG–biotin-coated coverslip and thus lack the structured stator interactions of the target rotors, resulting in an approximately flat intrinsic angular free-energy landscape [18] that allows them to serve as local probes of the electric-field-induced bias.

**Fig. 2.**
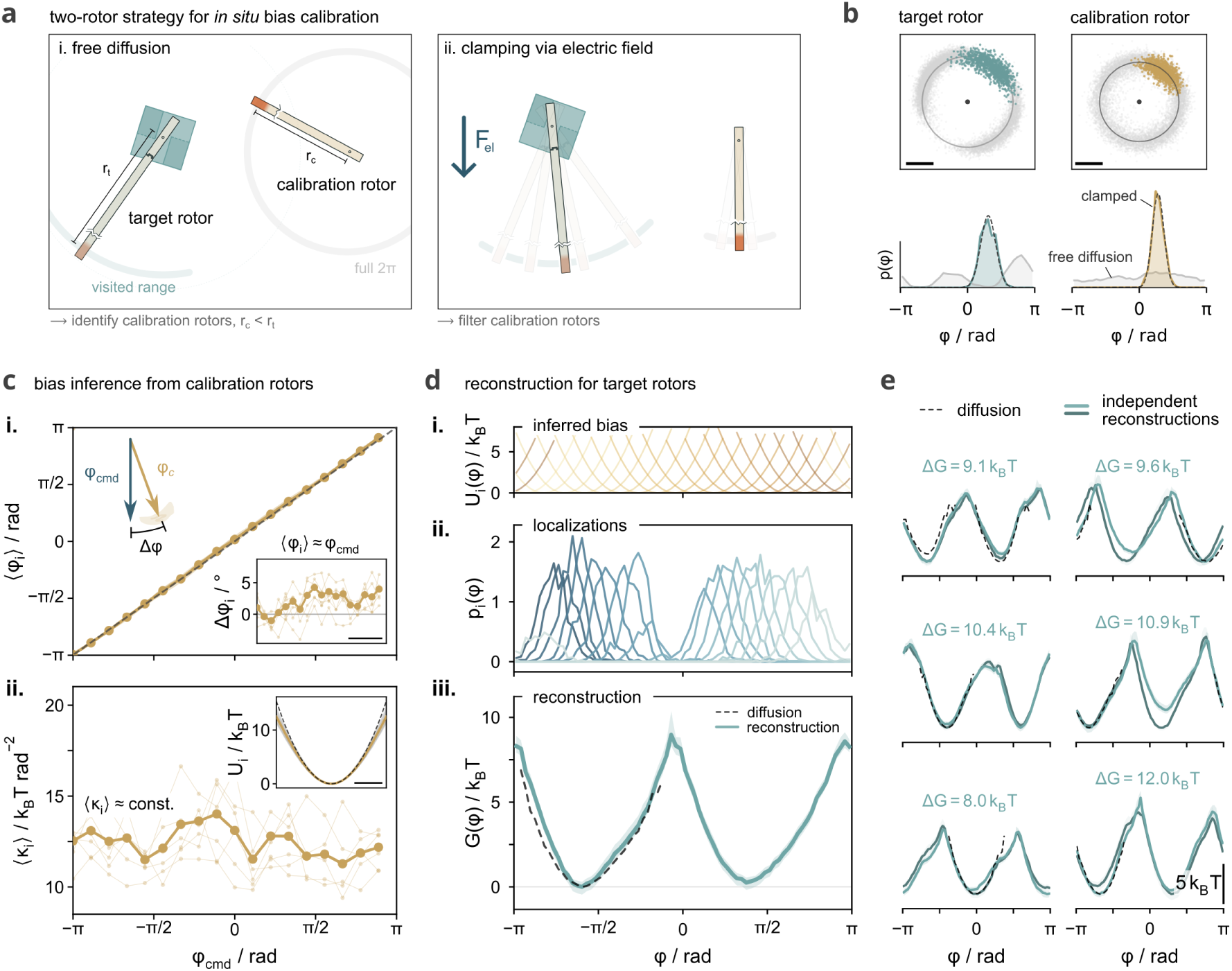
*In situ* calibration enables quantitative reconstruction of free-energy landscapes. a. Two-rotor strategy for local bias calibration. Calibration rotors are recorded alongside target rotors within the same field of view and identified by their smaller tracking radii (*r*_c_ = 375 nm and *r*_t_ = 440 nm, respectively) and approximately flat bias-free energy landscape. **b** Localization patterns and corresponding angular probability distributions *p*(*φ*) of exemplary calibration and target rotors during bias-free diffusion and during a 5 s umbrella window at 300 V. Scale bar, 200 nm. **c** Bias inference across successive umbrella windows. **i**, calibration-rotor ensemble-averaged window centers *⟨φ_i_⟩*. Inset, steering error Δ*φ* between commanded and realized window centers. **ii**, bias strengths *⟨κ_i_⟩* used to define the restraint potentials for reconstruction. Inset, mean cosine bias potential and its harmonic approximation (dashed). Scale bars, *π/*2 in **i** and *π/*4 in **ii**. **d** Energy-landscape reconstruction for an exemplary target rotor. **i**, inferred restraint potentials. **ii**, recorded angular probability distributions *p_i_*(*φ*). **iii**, MBAR-derived free-energy landscape *G*(*φ*) together with the bias-free profile (dashed). Shading denotes the 95% confidence interval from block-bootstrap resampling. **e** Paired free-energy landscapes from two independent umbrella-sampling runs for exemplary target rotors, compared with free-diffusion profiles (dashed). Barrier heights Δ*G* are indicated for each rotor.

To distinguish the two populations, calibration and target rotors were designed such that their fluorescently labeled tips trace circles of different radii, *r*_c_ = 375 nm and *r*_t_ = 440 nm, respectively, which are directly apparent in the localization patterns (Fig. 2a). As expected, the calibration rotors show a nearly uniform angular distribution during passive diffusion, with free-energy variations Δ*G <* 1 *k*_B_*T*, in contrast to the confined angular motion of the target rotors caused by rotor–stator interactions (Fig. 2b and Figs. S2–S4). At an applied voltage of 300 V (in a high-viscosity buffer with *η* = 12.5 *η*_water_), the calibration rotors become strongly localized around the bias center, with an approximately Gaussian angular distribution, as illustrated for a window with an angular trap stiffness of *κ_i_* = 13.4 *k*_B_*T* rad*^−^*^2^ (≈ 55 pN nm rad*^−^*^2^). From such distributions, we determined the bias center and trap stiffness for each calibration rotor and umbrella window. Their averages over the calibration-rotor ensemble, ⟨*φ_i_*⟩ and ⟨*κ_i_*⟩, define the bias potentials *U_i_*(*φ*) used for reconstruction (Fig. 2c). Importantly, reconstruction does not require these parameters to remain constant across windows, as long as the realized bias potential of each window is known. In the experiment shown in Fig. 2c, the realized bias centers stayed within |Δ*φ*| *<* 5*^◦^* of the nominal field directions *φ*_cmd_(*t*), while ⟨*κ_i_*⟩ remained approximately constant at 12.5 *k*_B_*T* rad*^−^*^2^, indicating stable conditions throughout.

The different arm lengths of calibration and target rotors, however, result in different electric-field-induced torques, because the torque depends on the distribution of charge relative to the rotational axis [24, 25]. The trap stiffness measured with the calibration rotors, *κ*_c_, therefore cannot be transferred directly to the target rotors but must be rescaled to obtain their stiffness *κ*_t_. We assume a uniform effective line charge density *λ* along the six-helix bundles. For a rotor at angle *φ* in an inplane electric field of magnitude *E* directed along *φ_i_*, the field component perpendicular to the rotor is *E_⊥_* = *E* sin(*φ* − *φ_i_*). The rotational axis divides each rotor into a long and a short arm segment of lengths *x*_1_ and *x*_2_ (Fig. 1a), whose contributions to the torque have opposite sign, so that the net torque is

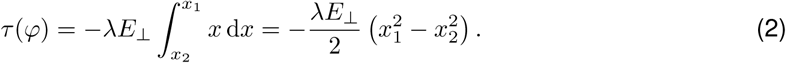

Comparison with *τ* (*φ*) = −d*U_i_/*d*φ* = −*κ_i_* sin(*φ* − *φ_i_*) yields 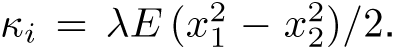 Since *λ* and *E* are the same for both rotor types within a field of view, *κ*_t_ and *κ*_c_ differ only by the geometric factor *x*_1_^2^-*x*_2_^2^. With arm lengths of 440 nm and 25 nm for the target rotors and 375 nm and 40 nm for the calibration rotors, we obtain *κ*_t_ = 1.39 *κ*_c_. The bias potential of the target rotors in window *i* thus has its center at ⟨*φ_i_*⟩ and the stiffness 1.39 ⟨*κ_i_*⟩, both determined *in situ* from the calibration rotors.

Using these calibrated bias potentials, we combine the target-rotor localizations from all umbrella windows and reconstruct the free-energy landscape of each individual rotor with MBAR (Fig. 2d). In these experiments, we used 18 umbrella windows with centers 20*^◦^* apart, each recorded for 5 s at an applied voltage of 300 V, corresponding to a total acquisition time under bias of 90 s. The reconstructed landscapes in Fig. 2d,e reveal a pronounced asymmetry – consistent with the asymmetric stator geometry – and free-energy barriers of about 10 *k*_B_*T*. Importantly, the reconstructed landscapes closely reproduce the profiles obtained from passive diffusion within the angular range sampled there. To assess reproducibility, we performed a second umbrella-sampling run on the same target rotors. The paired reconstructions in Fig. 2e agree well (root-mean-square error, RMSE, typically below 1 *k*_B_*T*), with the main landscape features and barrier heights consistently recovered (Figs. S5–S8).

We further quantified the statistical uncertainty of individual reconstructions by block bootstrapping. We resampled the calibration-and target-rotor trajectories in 250 ms blocks and reconstructed the landscape for each of 100 bootstrap replicates. The resulting 95% confidence intervals (shading in Fig. 2d,e) typically amount to 0.1–0.3 *k*_B_*T* and are thus small compared with the free-energy barriers of about 10 *k*_B_*T* (Fig. S9).

Experimental umbrella sampling has the same statistical requirements as its computational counterpart: each umbrella window must be sufficiently sampled, and the distributions of neighboring windows must overlap (Figs. S10–S16). In addition, the realized bias potential has to be determined reliably from the experiment itself. We therefore tested how sensitively the calibration depends on the number of calibration rotors. Even with a single calibration rotor, the center and stiffness of the bias potential deviated only modestly from the values obtained with the full ensemble, and the reconstructed landscapes differed from the full-ensemble reconstruction by an RMSE of approximately 0.4 *k*_B_*T* on average (Figs. S17 and S18). A small number of calibration rotors is thus sufficient for a consistent calibration.

The experimental conditions considered so far – an applied voltage of 300 V, 18 windows with centers 20*^◦^* apart, 5 s per window, and a high-viscosity buffer – yielded reproducible *G*(*φ*), but the accuracy of these estimates depends on how the applied voltage, acquisition time, window spacing, and camera exposure time affect confinement, sampling, window overlap, and temporal resolution of the biased trajectories. We therefore systematically varied these parameters to establish the resulting constraints on experimental umbrella sampling.

### Requirements for accurate reconstruction

We first examined how the available acquisition time should be divided between the acquisition time per window and the number of windows. From a purely statistical perspective, a longer acquisition time per window improves the estimate of the stationary angular distribution, while a smaller window spacing increases the overlap between neighboring windows (Figs. S19 and S20). Experimentally, however, the total acquisition time under bias cannot be increased arbitrarily, because prolonged electric-field exposure and fluorescence imaging increase the risk of drift, electrochemical effects, Joule heating, and photobleaching [9, 19]. For a fixed total acquisition time, the optimal balance between the number of windows and the acquisition time per window depends on the free-energy landscape and on the relaxation time, that is, the time the rotor needs to equilibrate within a window. In our experiments, reducing the acquisition time from 5 s to 2.5 s per window in favor of a smaller window spacing resulted in pronounced reconstruction errors, whereas a larger window spacing produced considerably smaller deviations, with an average RMSE *<* 1 *k*_B_*T* (Figs. S21–S23). Thus, as long as neighboring windows retained sufficient overlap, a smaller window spacing provided little additional improvement in the reconstructed *G*(*φ*). Instead, the acquisition time per window had to be long enough for the rotor to approach the stationary biased distribution. This was most critical near the free-energy barriers, where the relaxation time was substantially longer than elsewhere (Fig. S15).

As the second parameter, we investigated the influence of the applied voltage, which determines the stiffness of the bias potential. The bias must be strong enough to give access to regions of high free energy, while the overlap between neighboring windows must be maintained. To examine this trade-off, we performed umbrella-sampling runs on the same field of view at 200, 250, and 300 V (Figs. S24–S27). As shown in Fig. 3a, at voltages below 300 V the calibration rotors still follow the field direction, whereas the target rotors increasingly lag behind the window centers because the rotor–stator interactions oppose the applied torque. Increasing the voltage increases the stiffness ⟨*κ_i_*⟩ and confines the rotors more tightly, but simultaneously narrows the biased distributions and thereby reduces the overlap *O_i,i_*_+1_ between neighboring windows (Fig. 3b). Accordingly, the angular lag ⟨|Δ*φ_i_*|⟩ of the target rotors decreased with increasing voltage, while the number of recorded localizations near the barrier, *N*_eff,b_, increased (Fig. 3c). Sufficient sampling of these regions of high free energy is critical for accurate reweighting. At 200 V, the barriers remained sparsely sampled, which compromised the reconstruction and led to overestimated barrier heights, whereas the reconstructions at 250 and 300 V nearly coincided (Fig. 3d and Figs. S28–S29).

**Fig. 3.**
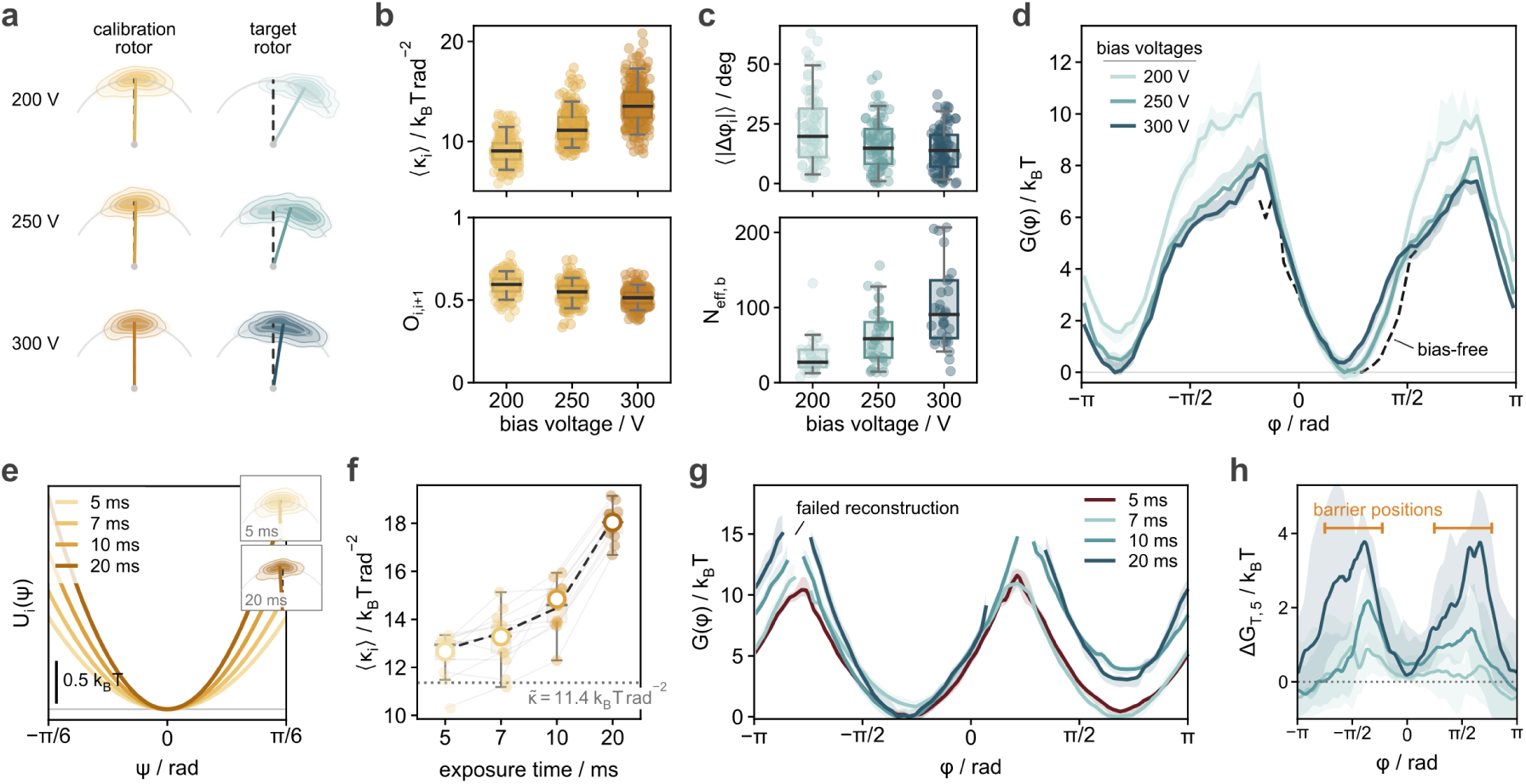
Practical limits for experimental umbrella sampling. **a** Exemplary angular distributions of calibration and target rotors during umbrella sampling at steering voltages of 200, 250, and 300 V. **b** Voltage dependence of the bias strength *⟨κ_i_⟩* (top) and overlap *O_i,i_*_+1_ between neighboring umbrella windows (bottom) for calibration rotors. **c** Steering error *⟨|*Δ*φ_i_|⟩* of target rotors (top) and effective number of samples obtained near the target-rotor barrier regions, *N*_eff,b_ (bottom) as function of voltage. **d** Reconstructed *G*(*φ*) for an exemplary target rotor from independent umbrella-sampling measurements at 200, 250, and 300 V. **e** Apparent restraint harmonic potentials *U_i_*(*φ*) obtained at camera exposure times of 5, 7, 10, and 20 ms from a single calibration rotor under otherwise identical steering conditions. Insets show the angular distribution for a single umbrella window. **f** Inferred bias strength as function of the camera exposure time. Open symbols indicate the calibration-rotor ensemble response. The dotted curve displays the underlying restraint stiffness *κ̃* obtained from the Ornstein–Uhlenbeck model (dashed fit) in the limit of vanishing exposure time. **g** Reconstructed *G*(*φ*) of an exemplary target rotor from measurements acquired at different camera exposure times. **h** Exposure-dependent reconstruction deviations across target rotors, quantified as local energy difference Δ*G_T,_*_5_ relative to the 5 ms reference reconstruction. Individual energy landscapes were aligned to a consensus with a minimum located at *φ* = 0.

As the third parameter, we examined the camera exposure time. Each localization is an average of the rotor position over the exposure time rather than an instantaneous angular position. When the exposure time approaches the relaxation time of the rotor in the bias potential, temporal averaging narrows the measured angular distribution. For the calibration rotors, this effect can be described quantitatively. Close to the minimum of the cosine-shaped bias potential, their dynamics correspond to an overdamped Ornstein–Uhlenbeck process (Supplementary Notes 7.1), for which averaging over the exposure time reduces the variance of the measured positions. Since a narrower distribution corresponds to a stiffer potential, the stiffness inferred from the calibration rotors is overestimated. To test this prediction, we repeated the same electric-field protocol and varied only the camera exposure time (5, 7, 10, and 20 ms). As predicted, longer exposure times progressively narrowed the measured calibration-rotor distributions, corresponding to increasingly steep apparent bias potentials (Fig. 3e) and larger apparent stiffness values (Fig. 3f). Fitting the Ornstein–Uhlenbeck model to these data yields the relaxation time *τ̃* and the stiffness *κ̃* in the limit of vanishing exposure time. Across the calibration-rotor ensemble, we obtained *τ̃* ≈ 12.5 ms.

Our standard exposure time of 5 ms is thus shorter than *τ̃* but not negligible compared with it, and the stiffness obtained at 5 ms exceeds *κ̃* by approximately 14% (Fig. S30). This does not translate into a comparable error in *G*(*φ*). The error in the bias potential *U_i_*(*φ*) vanishes at the window center *φ_i_* and grows with the distance from it, whereas each window contributes to the reconstruction mainly close to its center. Because the windows overlap, every angle lies close to the center of at least one window. We therefore use the calibration obtained at 5 ms without correction and accept a small systematic error in *G*(*φ*).

For the target rotors, the effect of temporal averaging cannot be described by the same Ornstein–Uhlenbeck model, because their dynamics are additionally governed by the unknown free-energy landscape *G*(*φ*), so that relaxation time and confinement vary with *φ*. We therefore compared reconstructions obtained at different exposure times. With increasing exposure time, the angular range covered by the reconstructed *G*(*φ*) decreased, particularly around the barriers (Fig. 3g), and the reconstructions deviated substantially from the reference reconstruction at 5 ms near the barrier positions (Fig. 3h and Fig. S31). Because the acquisition time per window was fixed, longer exposure times also resulted in fewer localizations, with four times fewer at 20 ms than at 5 ms. However, the deviations persisted when the number of localizations was equalized across exposure times, which shows that they arise primarily from temporal averaging rather than from the smaller number of samples (Figs. S32 and S33).

These experiments establish three practical requirements for an accurate reconstruction of the free-energy landscape: an acquisition time per window that is long enough for the rotor to equilibrate, a bias that is strong enough to give access to regions of high free energy while the window overlap is maintained, and a camera exposure time that is short compared with the relaxation time of the rotor. For the landscape studied here, we therefore favored a longer acquisition time per window over a smaller window spacing. The exposure-time requirement can be quantified by the ratio of the exposure time *t*_e_ to the relaxation time, with temporal averaging becoming increasingly relevant as this ratio approaches one. In our high-viscosity buffer, *t*_e_ = 5 ms and *τ̃* ≈ 12.5 ms give a ratio of 0.4, for which the effect remained small. Changes in buffer conductivity and viscosity alter both the attainable bias and the relaxation time [24], so that these experimental parameters have to be adjusted accordingly.

### Stator design shapes the free-energy landscape

We next applied experimental umbrella sampling to determine how the design of the stator shapes the free-energy landscape of the rotor. In our structure, the rotor moves across flexible flaps that follow the helix direction of the stator plate (Fig. 1a). Clockwise and counterclockwise motion therefore take different paths across a flap, which results in an asymmetric free-energy landscape. Such asymmetry is particularly relevant for nanoscale machinery because it can provide the symmetry breaking required to rectify Brownian motion under nonequilibrium driving [13, 15, 26, 27]. To modify the landscape without changing the overall rotor–stator structure, we repositioned the single-stranded DNA tethers that connect each flap to the lower stator layer. This yielded three stator designs, which we termed the outer-, dual-, and inner-anchor variants (Fig. 4a; the outer-anchor variant was used in the experiments described above). oxDNA simulations predicted that the tether positions alter the equilibrium conformation of the flaps, with median flap heights of approximately 6.0, 6.5, and 7.0 nm for the outer-, dual-, and inner-anchor variants, respectively (Fig. 4b).

**Fig. 4.**
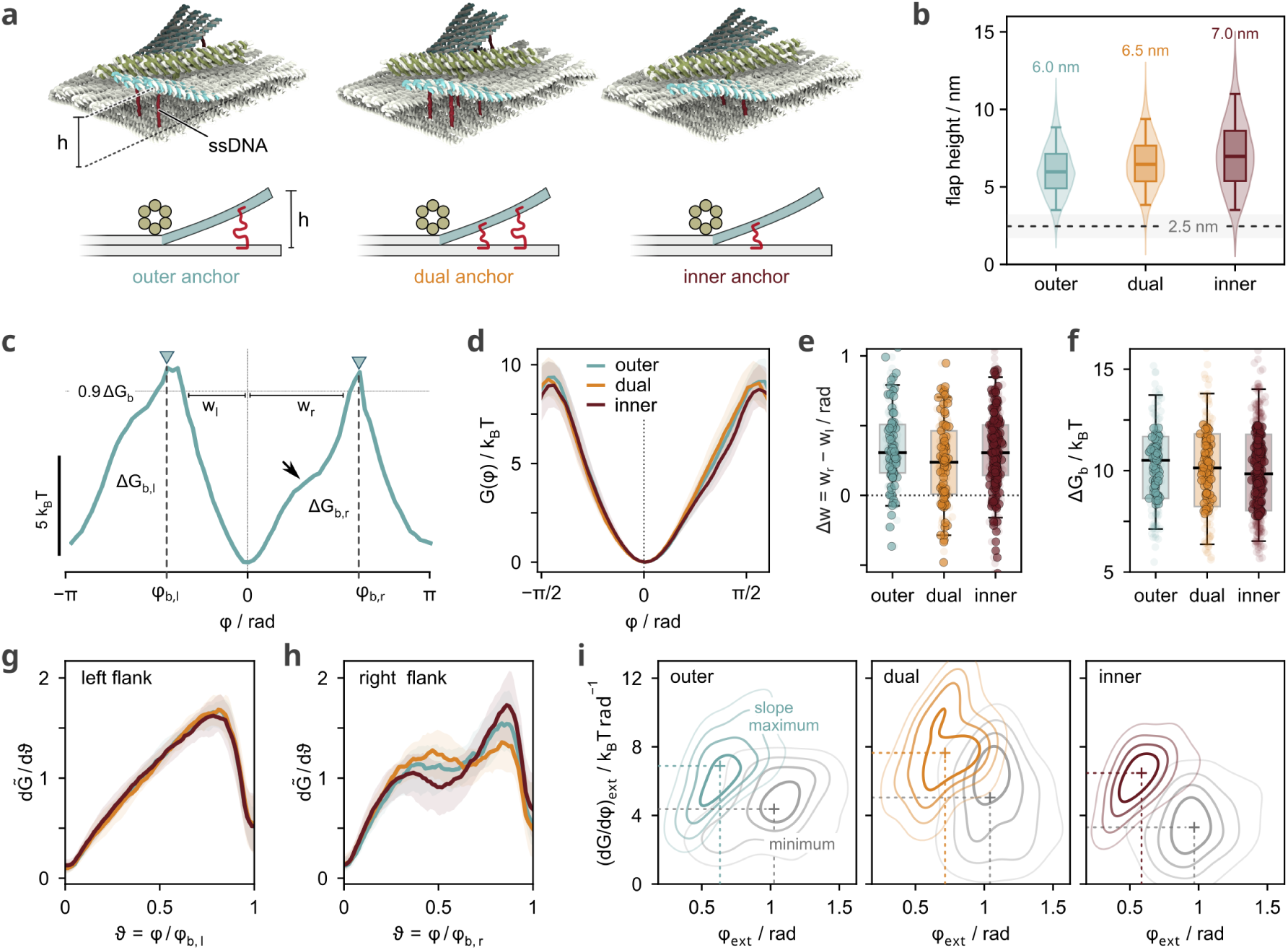
Stator design-dependent energy landscapes. **a** oxDNA illustrations and schematic side views of the outer-, dual-, and inner-anchor stator variants. **b** Flap-height distributions for the three stator variants, measured as the distance between the top and bottom layers as shown in **a**. Boxes indicate the interquartile range and median. The dashed line marks the 2.5 nm height obtained for a stator plate without flaps, due to fraying and thermal fluctuations. **c** Definition of the free-energy landscape metrics. Δ*G_b,l_* and Δ*G_b,r_* denote the left and right barrier heights, while the angular distances *w_l_* and *w_r_* are determined at 90% of the lower barrier height. The arrow highlights the characteristic local slope feature evaluated in panels **h** and **i**. **d** Consensus free-energy landscapes *G*(*φ*). **e** Distribution of barrier asymmetries Δ*w* = *w_r_ − w_l_*. **f** Distribution of the barrier height. For each rotor, Δ*G_b_* is the mean of the left and right barrier heights. **g,h** Slope *dG̃/dϑ* of the normalized left (**g**) and right (**h**) flanks, with energy normalized by the respective barrier height and angle by the corresponding minimum-to-barrier distance. **i** Joint distributions of the positions and slope values of the local extrema of the right-flank slope, characterizing the slope-flattening feature. Colored and gray contours denote maxima and minima, respectively, and crosses mark ensemble medians. Analyses in **d** and **g**–**i** use rotors within the 20th–80th percentile of the barrier-height distribution shown in **f** (*n* = 81, 76, and 174 rotors for the outer-, dual-, and inner-anchor stator variants, respectively). Data in **e** and **f** are based on *n* = 135, 126, and 290 rotors, respectively.

All three variants were measured under the conditions established above: an applied voltage of 300 V, a high-viscosity buffer, window centers 20*^◦^* apart, and an acquisition time of 5 s per window. We assume that the electric field acts predominantly on the rotor and does not substantially alter the conformation of the stator, because a field-induced deformation of the flaps is limited by their bending stiffness and by the restoring forces of the single-stranded DNA tethers.

To compare the variants, the landscape of each rotor was shifted such that its global minimum lies at *φ* = 0 and was characterized by the barrier height Δ*G_b_*, the mean of the left and right barrier heights, and by the width asymmetry Δ*w* = *w_r_* − *w_l_*, where *w_l_* and *w_r_* are the widths of the left and right flanks as defined in Fig. 4c (see also Figs. S34–S39). Despite substantial rotor-to-rotor heterogeneity, the consensus landscapes of the three variants differed slightly in their overall shape (Fig. 4d–f). All three variants exhibited a positive width asymmetry, Δ*w >* 0, corresponding to a broader right than left flank (Fig. 4e), and Δ*w* differed between the variants (Kruskal–Wallis test, *p <* 0.05). In addition, some landscapes showed a shoulder on the right flank, that is, a region in which the slope is locally reduced (arrow in Fig. 4c).

The barrier heights were similar for the three variants, with medians of 10.49, 10.13, and 9.84 *k*_B_*T* for the outer-, dual-, and inner-anchor variants, respectively (Fig. 4f). The systematic variation in equilibrium flap height predicted by oxDNA was thus not reflected in the barrier height. A possible reason is that barrier crossing depends on the mechanical response of the flap rather than on its equilibrium position alone. In addition, rotor-to-rotor heterogeneity and nonspecific surface interactions may obscure subtle effects [28].

To compare local features independently of barrier height and width, the two flanks extending from the minimum to the respective barriers were normalized separately. For each flank, we define the normalized free energy *G̃* as *G* divided by the barrier height, and the normalized angle *ϑ* as the angular distance from the minimum divided by the distance from the minimum to the barrier, so that both quantities run from 0 at the minimum to 1 at the barrier. The slopes *dG̃/dϑ* of the normalized flanks were then compared for the three variants (Fig. 4g,h). The left flank was largely the same for the three variants, whereas the right flank differed between them, in particular in the shoulder described above. We quantified the shoulder by the positions and slope values of the corresponding local extrema of the slope (Fig. 4i). Both quantities differed between the three variants (Kruskal–Wallis test, *p <* 0.05), with the stronger differences in the slope values. For both extrema, the median position was closest to the minimum for the inner-anchor variant. Experimental umbrella sampling thus resolved differences in the local free-energy gradient that result from comparatively small changes in the stator design.

The slope of the landscape has a direct mechanical meaning: it is the torque that the stator exerts on the rotor, *τ* (*φ*) = −d*G/*d*φ*. The shoulder thus marks an angular range in which the restoring torque is reduced. A comparison of the variants shows that the tether positions change the torque profile on the right flank but not on the left. Moreover, the torque is larger on the steeper left flank (Fig. 4d), which the rotor climbs during clockwise motion. This is somewhat unexpected, as clockwise motion follows the helix direction of the flaps, along which they should be displaced more readily.

The reconstructed landscape therefore suggests a preference for counterclockwise rotation, contrary to the intuitive expectation based on flap geometry. To test this experimentally, we applied a rotating electric field at increasing voltages, alternating between clockwise and counterclockwise rotation at 1 Hz. The resulting rotor trajectories indeed favor counterclockwise motion (Fig. **??**), demonstrating that the reconstructed free-energy landscape can predict directional dynamics that are not immediately apparent from the stator geometry alone.

## Discussion

This work transfers umbrella sampling, a method widely used in molecular simulations, to a single-molecule experiment for reconstructing free-energy landscapes beyond the regions efficiently explored by equilibrium fluctuations. By combining external biases, *in situ* calibration of the bias potentials, and statistical reweighting, we reconstruct the angular free-energy landscapes of individual rotary nanomachines, including barrier regions that are sparsely sampled during passive diffusion. Applied to different stator designs, the method reveals mechanical responses that are not readily inferred from structure alone.

Unlike in simulations, the bias potential in an experiment is not known *a priori*, and the measurement has a finite temporal resolution. We addressed both issues: calibration rotors measured in the same field of view provide the bias potential *in situ*, and a systematic variation of acquisition time, applied voltage, and camera exposure time established the conditions under which the biased trajectories can be reweighted quantitatively. Under these conditions, the method resolved differences between landscapes that result from small modifications of the stator design. More generally, the approach requires that the applied bias changes the probabilities of the sampled states without substantially altering the intrinsic landscape itself. An actuation that acts on the rotor alone, for example through magnetic nanoparticles bound to the rotor arm, would leave the stator unaffected and thus preserve the intrinsic landscape more closely, and would extend experimental umbrella sampling beyond the electric-field-based implementation demonstrated here [28].

The approach is not restricted to DNA nanostructures. It requires only a coordinate that can be biased and tracked, and a means of calibrating the applied bias *in situ*. It is therefore also applicable to single-molecule assays of biological or artificial molecular motors in which external torques or forces are applied through attached probes. For the design of artificial machines, the method addresses a specific need: a Brownian motor generates directed motion by switching between free-energy landscapes, and its directionality and efficiency are determined by the barriers and asymmetries of these landscapes [15, 16]. Measuring the landscape of each switching state, for example with and without a bound effector, would allow the performance of a motor design to be rationalized and thus potentially optimized.

While the function of a molecular machine is governed by its free-energy landscape, artificial nanomachines are still designed primarily in terms of structure, i.e., the shape of their components and the way these are connected. Experimental umbrella sampling bridges the gap between structural design and free-energy landscape: it reveals the energetic consequences of a given design and thereby connects molecular structure to predictive physical models of dynamics and function.

## Methods

### Preparation of rotary DNA origami structures

The target rotor comprises two DNA origami structures: a stator–rotor unit and a rotor extension for TIRFM-based tracking (Fig. S1), folded from 8,064-nt and 7,249-nt scaffolds, respectively. The stator plate consists of two stacked 55 nm×55 nm layers and is connected through a 3-nt flexible joint to a 50 nm proximal rotor segment, which is extended by a 415 nm-long six-helix bundle carrying 42 Atto655 fluorophores for TIRFM tracking. Structural asymmetry in the stator plate was introduced by two semi-mobile flaps of approximately 20 nm×30 nm at diagonally opposed stator corners. The inner-and outer-anchor variants connect these flaps to the bottom layer of the plate through single-stranded 15-nt inner and 30-nt outer tethers, respectively, whereas the dual-anchor variant contains both connections. The calibration rotor consists of a six-helix bundle with a biotin anchor positioned 40 nm from one end and was folded, functionalized, and purified following the same procedure as the target-rotor extension. The folding and purification of the DNA origami structures followed established procedures [8, 19, 24]. The 8,064-nt scaffold was enzymatically linearized with HindIII prior to folding. The stator–rotor and rotor extension were folded with a 5-fold molar excess of staple strands in 1× TE containing 5 mM NaCl and 20 mM or 12 mM MgCl_2_, respectively. Folded structures were purified by PEG precipitation followed by agarose gel electrophoresis as described previously [19]. Prior to experiments, stator–rotor modules and rotor extensions were isothermally incubated at 37 *^◦^*C at an equimolar ratio to assemble the complete target structures at a final concentration of approximately 1 nM.

### Transmission electron microscopy

TEM samples were prepared and imaged by negative-stain transmission electron microscopy as described previously [19]. Micrographs were acquired on an FEI Tecnai T12 transmission electron microscope operated at 120 kV.

### TIRF microscopy

Experiments were performed using the same custom-built TIRF microscopy and electrical steering setup with PEG–biotin-coated coverslips described previously [8, 9, 24]. Calibration and target rotors were co-incubated on the coverslip in 1× TE containing 500 mM NaCl and 0.05 % (v/v) Tween 20. Following immobilization, the electrophoretic channel was thoroughly washed with 30 mM Tris, 30 mM borate, and 3 mM MgCl_2_. For measurements, the buffer was supplemented with 45 % (w/w) sucrose to increase viscosity while maintaining low conductivity to minimize Joule heating.

Unless stated otherwise, experiments were performed at a steering voltage of 300 V and a camera exposure time of 5 ms. Each umbrella-sampling protocol comprised a 1 s passive-diffusion phase followed by 5 s umbrella windows spanning the full 2*π* range in clockwise direction, resulting in a total acquisition time of 91 s. Typically, a 60 s passive-diffusion recording was followed by two umbrella-sampling protocols per field of view, containing approximately 50–200 target rotors. To minimize effects of prolonged electrical actuation, including Joule heating and pH changes, the measurement buffer was replaced after each umbrella-sampling acquisition.

### Analysis of TIRF microscopy data

TIRFM recordings were processed in *picasso* [29] using the *localize* and *render* extensions. If required, drift correction based on sticky structures was applied prior to manual selection of rotor localization patterns. The localization data were exported to the custom Python package *roboarm-analysis* [30], where they were converted into angular trajectories by fitting the circular rotor motion and transforming the coordinates into polar coordinates. Calibration and target rotors were identified based on their radial position and trajectory characteristics, and the electric-field-induced bias potentials were inferred from the angular distributions of the calibration rotors. The unbiased free-energy landscapes of the target rotors were subsequently reconstructed from the overlapping umbrella windows using an MBAR approach [22, 23] adapted from the *pymbar* [31] package, while considering a cosine bias potential shape (Supplementary Notes 3.1).

### Coarse-grained molecular dynamics simulation

We performed molecular dynamics simulations of the DNA structures using oxDNA [32, 33], a coarse-grained model specifically developed to enable the simulation of large DNA systems while accurately capturing their essential mechanical properties, including the conformational dynamics of the flaps across the different variants studied. Simulations were performed at 293.15 K and at a salt concentration of 1 M. Temperature coupling was achieved using an Andersen-like thermostat, while electrostatic interactions were modeled using a Debye-Hückel potential. A diffusion coefficient of 2.5 in simulation units was employed to accelerate diffusive dynamics and enable sampling over longer timescales.

### Large language model tools

We used the online tool ChatGPT [34] to assist with programming.

## Data Availability

The Jupyter notebooks used for data analysis and figure generation are available at (GitHub) and is archived at (Zenodo).

## Acknowledgments

We thank Hendrik Dietz and his group for providing in-house-produced scaffold strands, Maria Ponetsmüller for enzymatic linearization of the scaffold strands, and Christoph Karfusher for TEM imaging.

## Author Contributions

L.J.K.W. designed the study. L.D. prepared the DNA origami structures. L.J.K.W. and L.D. performed the TIRFM experiments. L.J.K.W. analyzed the data. N.T. performed and analyzed the oxDNA simulations. F.C.S. and P.S. supervised the project, acquired funding and provided resources. L.J.K.W., F.C.S and L.D. wrote the manuscript. All authors provided critical feedback.

