## Supplementary Material for "Experimental umbrella sampling of a rotary nanomachine"

**9 Rotor dynamics in response to alternating rotating electric fields** **35**

**10 DNA scaffold sequences and staples** **36**

### 1 Design of the nanoscale rotor

The stator–rotor structure was designed from an 8,064-nt scaffold as a two-layer DNA origami plate connected to a six-helix-bundle rotor through a 3-nt scaffold joint (Fig. S1a). The two 55 nm $\times$ 55 nm stator layers were arranged orthogonally, with six biotinylated staples in the bottom layer for surface immobilization (Fig. S1b,c). Two diagonally opposed regions of the top layer were partially disconnected from the bottom layer to form semi-mobile flaps. The inner-, outer-, and dual-anchor variants were defined by 15-nt inner and 30-nt outer ssDNA tethers connecting the flaps to the bottom layer, with the dual-anchor variant containing both connections. The $50\,\mathrm{nm}$-long rotor segment (Fig. S1d) is centrally connected to the stator and extended by a separate, previously established $415\,\mathrm{nm}$-long six-helix-bundle structure[1–3], assembled through shape and sequence complementarity. The extension was folded from a 7,249-nt scaffold and carried 42 Atto655 fluorophores at its distal end for TIRFM tracking.

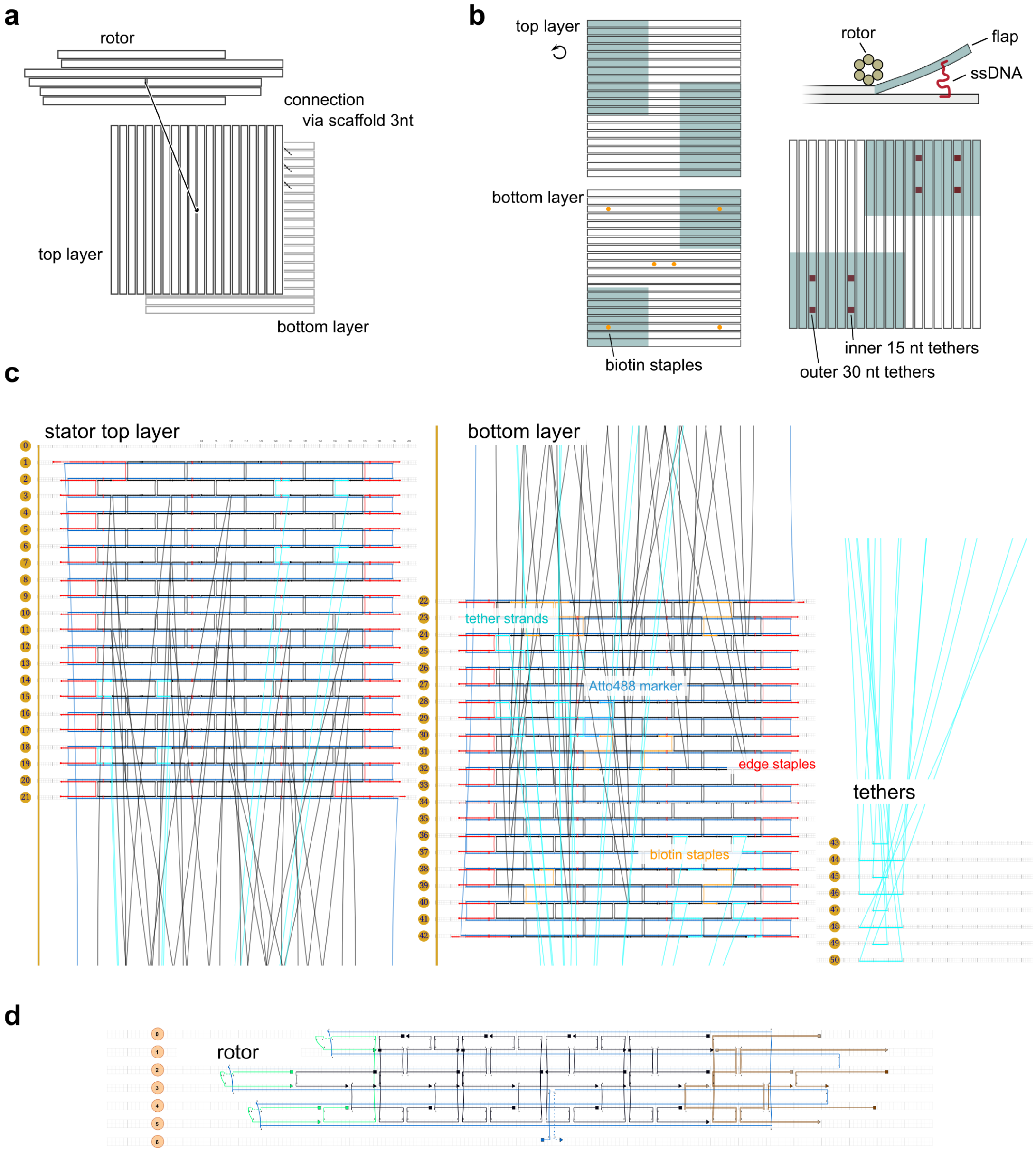

**Figure S1: DNA origami design of the stator–rotor structure.** **a** Schematic of the two-layer stator plate and centrally connected rotor. **b** Arrangement of the orthogonally stacked stator layers, surface-immobilization sites, and ssDNA tethers connecting the semi-mobile flaps to the bottom layer. Inner and outer tether positions correspond to 15-nt and 30-nt ssDNA tethers, respectively. **c** caDNAno design of the stator top and bottom layers, highlighting tether, biotin, and fluorescent-marker staples. **d** caDNAno design of the $50\,\mathrm{nm}$-long proximal six-helix-bundle rotor segment.

### 2 Inference of restraint potentials

#### 2.1 Identification of calibration rotors and bias strength

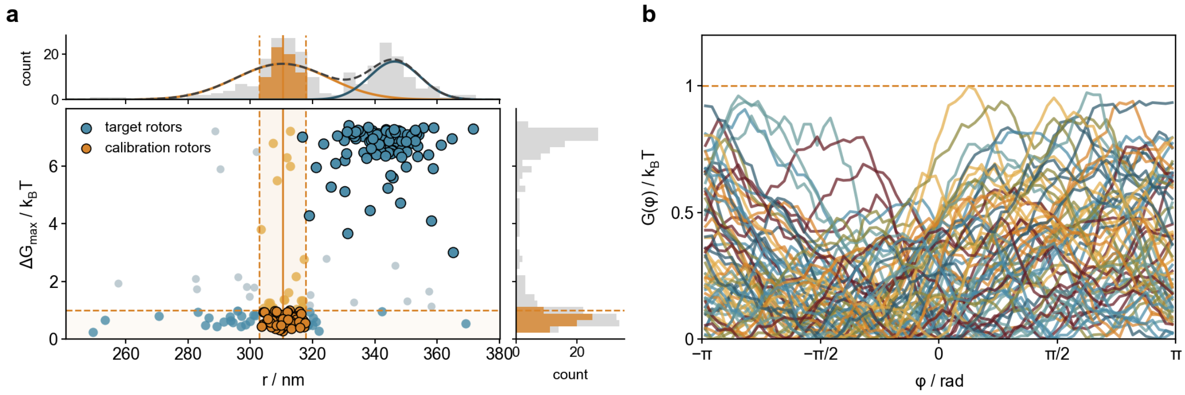

**Figure S2: Identification of calibration rotors.** **a** Rotor classification based on radius $r$ and their free-energy range $\Delta G_{\max}$ obtained from diffusive trajectories during 60 s of recording. Calibration rotors are identified by their smaller radius $r_c < r_t$ and their flat energy landscape $< 1\ k_{\mathrm{B}}t$. **b** Free-energy landscapes $G(\varphi)$ of identified calibration rotors.

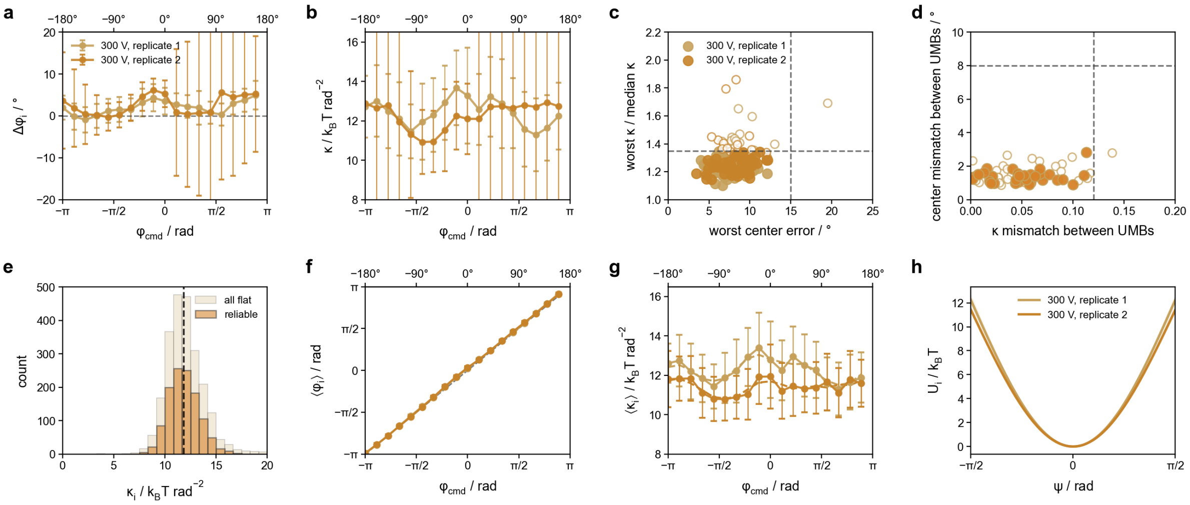

**Figure S3: Selection and characterization of calibration rotors for bias inference.** **a,b** Window-wise deviation of the inferred bias center from the commanded field angle, $\Delta\varphi_i$, and bias strength, $\kappa_i$, for two replicate measurements at $300\,\mathrm{V}$. **c** Calibration-rotor selection based on center error and variation in bias strength across umbrella windows. **d** Reproducibility of bias center and strength across replicate measurements. **e** Distribution of $\kappa_i$ before and after filtering for reliable calibration rotors; dashed line indicates the ensemble median. **f,g** Ensemble bias centers and strengths across umbrella windows after filtering. **h** Resulting ensemble bias potentials for the two replicate measurements. Dashed lines in **c,d** indicate the applied selection thresholds.

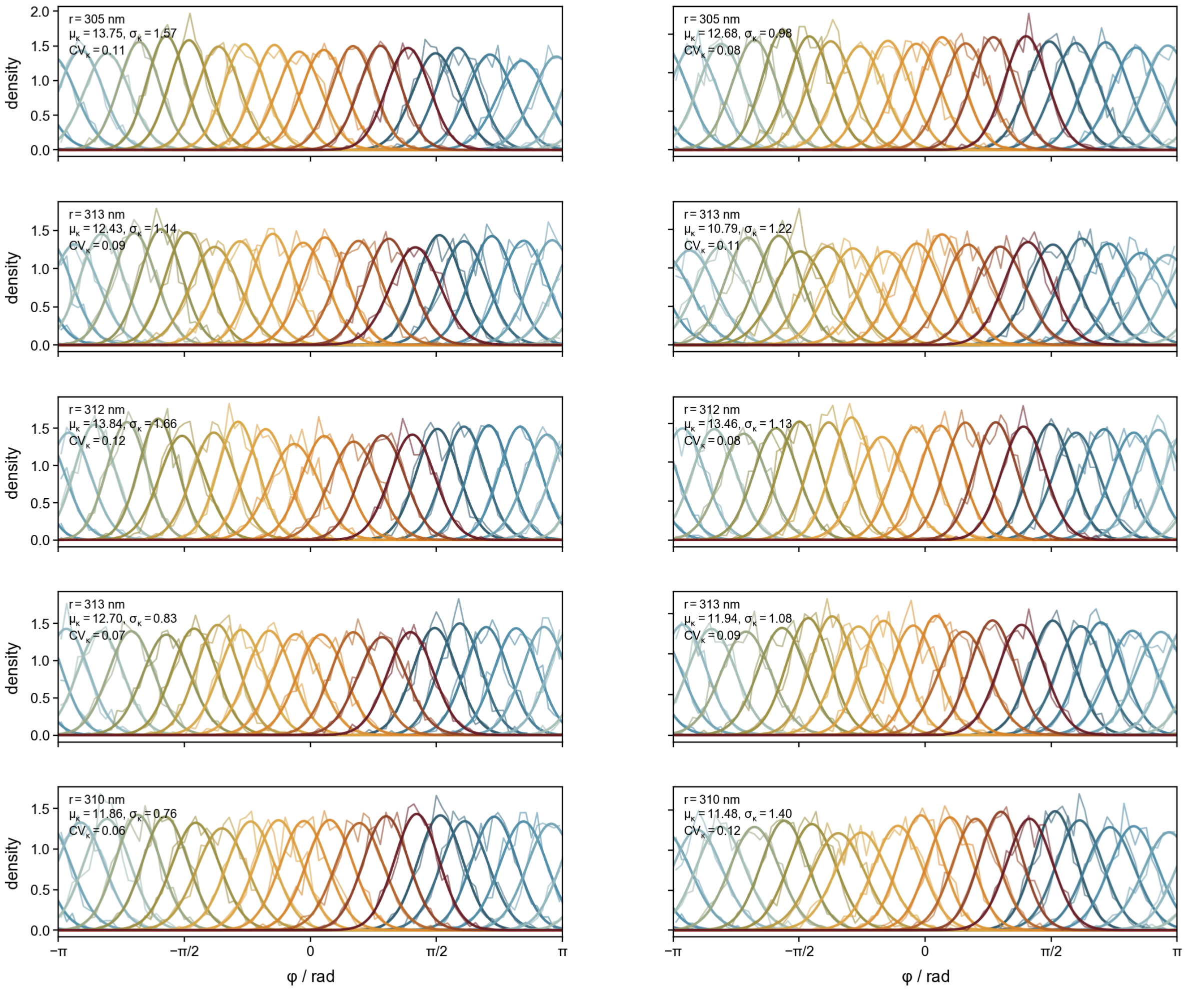

**Figure S4: Angular distributions of exemplary calibration rotors across umbrella windows.** Angular probability densities for five calibration rotors from each of two replicate measurements at $300\,\mathrm{V}$. Thin lines show the measured angular distributions and thick lines the corresponding fitted distributions for the individual umbrella windows. Colors indicate the commanded field orientations. Insets report the rotor radius $r$, mean bias strength $\mu_\kappa$, its standard deviation $\sigma_\kappa$, and coefficient of variation $\mathrm{CV}_\kappa$ across windows.

#### 2.2 Rescaling the bias strength

The electric-field-induced bias was determined *in situ* from calibration rotors with an approximately flat intrinsic free-energy landscape. Their angular distributions therefore directly report the realized bias center $\varphi_i$ and strength $\kappa_i$ in each umbrella window. Because calibration and target rotors differ in their pivot positions, their bias strengths were corrected for the resulting difference in electric-field-induced torque. Assuming a uniform effective line charge density $\lambda$, the maximum torque magnitude generated by an electric field acting on an arm segment of length $x_i$ is

$$\tau_{i,\max} = \int_0^{x_i} x\lambda E \,\mathrm{d}x = \frac{\lambda E}{2} x_i^2.$$

For a rotor with segment lengths $x_1$ and $x_2$ on either side of the pivot, the opposing contributions give

$$\tau_{\max} = \frac{\lambda E}{2} \left(x_1^2 - x_2^2\right).$$

The experimentally inferred angular bias is described by the cosine potential

$$U_i(\varphi) = \kappa_i \left[1 - \cos\left(\varphi - \varphi_i\right)\right].$$

The corresponding torque follows from $\tau = -\partial U / \partial \varphi$, such that the maximum torque magnitude is

$$\tau_i^{\max} = \kappa_i.$$

Thus, the bias strength for the six-helix-bundle rotors scales with the geometrical torque factor $\kappa \propto x_1^2 - x_2^2$. For the calibration and target rotors, the respective arm lengths $(x_1, x_2)$ are $(375, 40)$ nm and $(440, 25)$ nm, yielding

$$\kappa_\mathrm{t} = \frac{440^2 - 25^2}{375^2 - 40^2} \kappa_\mathrm{c} \approx 1.388\,\kappa_\mathrm{c}$$

Accordingly, the calibration-derived bias strengths were multiplied by 1.39 for reconstruction of the target-rotor landscapes.

### 3 Reconstruction of free-energy landscapes

#### 3.1 Statistical reweighting of biased angular distributions using MBAR

In umbrella window $i$ ($i = 1, \ldots, m$), the rotor experiences the intrinsic free-energy landscape $G(\varphi)$ and an external bias potential $U_i(\varphi)$. The corresponding equilibrium probability density is

$$p_i(\varphi) = \frac{1}{Z_i} \exp\left[-\beta\left(G(\varphi) + U_i(\varphi)\right)\right],$$

where $\beta = 1/k_B T$ and $Z_i$ is the partition function of umbrella window $i$, defined as

$$Z_i = \int_0^{2\pi} \exp\left[-\beta\left(G(\varphi) + U_i(\varphi)\right)\right] \, d\varphi.$$

Rearranging the biased equilibrium distribution gives $G(\varphi) = -k_B T \ln p_i(\varphi) - U_i(\varphi) - k_B T \ln Z_i$. The intrinsic free-energy landscape can therefore be recovered by accounting for the applied bias potential and the corresponding partition function. Whereas the bias potentials are determined from the calibration rotors and described by $U_i(\varphi) = \kappa_i[1 - \cos(\varphi - \varphi_i)]$, the partition functions $Z_i$ of the individual umbrella windows are also unknown.

To determine their relative values $Z_j/Z_i$, we exploit the fact that the same rotor angle $\varphi$ can be evaluated under the bias potentials of different umbrella windows. For windows $i$ and $j$, the difference between their effective free-energy landscapes is $[G(\varphi) + U_j(\varphi)] - [G(\varphi) + U_i(\varphi)] = U_j(\varphi) - U_i(\varphi)$, since the unknown intrinsic landscape cancels. Consequently, their equilibrium distributions are related by

$$p_i(\varphi) e^{-\beta[U_j(\varphi) - U_i(\varphi)]} = \frac{Z_j}{Z_i} p_j(\varphi).$$

This relationship allows angular observations from one umbrella window to be statistically reweighted to another, providing access to their relative partition functions.

Following Bennett's approach for estimating free-energy differences from observations of different equilibrium states [4], we employ the Multistate Bennett Acceptance Ratio (MBAR) method [5] similar to the python package pymbar [6], which combines observations from all umbrella windows to determine their relative partition functions self-consistently. Let $N_i$ denote the number of angular observations recorded in umbrella window $i$, and $N = \sum_{i=1}^{m} N_i$ be the total number of observations. The combined angular observations from all umbrella windows can be described by the mixture distribution

$$p_{\mathrm{mix}}(\varphi) = \sum_{i=1}^{m} \frac{N_i}{N} p_i(\varphi),$$

where each umbrella window contributes to the mixture in proportion to its total number of angular observations. Substituting $p_i(\varphi)$ gives

$$p_{\mathrm{mix}}(\varphi) = \frac{1}{N} \sum_{i=1}^{m} N_i Z_i^{-1} e^{-\beta[G(\varphi) + U_i(\varphi)]}.$$

We can now express the partition function $Z_i$ in terms of the pooled distribution $p_{\mathrm{mix}}(\varphi)$ by extending and rewriting its defining integral as

$$\begin{aligned} Z_i &= \int_0^{2\pi} p_{\mathrm{mix}}(\varphi) \, \frac{e^{-\beta[G(\varphi) + U_i(\varphi)]}}{p_{\mathrm{mix}}(\varphi)} \, d\varphi \\ &= \int_0^{2\pi} p_{\mathrm{mix}}(\varphi) \, h_i(\varphi) \, d\varphi \end{aligned}$$

where we have introduced the function $h_i(\varphi) = e^{-\beta[G(\varphi) + U_i(\varphi)]} / p_{\mathrm{mix}}(\varphi)$. The partition function $Z_i$ can thus be expressed as the expectation value of $h_i(\varphi)$ over the pooled distribution $p_{\mathrm{mix}}(\varphi)$. This expectation value can be approximated using the angular observations concatenated from all umbrella windows

$$Z_i \approx \frac{1}{N}\sum_{n=1}^{N} h_i(\varphi_n).$$

Substituting the definition of $h_i(\varphi)$ gives

$$Z_i \approx \frac{1}{N}\sum_{n=1}^{N} \frac{e^{-\beta[G(\varphi_n)+U_i(\varphi_n)]}}{p_{\text{mix}}(\varphi_n)}.$$

Inserting the expression for the mixture distribution $p_{\text{mix}}(\varphi_n)$ into the denominator, the factors $1/N$ cancel, yielding

$$Z_i \approx \sum_{n=1}^{N} \frac{e^{-\beta[G(\varphi_n)+U_i(\varphi_n)]}}{\sum_{j=1}^{m} N_j Z_j^{-1} e^{-\beta[G(\varphi_n)+U_j(\varphi_n)]}}.$$

Since the free-energy landscape $G(\varphi)$ is identical for every umbrella window, the factor $e^{-\beta G(\varphi_n)}$ cancels, giving the MBAR self-consistency equations

$$Z_i \approx \sum_{n=1}^{N} \frac{e^{-\beta U_i(\varphi_n)}}{\sum_{j=1}^{m} N_j Z_j^{-1} e^{-\beta U_j(\varphi_n)}}.$$

Although the intrinsic free-energy landscape $G(\varphi)$ is unknown, the partition functions of the biased equilibrium states can be determined self-consistently using only the measured angular positions and the known umbrella potentials. However, their absolute values remain undetermined, since multiplying all $Z_i$ by the same positive constant leaves the equations unchanged, which reflects the arbitrary additive constant of $G(\varphi)$. MBAR therefore determines only the ratios of partition functions, which correspond to free-energy differences between the biased equilibrium states,

$$F_i - F_j = -k_B T \ln\left(\frac{Z_i}{Z_j}\right),$$

where $F_i = -k_B T \ln Z_i$ denotes the free energy associated with biased equilibrium state $i$. The coupled MBAR equations were solved using damped fixed-point iteration in terms of the dimensionless free energies $f_i = -\ln Z_i$, with the reference fixed to $f_1 = 0$. Iterations were continued until the maximum absolute change in $f_i$ between successive iterations was below $10^{-7}$, or until a maximum of 1000 iterations was reached.

Having determined the relative partition functions $Z_i$, we can reconstruct $G(\varphi)$ from the pooled observations by accounting for the applied bias potentials and their relative partition functions. Using the definition of the pooled distribution, we obtain

$$N p_{\text{mix}}(\varphi) = e^{-\beta G(\varphi)} \sum_{i=1}^{m} N_i Z_i^{-1} e^{-\beta U_i(\varphi)}.$$

Rearranging and taking the logarithm yields the intrinsic free-energy landscape

$$G(\varphi) = -k_B T \ln\left(\frac{N p_{\text{mix}}(\varphi)}{\sum_{i=1}^{m} N_i Z_i^{-1} e^{-\beta U_i(\varphi)}}\right).$$

Thus, the unbiased angular probability density, $p_0(\varphi) \propto e^{-\beta G(\varphi)}$, is reconstructed by assigning each recorded datapoint $\varphi_n$ a statistical weight

$$w_n = \left(\sum_{i=1}^{m} N_i Z_i^{-1} e^{-\beta U_i(\varphi_n)}\right)^{-1}.$$

These weights are normalized such that their sum over all observations is 1. The angular range is then divided into 72 equally spaced bins, and the normalized weights of all observations within each bin are summed and divided by the bin width to obtain $p_0(\varphi)$. The free energy landscape is then calculated from Boltzmann inversion of $p_0$.

#### 3.2 Reconstruction gallery

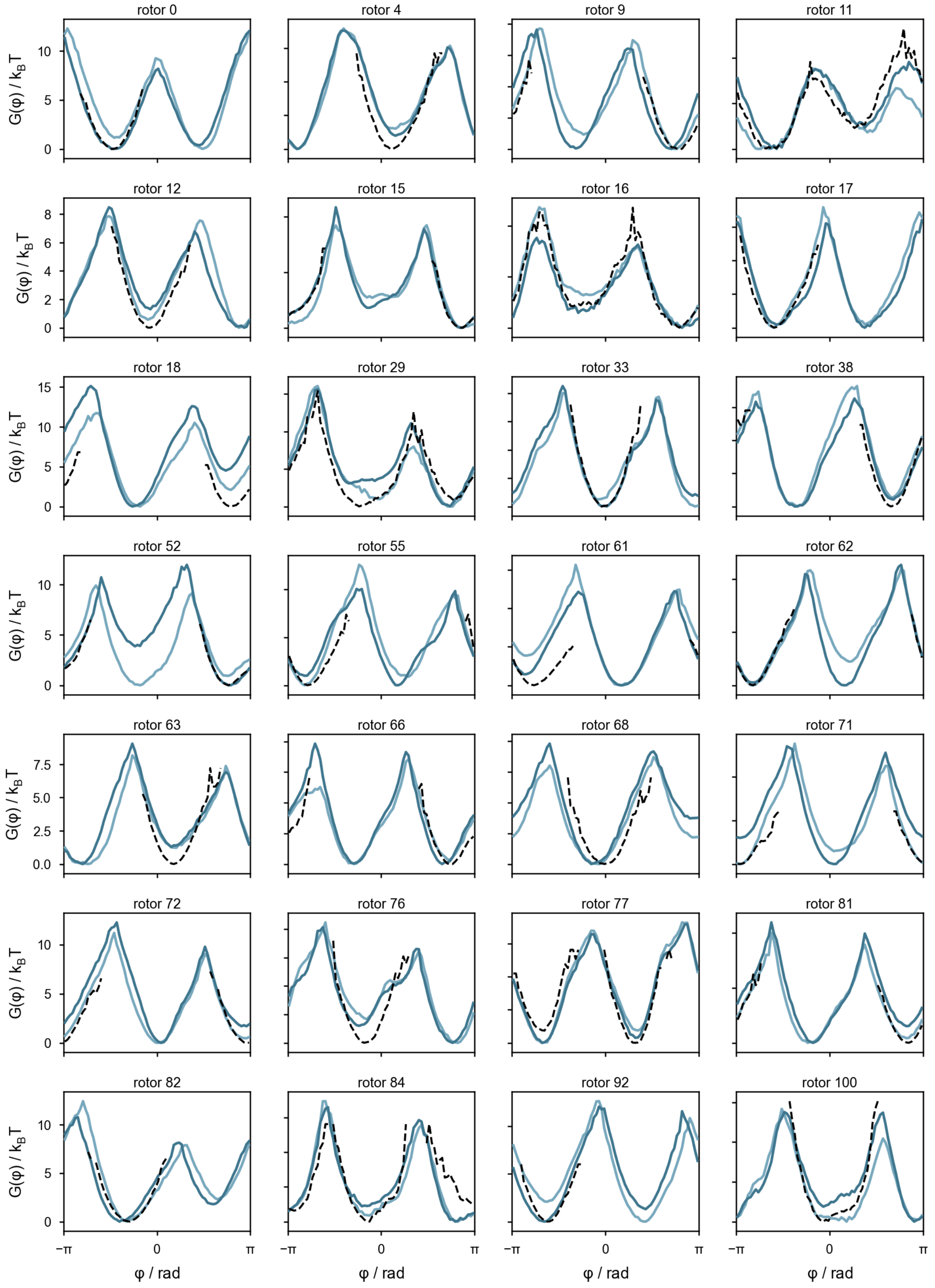

**Figure S5: Reconstructed free-energy landscapes for exemplary target rotors.** Free-energy landscapes $G(\varphi)$ reconstructed from two independent umbrella-sampling measurements of the same target rotors (blue), compared with the corresponding landscape obtained from 60 s of passive diffusion (black dashed line).

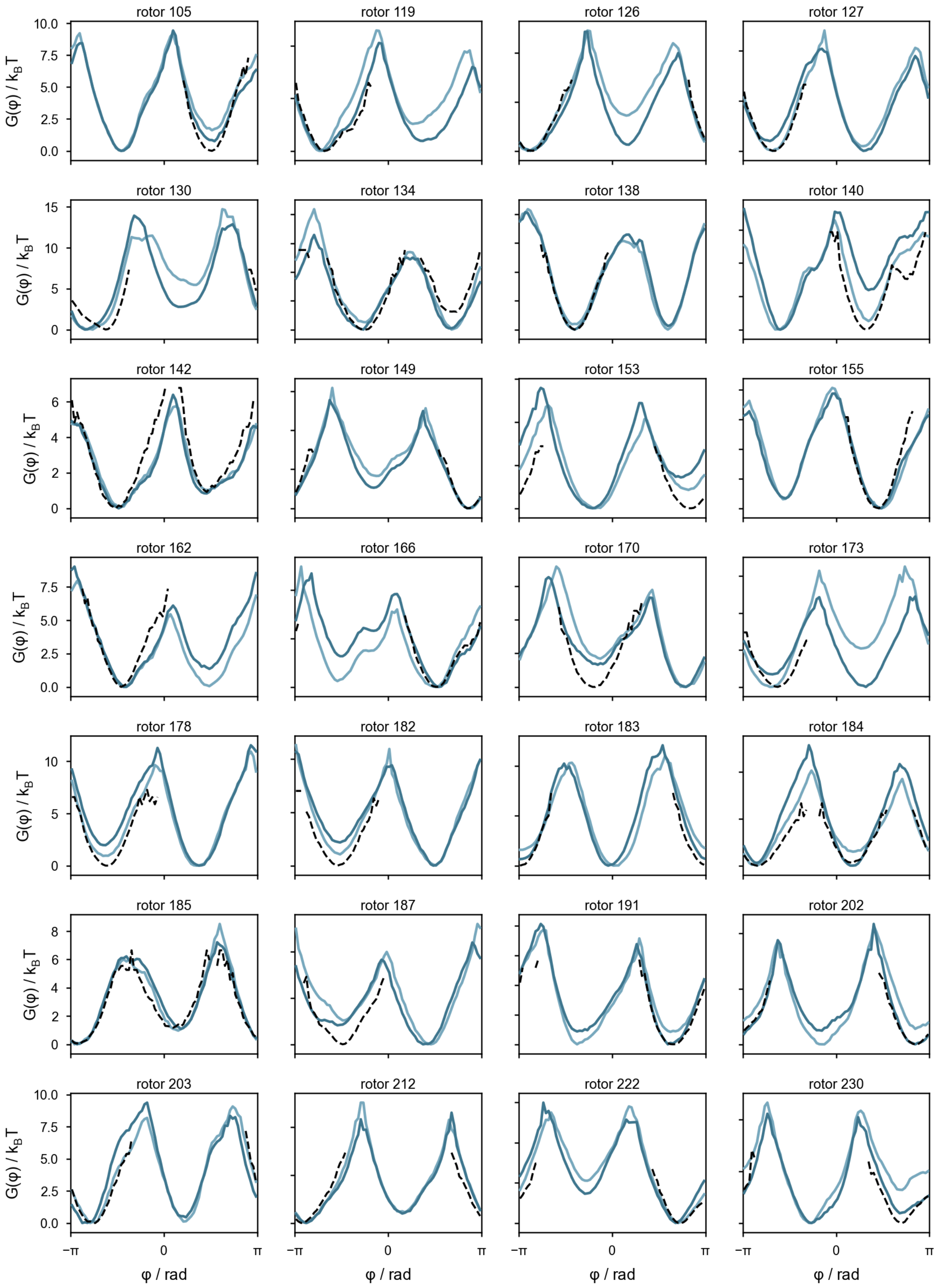

**Figure S6: Reconstruction gallery of individual target rotors (continued).**

#### 3.3 Run-to-run reproducibility of free-energy reconstruction

We quantified the run-to-run reproducibility by comparing independently reconstructed free-energy landscapes of the same rotor using the root-mean-square error (RMSE), the Pearson correlation coefficient $\rho_{1,2}$ and the difference in barrier height. The RMSE measures the absolute difference in free energy between the two landscapes across all valid angular bins $N \leq 72$,

$$\mathrm{run-to-runRMSE} = \sqrt{\frac{1}{N}\sum_{i=1}^{N}\left[G_1(\varphi_i) - G_2(\varphi_i)\right]^2}, \tag{S1}$$

whereas $\rho$ quantifies the similarity in shape of the two landscapes, independently of their absolute energy scale and offset. Both metrics were calculated before and after angular alignment of the landscapes, allowing relative shifts of up to $\pm 15°$.

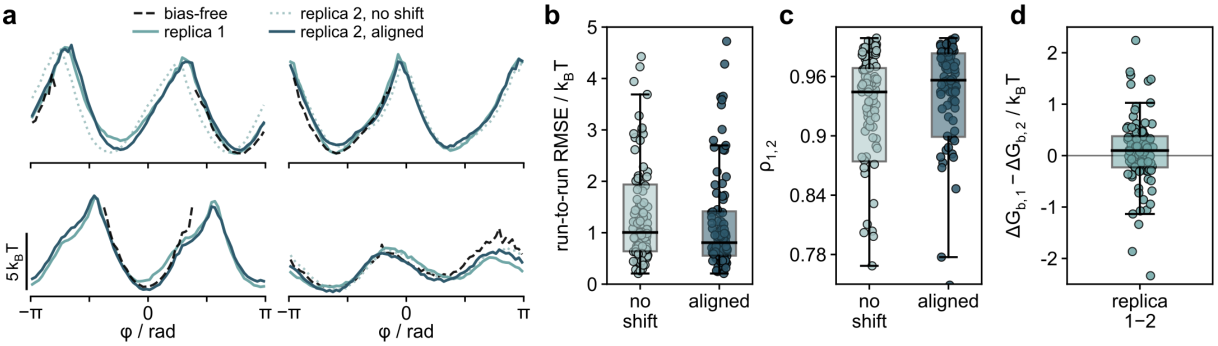

**Figure S7: Reproducibility of free-energy reconstruction across replicate measurements.** **a** Exemplary free-energy landscapes reconstructed from two independent umbrella-sampling measurements before and after alignment, compared with the corresponding bias-free landscapes. **b** Rotor-wise RMSE between replicate reconstructions before and after alignment. **c** Rotor-wise correlation between replicate landscapes before and after alignment. **d** Difference in reconstructed barrier height, $\Delta G_\mathrm{b}$, between replicate measurements.

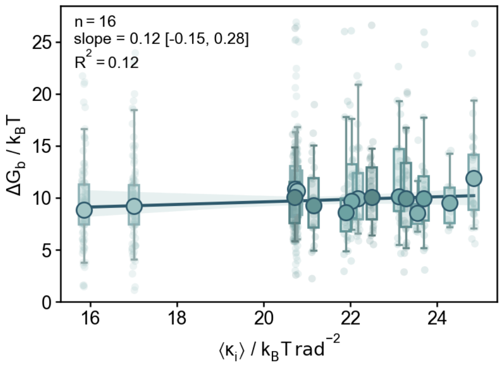

**Figure S8: Independence of reconstruction from inferred bias strength.** Mean reconstructed barrier height, $\Delta G_\mathrm{b}$, as a function of the mean bias strength, $\langle \kappa_i \rangle$, across 14 measurements. Small points and box plots show the distributions of individual target rotors, while large circles indicate the measurement-wise means. The linear fit (95% confidence level) shows no systematic dependence of the reconstructed barrier height on bias strength.

#### 3.4 Statistical uncertainty of free-energy reconstruction

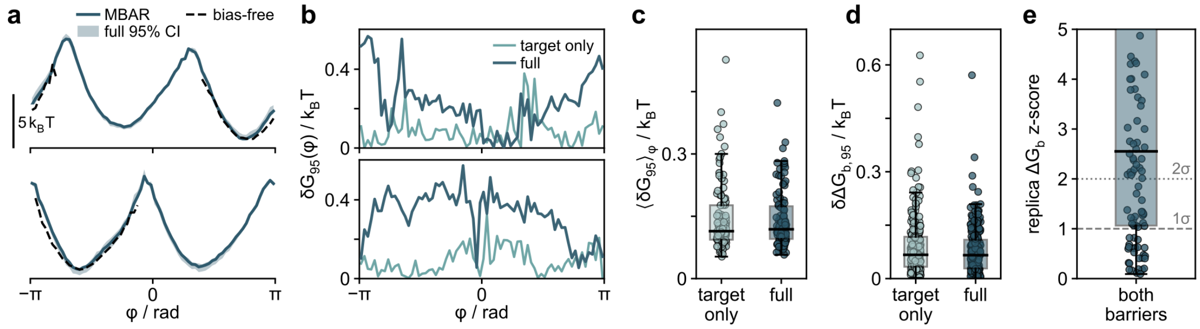

**Figure S9: Statistical uncertainty of the reconstructed free-energy landscapes.** **a** Exemplary MBAR-reconstructed free-energy landscapes with 95% confidence intervals (CI) obtained by block bootstrapping, compared with the corresponding bias-free landscapes. **b** Angular dependence of the 95% confidence-interval half-width, $\delta G_{95}(\varphi)$, for the same exemplary rotors, obtained by resampling either only the target-rotor trajectories (target only) or both target- and calibration-rotor trajectories with re-inference of the bias potentials (full). **c** Distribution of the mean 95% confidence-interval half-width, $\langle \delta G_{95} \rangle_\varphi$, obtained by averaging over the angular coordinate for each rotor. Points represent individual rotors and box plots summarize the rotor ensemble. **d** Distribution of the 95% confidence-interval half-width of the reconstructed barrier heights, $\delta \Delta G_{\mathrm{b},95}$, for target-only and full bootstrapping. Points represent individual barriers and box plots summarize the rotor ensemble. **e** Run-to-run consistency of the reconstructed barrier heights, expressed as the $z$-score defined in the text. Both barriers of each rotor are included. Points represent individual barriers and the box plot summarizes their distribution; dashed and dotted lines indicate $z = 1$ and $z = 2$, respectively.

Statistical uncertainty was estimated by block bootstrapping the angular trajectories within each umbrella window using $250\,\mathrm{ms}$ blocks. For each bootstrap realization, the resampled trajectories were used to reconstruct a new free-energy landscape, and the 95% confidence interval was obtained from the resulting distribution of $G(\varphi)$. Figure S9a shows the reconstructed landscapes and confidence intervals for two exemplary rotors, while Fig. S9b shows the corresponding confidence-interval width, $\delta G_{95}(\varphi)$, along the angular coordinate.

To assess the contribution of the experimentally inferred bias to the statistical uncertainty, we compared two bootstrap procedures. In the target-only bootstrap, only the target-rotor trajectories were resampled while the inferred bias potentials were kept fixed. In the full bootstrap, both target- and calibration-rotor trajectories were resampled, and the bias center and strength were re-inferred before each MBAR reconstruction. For each rotor, $\delta G_{95}(\varphi)$ was averaged over the full angular range, and the resulting distributions of $\langle \delta G_{95} \rangle_\varphi$ across the rotor ensemble are shown in Fig. S9c. The resulting rotor-wise uncertainties were similar for target-only and full bootstrapping, indicating only a moderate additional contribution from uncertainty in the inferred bias potentials.

Because the principal gain of umbrella sampling lies in reconstructing the high-energy barrier regions that are poorly sampled during passive diffusion, we next examined the uncertainty specifically at the barriers. The 95% confidence-interval half-width of the reconstructed barrier heights, $\delta \Delta G_{\mathrm{b},95}$, remained small for both target-only and full bootstrapping (Fig. S9d).

Finally, we tested whether the bootstrap-derived barrier uncertainty was consistent with the experimental run-to-run variability. For each rotor and each of the two barriers, we calculated the $z$-score

$$\Delta G_{\mathrm{b}}\,\mathrm{z-score} = \frac{\left|\Delta G_{\mathrm{b}}^{(1)} - \Delta G_{\mathrm{b}}^{(2)}\right|}{\sqrt{\sigma_{\mathrm{b},1}^2 + \sigma_{\mathrm{b},2}^2}},$$

where $\sigma_{\mathrm{b},1}$ and $\sigma_{\mathrm{b},2}$ are the bootstrap standard deviations of the barrier heights in the two independent measurements (Fig. S9e). The $z$-score distribution had a median above $2$, indicating that the differences in barrier height between independent measurements generally exceeded the statistical uncertainty estimated from within-run block bootstrapping. Thus, while the bootstrap analysis indicates small statistical uncertainties within individual reconstructions, it does not fully capture the experimental run-to-run variability, suggesting additional sources of variation between independent runs.

### 4 Conditions for reliable experimental umbrella sampling

#### 4.1 Stationarity and connectivity of umbrella windows

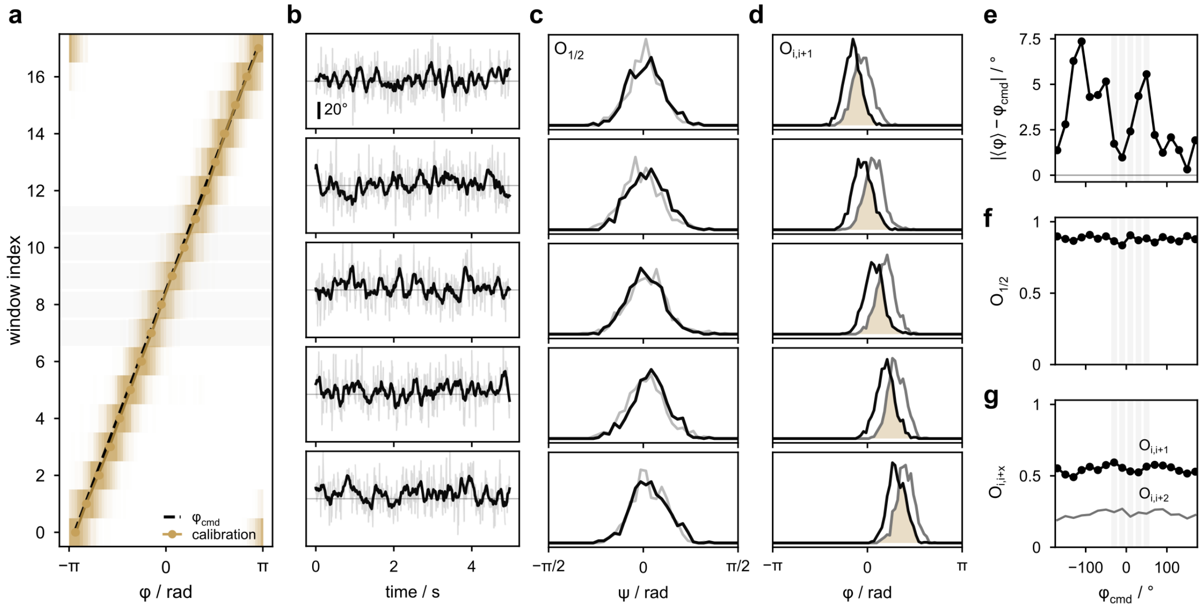

**Figure S10: Stationarity and statistical connectivity for an exemplary calibration rotor.** **a** Angular distributions of the calibration rotor across the 18 umbrella windows. The dashed line indicates the commanded field orientation, $\varphi_{\mathrm{cmd}}$, and points indicate the mean angular position in each window. **b** Angular trajectories of the calibration rotor within the highlighted $5\,\mathrm{s}$ umbrella windows in **a**. **c** Within-window overlap, $O_{1/2}$, between the angular distributions sampled during the first and second halves of five exemplary umbrella windows. Black and grey lines show the corresponding probability distributions. **d** Overlap, $O_{i,i+1}$, between the angular distributions of neighboring umbrella windows for the same examples. Shaded regions indicate the overlap between the distributions. **e** Mean steering deviation of the calibration rotor $\langle\varphi\rangle - \varphi_{\mathrm{cmd}}$, as a function of commanded field orientation. **f** Within-window overlap, $O_{1/2}$, between the first and second halves of each umbrella window. **g** Overlap between neighboring, $O_{i,i+1}$, and next-nearest-neighbor, $O_{i,i+2}$, umbrella windows as a function of commanded field orientation.

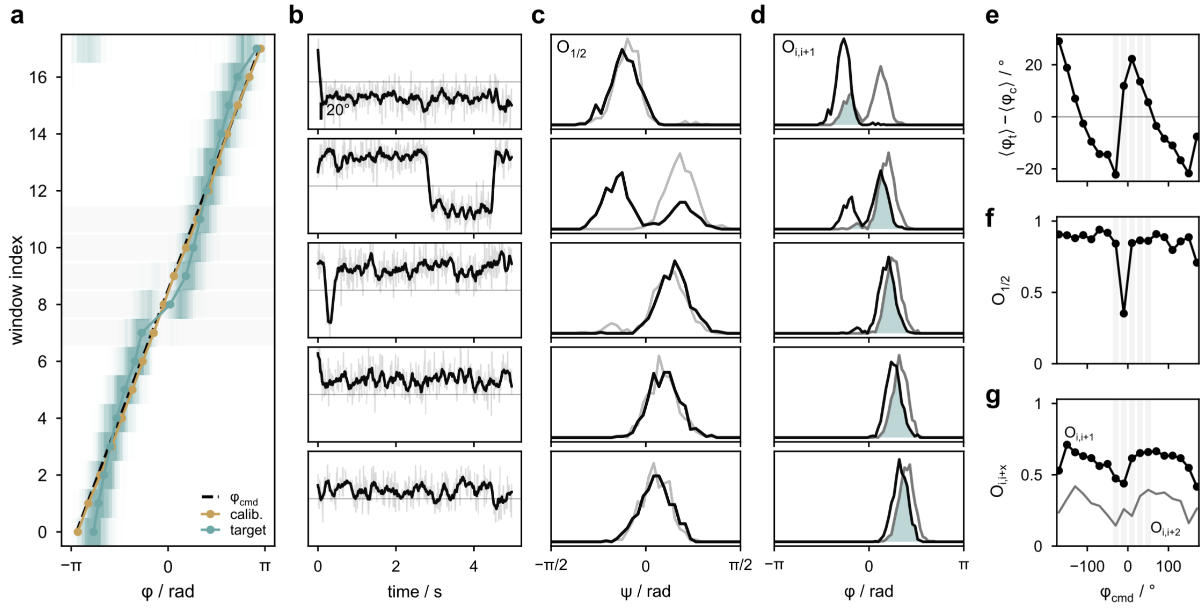

**Figure S11: Stationarity and statistical connectivity for an exemplary target rotor.** **a–d** Angular sampling, trajectories, within-window overlap, and neighboring-window overlap, respectively, as defined for the calibration rotor in Fig. S10. **e** Mean steering deviation of the target rotor from the calibration-rotor ensemble, $\langle\varphi_t\rangle - \langle\varphi_c\rangle$, as a function of commanded field orientation. **f** Within-window overlap, $O_{1/2}$. **g** Overlap between neighboring, $O_{i,i+1}$, and next-nearest-neighbor, $O_{i,i+2}$, umbrella windows.

#### 4.2 Convergence of calibration metrics and reconstruction results

To assess whether the $5\,\mathrm{s}$ acquisition time per umbrella window was sufficient, we examined convergence of calibration rotor and target rotor metrics as well as reconstruction results as progressively longer portions of each window were included.

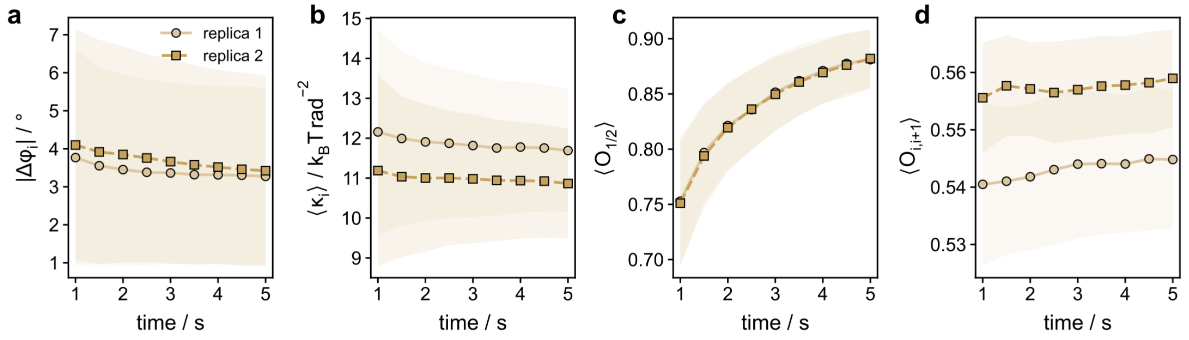

**Figure S12: Convergence of calibration-rotor metrics with increasing sampling time. a** Mean deviation of the inferred bias center from the commanded field orientation for two replicate measurements. **b** Mean inferred bias strength, $\langle\kappa\rangle$, for the two replicate measurements. **c** Mean within-window overlap, $\langle O_{1/2}\rangle$, between angular distributions sampled during the first and second halves of each umbrella window. **d** Mean overlap between neighboring umbrella windows, $\langle O_{i,i+1}\rangle$. Points and shaded regions show the ensemble mean ± standard deviation across calibration rotors.

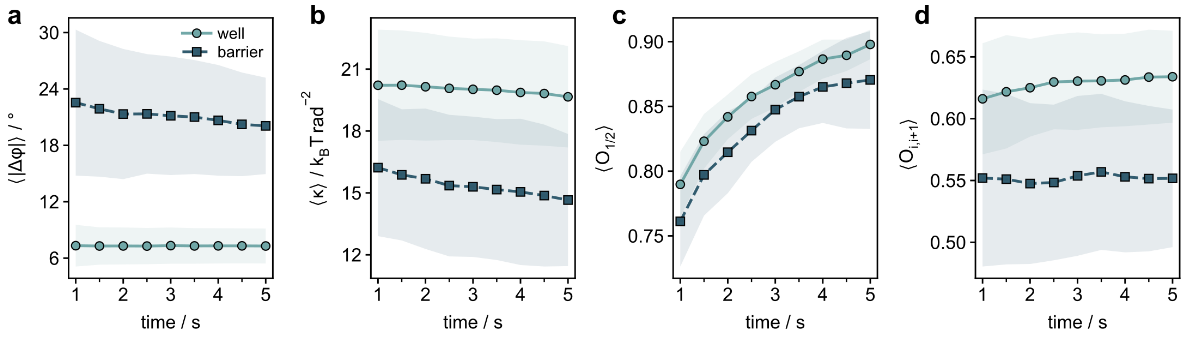

**Figure S13: Convergence of target-rotor metrics with increasing sampling time. a,b** Mean angular deviation, $\langle|\Delta\varphi|\rangle$, and effective confinement strength, $\langle\kappa\rangle$, for umbrella windows sampling the energy well and barrier. **c,d** Mean within-window overlap, $\langle O_{1/2}\rangle$, and neighboring-window overlap, $\langle O_{i,i+1}\rangle$, respectively. Points and shaded regions show the ensemble mean ± standard deviation across target rotors.

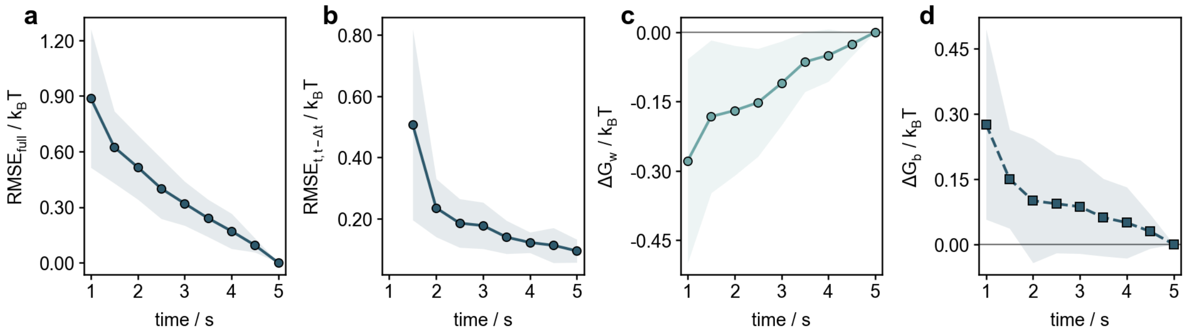

**Figure S14: Convergence of reconstructed free-energy landscapes with increasing sampling time. a** RMSE of the free-energy landscape reconstructed at each sampling time relative to the reference reconstruction obtained using the complete $5\,\mathrm{s}$ acquisition time of each umbrella window. **b** RMSE between free-energy landscapes reconstructed from successive sampling durations, $t$ and $t-\Delta t$. **c** Difference in depth of the energy minima, $\Delta G_{\mathrm{w}}$, relative to the value obtained from the complete $5\,\mathrm{s}$ windows. **d** Difference in barrier height, $\Delta G_{\mathrm{b}}$, relative to the value obtained from the complete $5\,\mathrm{s}$ windows. Points and shaded regions show the ensemble mean ± standard deviation across target rotors.

#### 4.3 Temporal correlation and effective sampling size

Because successive angular measurements within an umbrella window are temporally correlated, the number of recorded frames does not directly correspond to the number of statistically independent samples. We therefore characterized the temporal correlations of the rotor dynamics and the resulting effective sampling within individual umbrella windows. In particular, slowly decaying correlations (in windows sampling the barriers) reduce the effective number of independent samples and increase statistical uncertainty.

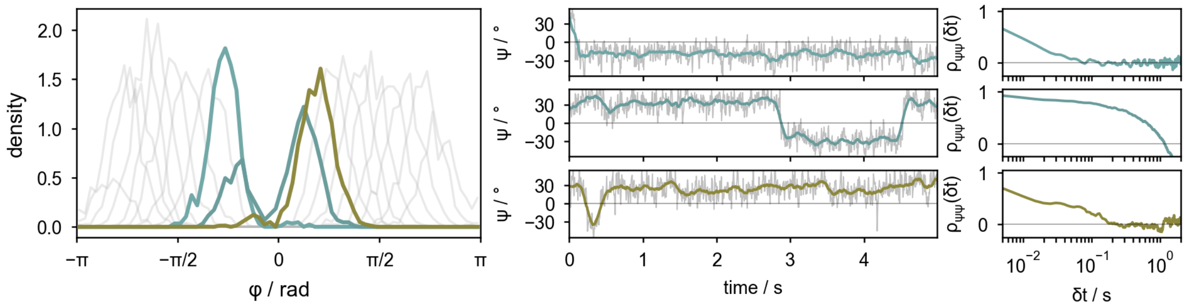

**Figure S15: Temporal correlations within umbrella windows for an exemplary target rotor.** Angular probability distributions, corresponding angular trajectories, and autocorrelation functions for three exemplary umbrella windows sampling different regions of the free-energy landscape. Grey lines in the probability-density panel show the distributions of the remaining umbrella windows. The autocorrelation functions illustrate the finite correlation time of the angular fluctuations within each biased window.

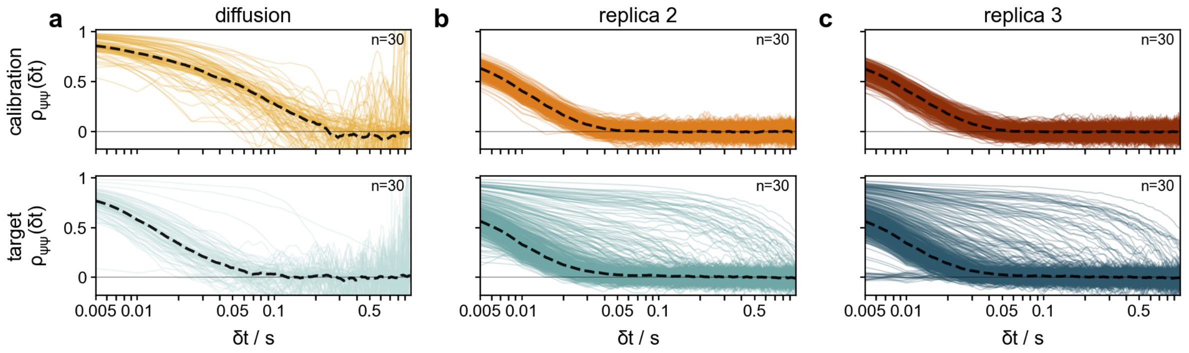

**Figure S16: Temporal autocorrelation of calibration and target rotors during passive diffusion and umbrella sampling.** **a** Angular autocorrelation functions, $\rho_{\psi\psi}(\delta t)$, during passive diffusion. **b, c** Angular autocorrelation functions during two independent umbrella-sampling measurements. Top and bottom rows show calibration and target rotors, respectively. Thin lines represent individual rotor trajectories during passive diffusion and during 18 umbrella sampling windows. Dashed lines show the corresponding ensemble-averaged autocorrelation functions. Each dataset contains $n = 30$ rotors.

#### 4.4 Sensitivity to the calibration-rotor ensemble

To assess the sensitivity of the reconstruction to the calibration-rotor ensemble, we repeated the bias inference and MBAR reconstruction using randomly selected subsets of calibration rotors.

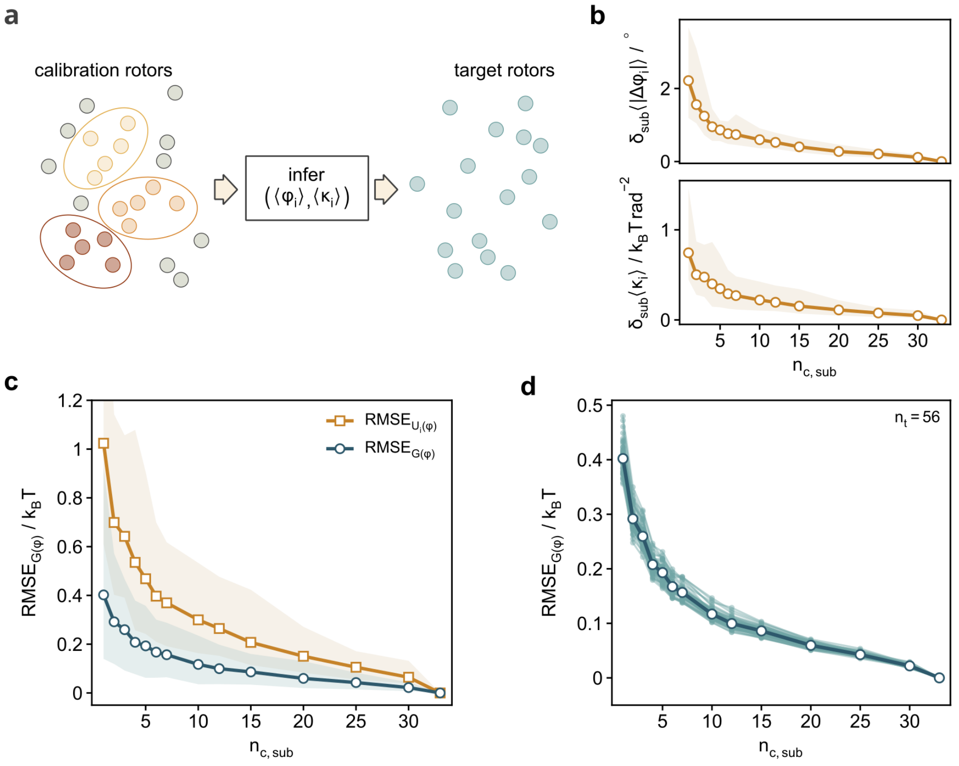

**Figure S17: Sensitivity of free-energy reconstruction to calibration-ensemble size.** **a** Schematic of the calibration-subset analysis, in which randomly selected subsets of calibration rotors were used for bias inference and subsequent reconstruction of the target-rotor landscapes. **b** Deviation of the inferred bias center and bias strength from the values obtained using the complete calibration ensemble as a function of calibration-subset size, $n_{\mathrm{c,sub}}$. **c** RMSE of the inferred bias potentials, $U_i(\varphi)$, and reconstructed free-energy landscapes, $G(\varphi)$, relative to the results obtained using the complete calibration ensemble. **d** RMSE of $G(\varphi)$ for individual target rotors relative to the complete-ensemble reconstruction. Thin lines show individual target rotors and the dark line the ensemble mean. Shaded regions in **b, c** indicate the 95% interval across randomly selected calibration subsets.

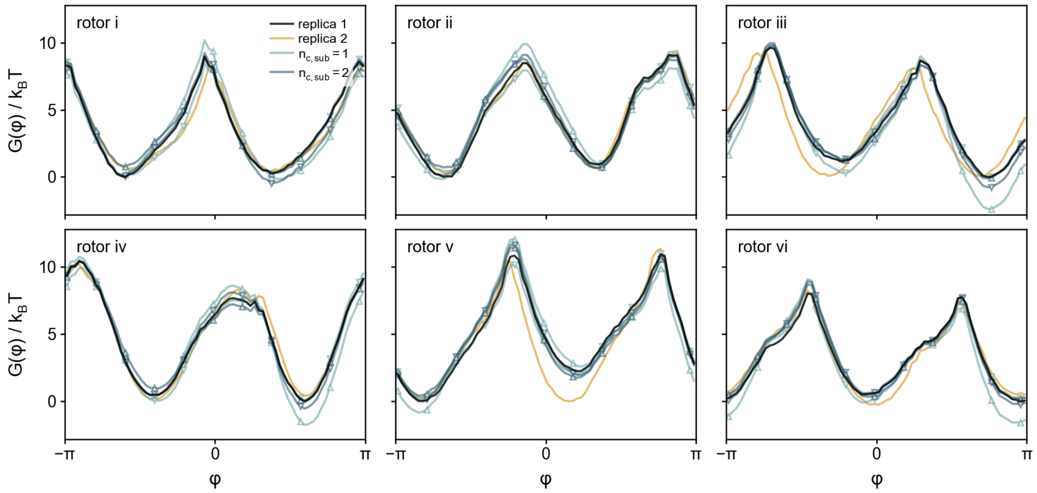

**Figure S18: Exemplary free-energy reconstructions for small calibration subsets.** Free-energy landscapes $G(\varphi)$ of six exemplary target rotors reconstructed using subsets of $n_{\mathrm{c,sub}} = 1$ and $2$ calibration rotors. Reconstructions from the two independent umbrella-sampling measurements using the complete calibration ensemble are shown for comparison.

### 5 Influence of umbrella-window spacing

#### 5.1 Connectivity between neighboring umbrella windows

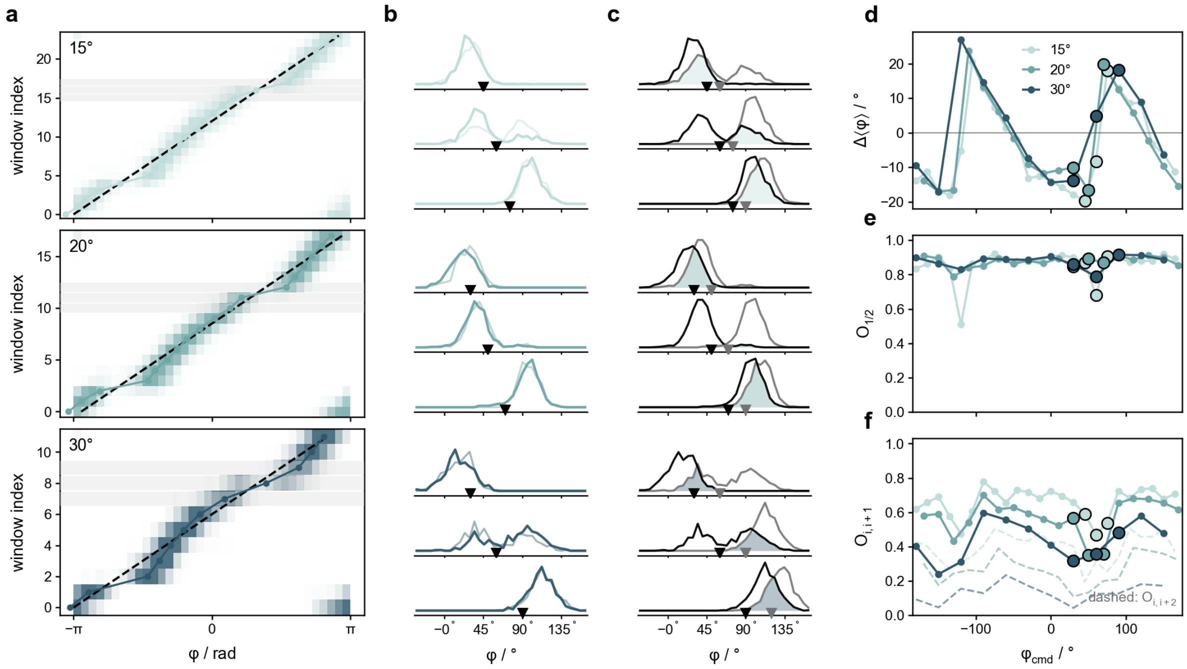

**Figure S19: Statistical connectivity across different umbrella-window spacings for an exemplary target rotor. a** Angular distributions across umbrella windows recorded with $15°$, $20°$, and $30°$ spacing. Dashed lines indicate the commanded field orientations. **b** Angular distributions for three exemplary umbrella windows at each window spacing; triangles indicate the corresponding commanded field orientations. **c** Overlap between neighboring angular distributions for the same exemplary windows. Shaded regions indicate the overlap between distributions. **d** Mean angular deviation, $\Delta\varphi$, from the commanded field orientation across umbrella windows. **e** Within-window overlap, $O_{1/2}$, between the angular distributions sampled during the first and second halves of each umbrella window. **f** Overlap between neighboring, $O_{i,i+1}$, and next-nearest-neighbor, $O_{i,i+2}$, umbrella windows. Solid and dashed lines indicate $O_{i,i+1}$ and $O_{i,i+2}$, respectively.

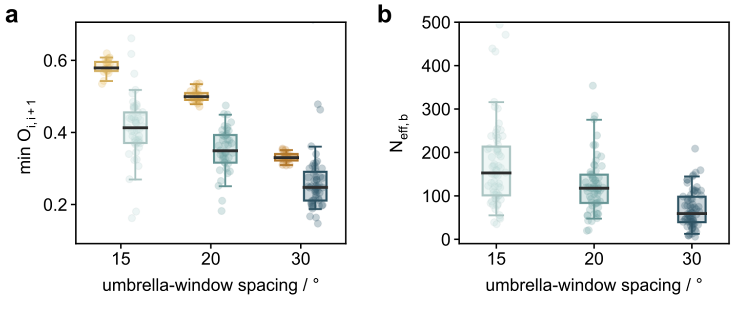

**Figure S20: Statistical connectivity across different umbrella-window spacings. a** Overlap, $O_{i,i+1}$, between neighboring retained umbrella windows for the original $15°$ spacing and the two subsampled window sets A and B. **b** Corresponding effective number of samples, $N_{\mathrm{eff,b}}$, in the barrier region. Points represent individual target rotors and box plots summarize the rotor ensemble.

#### 5.2 Trade-off between window spacing and sampling time per window

To examine the trade-off between window density and sampling time per window, we compared reconstructions obtained with $30°$, $20°$, and $15°$ window spacing, adjusting the acquisition time per window to ensure the same total acquisition time for each protocol.

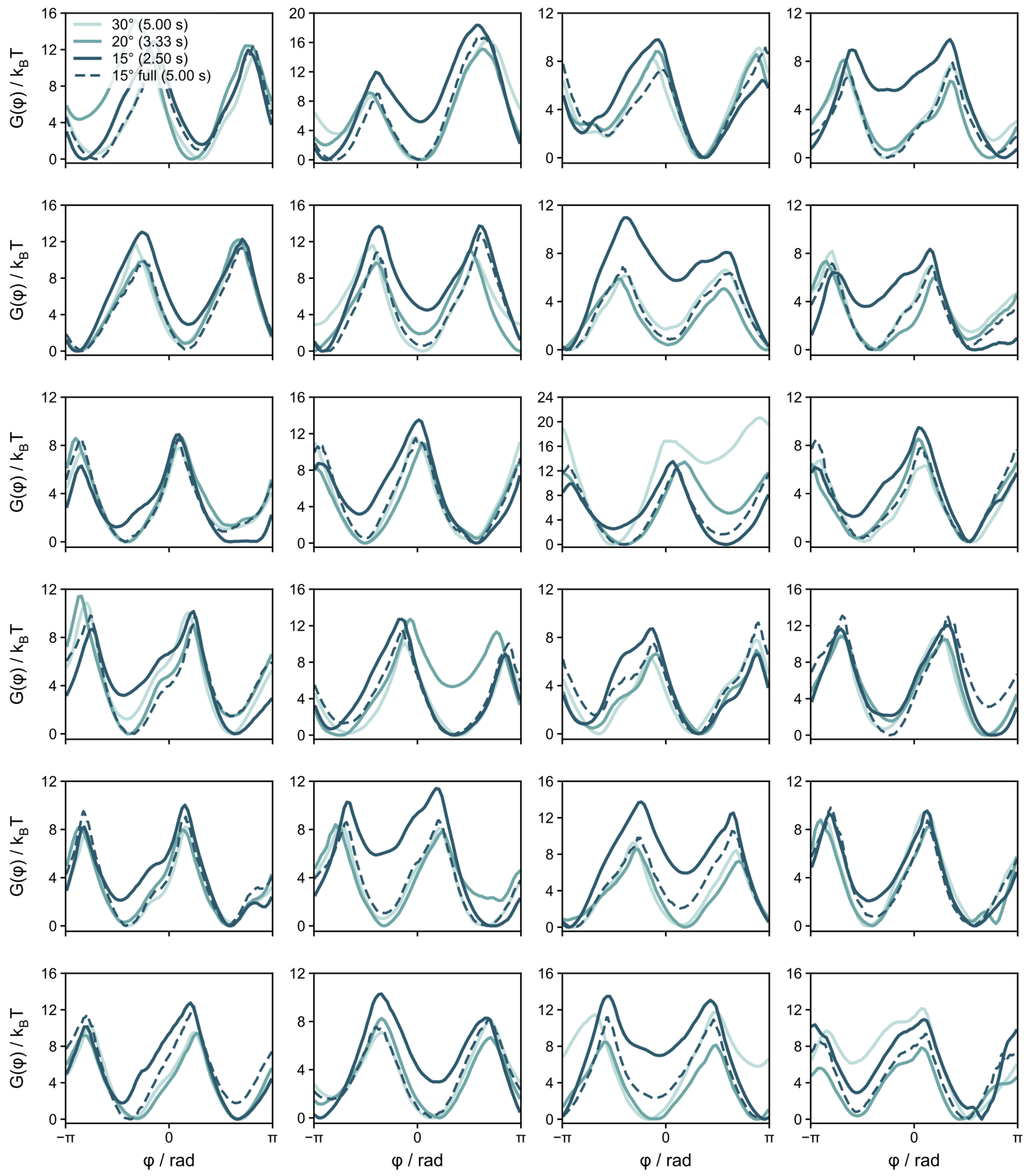

**Figure S21: Trade-off between umbrella-window spacing and sampling time per window at fixed total acquisition time.** Exemplary target-rotor free-energy landscapes reconstructed using protocols with different window spacings and corresponding acquisition times per window while maintaining the same total biased acquisition time. Reconstructions are compared with the corresponding reference landscapes obtained from passive diffusion.

#### 5.3 Artificial downsampling of densely sampled reconstructions

To isolate the effect of umbrella-window spacing from differences in sampling time per window, densely sampled $15°$-spaced recordings were artificially downsampled by retaining alternating windows, yielding two complementary $30°$-spaced subsets (A and B). The resulting sampling statistics and reconstructed free-energy landscapes were compared with those obtained from the complete $15°$ dataset. Despite the reduced overlap between neighboring windows, the reconstructed landscapes showed only moderate deviations from the complete reconstruction (Fig. S22 and S23).

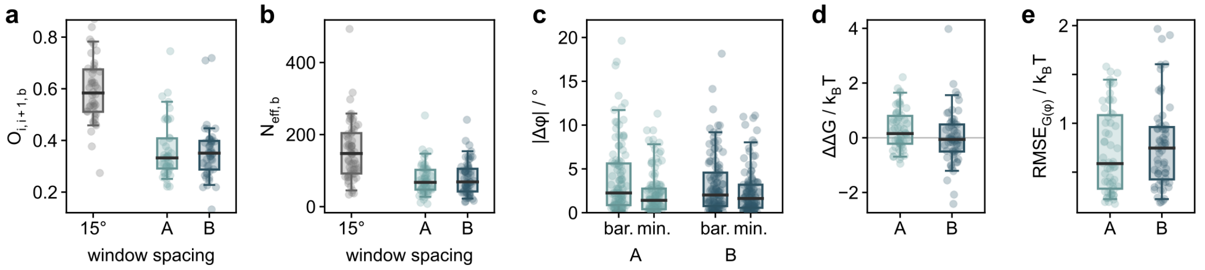

**Figure S22: Effect of artificial umbrella-window downsampling on sampling and free-energy reconstruction. a** Overlap, $O_{i,i+1}$, between neighboring retained umbrella windows for the complete $15°$-spaced recording and two complementary $30°$-spaced subsets (A and B) obtained by retaining alternating windows. **b** Effective number of samples, $N_{\mathrm{eff,b}}$, in the barrier region. **c** Angular deviation, $|\Delta\varphi|$, for windows sampling the barrier (bar.) and minimum (min.) regions in the two $30°$ subsets. **d** Change in reconstructed barrier height, $\Delta\Delta G$, relative to the complete $15°$ reconstruction. **e** RMSE of the reconstructed free-energy landscape, $G(\varphi)$, relative to the complete $15°$ reconstruction. Points represent individual target rotors and box plots summarize the rotor ensemble.

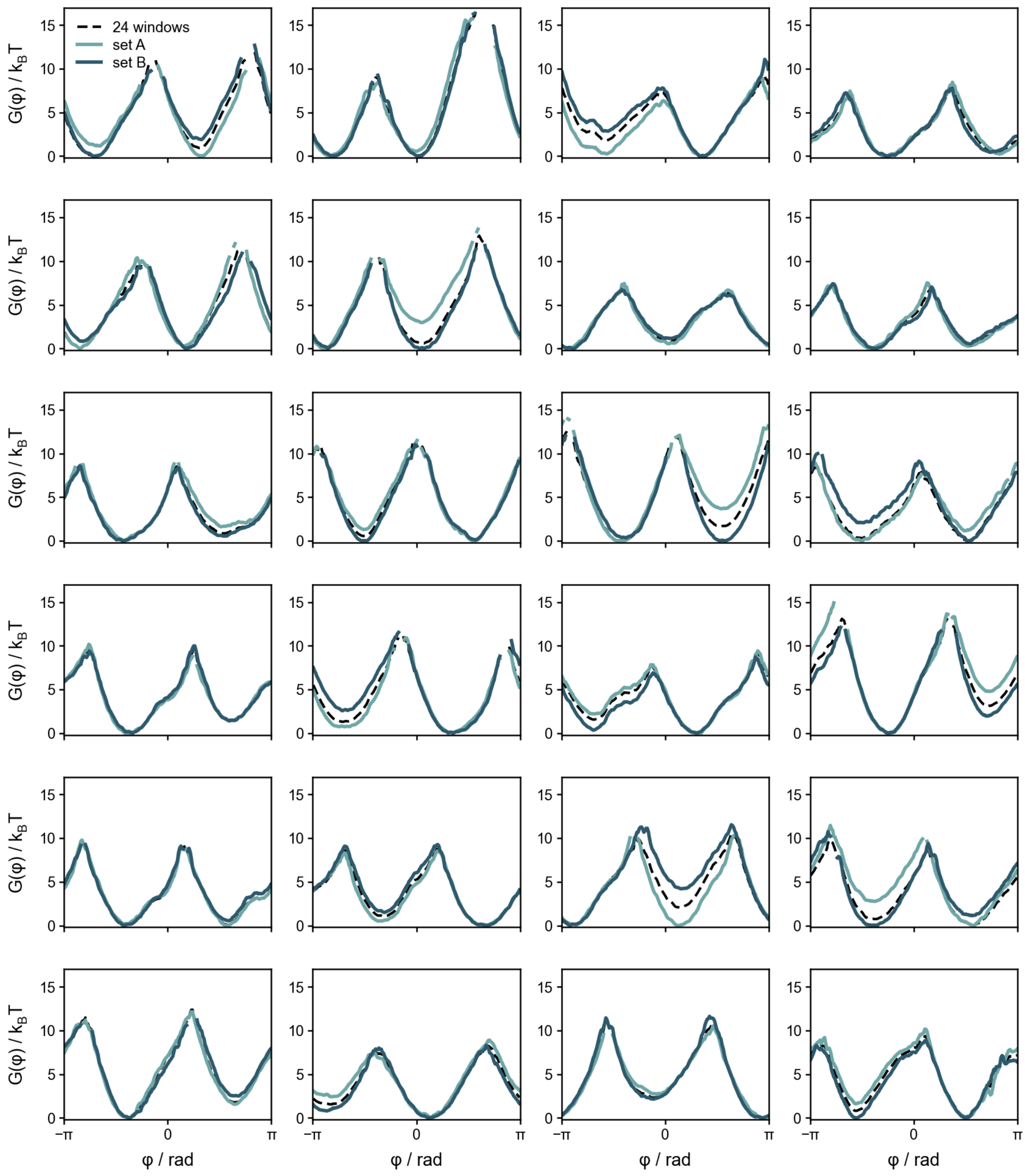

**Figure S23: Free-energy reconstruction after artificial downsampling of densely sampled umbrella windows.** Exemplary target-rotor free-energy landscapes reconstructed from the complete $15^{\circ}$-spaced recordings and from two complementary $30^{\circ}$-spaced subsets (A and B) obtained by retaining alternating umbrella windows.

### 6 Influence of bias voltage

#### 6.1 Steering voltage and restraint potentials

The steering voltage determines the strength of the experimentally realized restraint potential and therefore controls how strongly the rotor is confined around the commanded field orientation. We characterized the calibration-rotor response at $200\,\mathrm{V}$, $250\,\mathrm{V}$, and $300\,\mathrm{V}$.

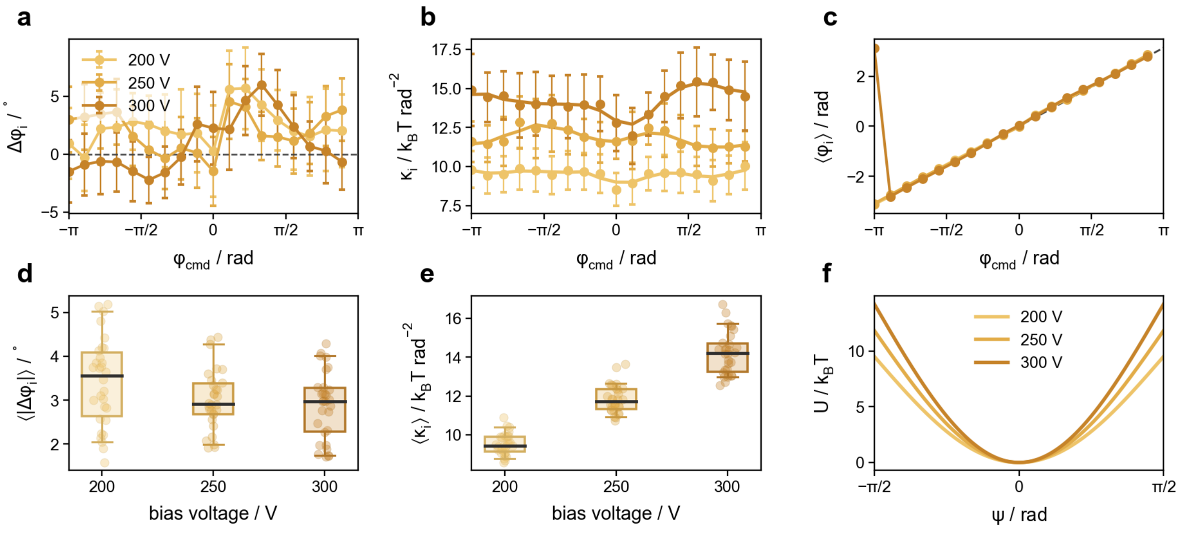

**Figure S24: Voltage dependence of the experimentally inferred restraint potentials.** **a** Deviation of the inferred bias center from the commanded field orientation, $\Delta\varphi_i$, across umbrella windows at $200\,\mathrm{V}$, $250\,\mathrm{V}$, and $300\,\mathrm{V}$. **b** Corresponding inferred restraint strengths, $\kappa_i$. **c** Mean inferred bias center, $\langle\varphi_i\rangle$, as a function of commanded field orientation. **d** Distribution of the mean absolute bias-center deviation, $\langle|\Delta\varphi_i|\rangle$, across calibration rotors. **e** Distribution of the mean inferred restraint strength, $\langle\kappa_i\rangle$, across calibration rotors. **f** Resulting cosine restraint potentials for the three steering voltages. Points in **a,b** show ensemble means and error bars indicate standard deviations across calibration rotors; points in **d,e** represent individual calibration rotors and box plots summarize their distributions.

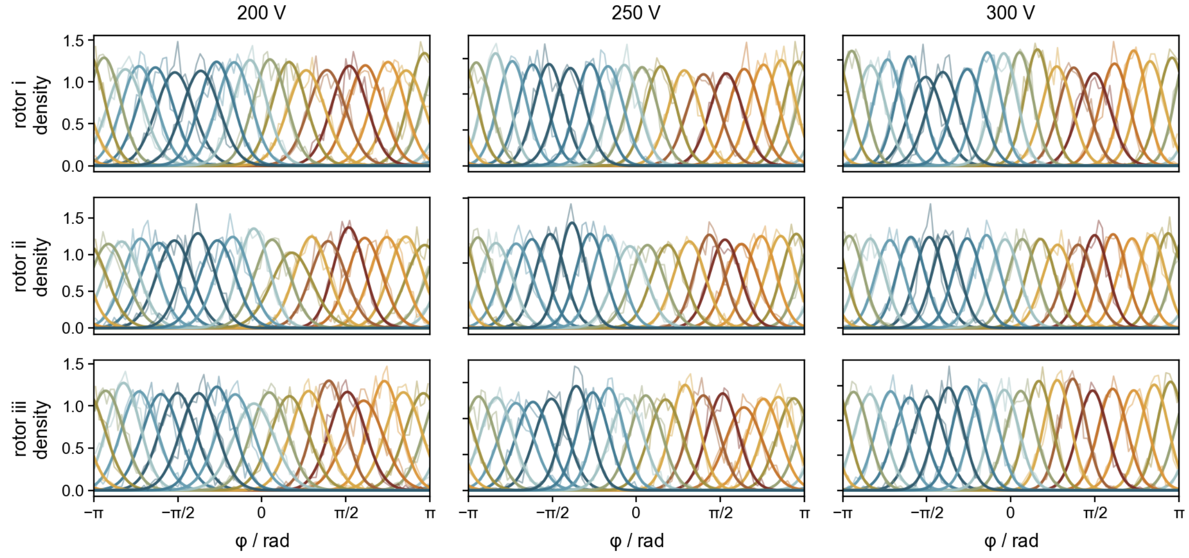

**Figure S25: Voltage dependence of angular confinement in individual calibration rotors.** Angular probability densities across the umbrella windows for three exemplary calibration rotors at $200\,\mathrm{V}$, $250\,\mathrm{V}$, and $300\,\mathrm{V}$. Thin lines show the measured angular distributions and thick lines the corresponding fitted distributions. Colors indicate the commanded field orientations. Increasing steering voltage produces progressively stronger angular confinement while maintaining coverage across the full $2\pi$ range.

#### 6.2 Autocorrelation, stationarity and connectivity as function of bias strength

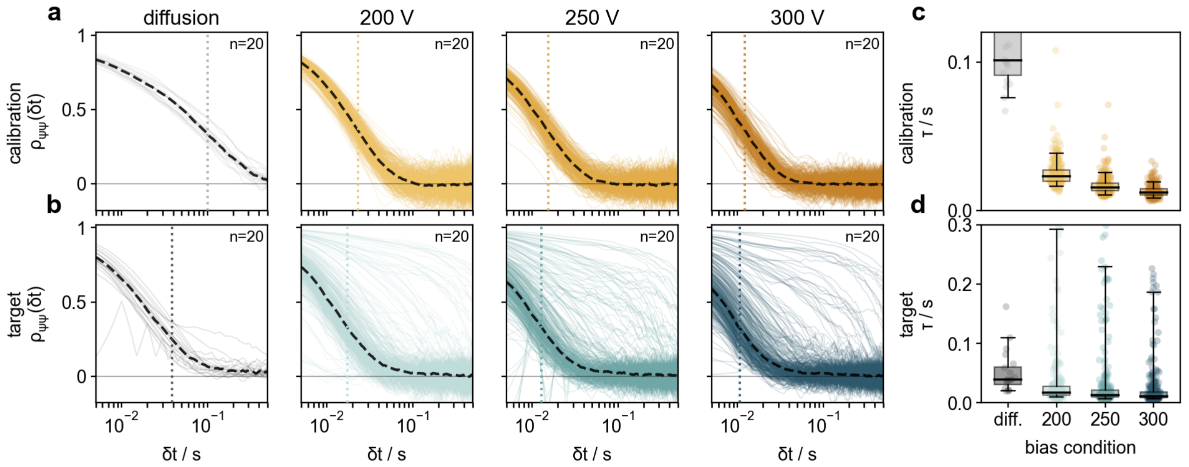

**Figure S26: Voltage dependence of temporal correlations during umbrella sampling. a,b** Angular autocorrelation functions of calibration and target rotors, respectively, during passive diffusion and umbrella sampling at $200\,\mathrm{V}$, $250\,\mathrm{V}$, and $300\,\mathrm{V}$. Thin lines show individual trajectories and dashed lines the corresponding ensemble-averaged autocorrelation functions. Dotted vertical lines indicate the extracted correlation times. **c,d** Distributions of the corresponding correlation times, $\tau$, for calibration and target rotors, respectively. Points represent individual trajectories and box plots summarize their distributions. Increasing steering voltage reduces the correlation time, particularly for the calibration rotors.

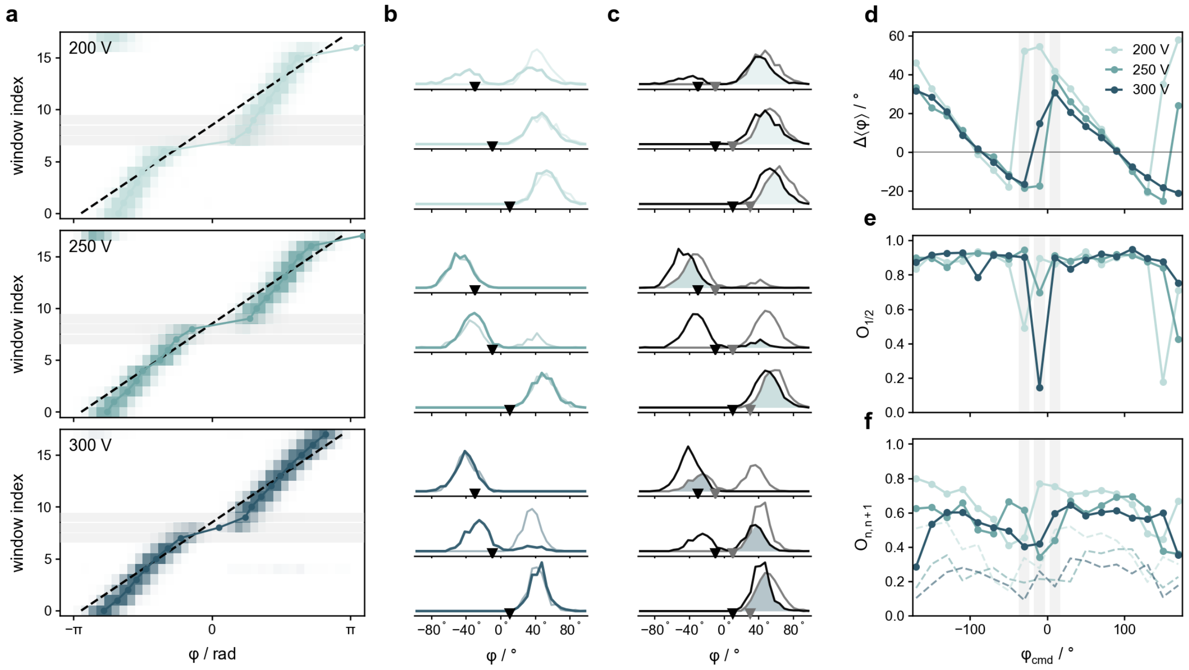

**Figure S27: Stationarity and statistical connectivity of target-rotor sampling as a function of steering voltage. a** Angular distributions of an exemplary target rotor across the umbrella windows at $200\,\mathrm{V}$, $250\,\mathrm{V}$, and $300\,\mathrm{V}$. The dashed line indicates the commanded field orientation and points indicate the mean angular position in each window. **b** Angular distributions sampled during the first and second halves of three exemplary umbrella windows. Triangles indicate their mean angular positions. **c** Angular distributions of neighboring umbrella windows for the same examples; shaded regions indicate their overlap. **d** Mean angular deviation of the target rotor from the calibration-rotor ensemble, $\langle\varphi_\mathrm{t}\rangle - \langle\varphi_\mathrm{c}\rangle$, as a function of commanded field orientation. **e** Within-window overlap, $O_{1/2}$, between the first and second halves of each umbrella window. **f** Overlap between neighboring, $O_{i,i+1}$, and next-nearest-neighbor, $O_{i,i+2}$, umbrella windows. Solid and dashed lines denote neighboring and next-nearest-neighbor overlap, respectively.

#### 6.3 Insufficient barrier sampling at low bias strength

The reduced confinement at low steering voltage becomes particularly relevant in high-energy regions of the intrinsic landscape. At $200\,\mathrm{V}$, several target rotors showed substantial deviations in the reconstructed barrier regions, whereas the $250\,\mathrm{V}$ and $300\,\mathrm{V}$ reconstructions were generally more consistent with each other and with the bias-free landscape where the latter was available (Fig. S28).

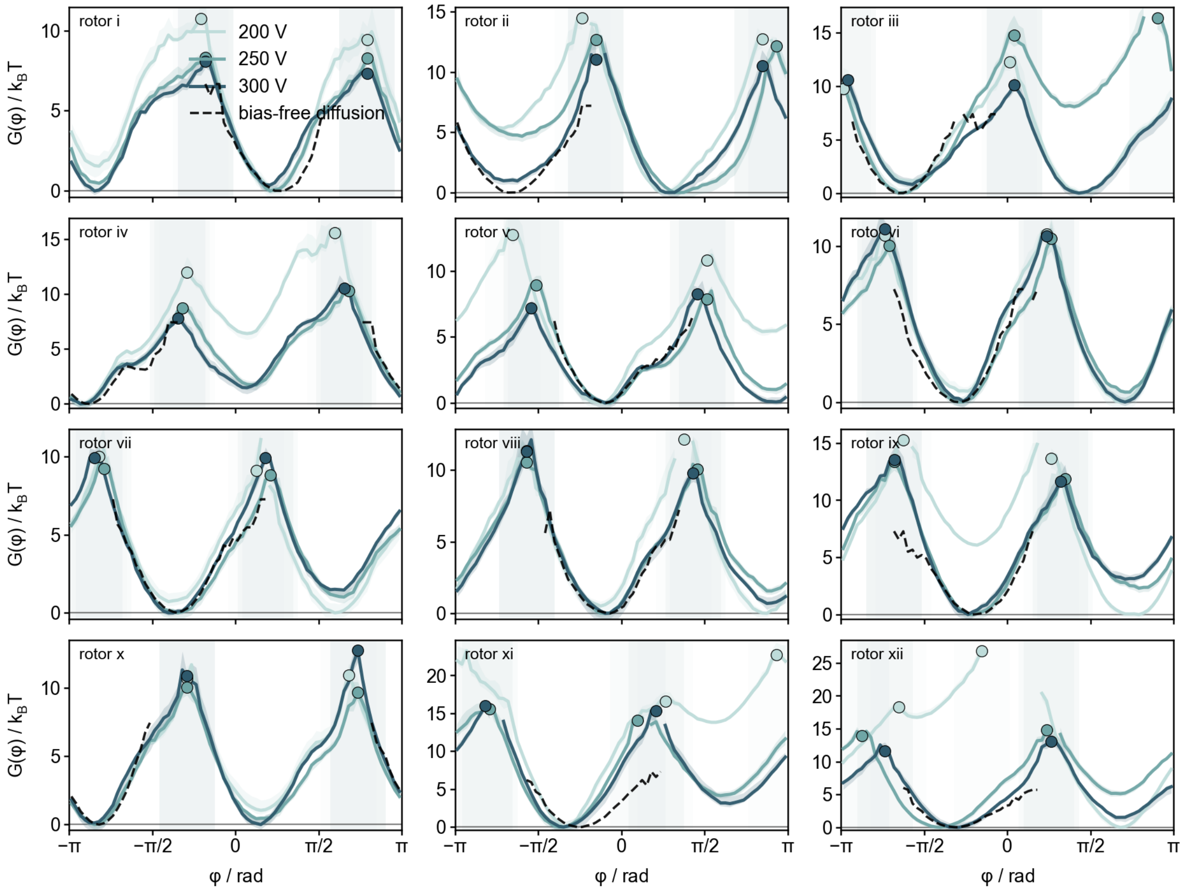

**Figure S28: Bias-voltage dependence of reconstructed free-energy landscapes.** Free-energy landscapes of individual target rotors reconstructed from umbrella-sampling measurements at $200\,\mathrm{V}$, $250\,\mathrm{V}$, and $300\,\mathrm{V}$. Dashed black lines show the corresponding landscapes obtained from bias-free diffusion over the angular range sampled during passive diffusion. Circles indicate the identified barrier positions for each reconstruction, and shaded regions indicate the angular regions around the barriers.

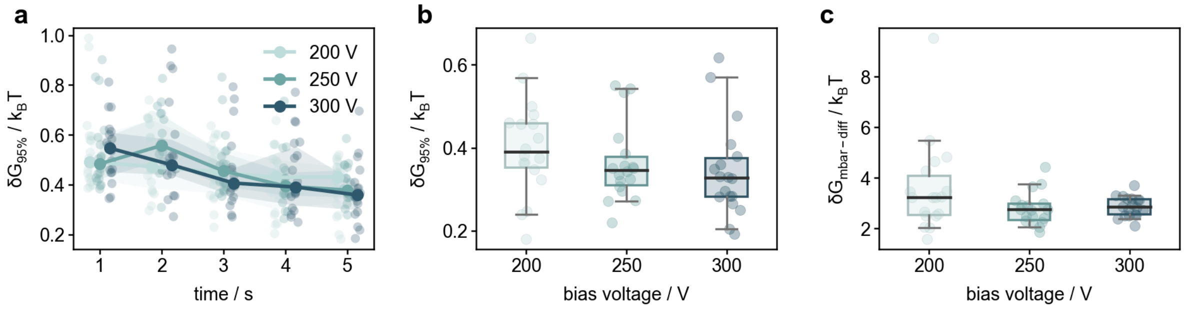

**Figure S29: Bias-voltage dependence of reconstruction uncertainty and agreement with bias-free diffusion. a** Median angular half-width of the 95% bootstrap confidence interval, $\delta G_{95\%}$, for each reconstructed free-energy landscape as a function of the amount of data used per umbrella window. Small points represent individual target rotors; lines show the median across rotors and shaded regions the interquartile range for each bias voltage. **b** Distribution of $\delta G_{95\%}$ for the reconstructions using the full $5\,\mathrm{s}$ acquisition time per umbrella window. **c** RMSE between the MBAR-reconstructed free-energy landscape and the corresponding landscape obtained from bias-free diffusion over the angular range sampled by both measurements. A constant energy offset was allowed before calculating the RMSE. Points in **b,c** represent individual target rotors; box plots show the median and interquartile range, with whiskers spanning the 5th–95th percentiles.

### 7 Influence of exposure time averaging

#### 7.1 Exposure averaging of restrained angular motion

The bias induced by the electric field during an umbrella window can be described by a cosine-shaped angular potential. Locally around its minimum, it can be approximated by a harmonic potential,k

$$U(\tilde{\psi}) = \frac{1}{2}\tilde{\kappa}\tilde{\psi}^2,$$

where $\tilde{\psi} = \varphi - \varphi_{\mathrm{cmd}}$ is the angular displacement and $\tilde{\kappa}$ is the trap stiffness / bias strength in units of $k_{\mathrm{B}}T\,\mathrm{rad}^{-2}$. The corresponding restoring torque is $-\mathrm{d}U/\mathrm{d}\tilde{\psi} = -\tilde{\kappa}\tilde{\psi}$. In the overdamped regime, the motion of the rotor in this harmonic potential is described by the Langevin equation,

$$\gamma\frac{\mathrm{d}\tilde{\psi}}{\mathrm{d}t} = -\tilde{\kappa}\tilde{\psi} + \sqrt{2\gamma k_{\mathrm{B}}T}\,\eta(t),$$

where $\gamma$ is the rotational friction coefficient and $\eta(t)$ represents Gaussian white noise with zero mean and $\langle\eta(t)\,\eta(t')\rangle = \delta(t-t')$. Introducing the rotational diffusion coefficient $D = k_{\mathrm{B}}T/\gamma$ and the characteristic relaxation time $\tau = \gamma/\tilde{\kappa}$, the Langevin equation can be rewritten as

$$\frac{\mathrm{d}\tilde{\psi}}{\mathrm{d}t} = -\frac{1}{\tau}\tilde{\psi} + \sqrt{2D}\,\eta(t),$$

which describes an Ornstein–Uhlenbeck process. This equation describes the instantaneous angular dynamics of the rotor, whereas the experimentally recorded trajectory is affected by the finite camera exposure time. Rather than directly measuring the instantaneous angular displacement $\tilde{\psi}(t)$, the camera averages the rotor position over the exposure time $\Delta t_{\mathrm{e}}$. The resulting angular displacement recorded in frame $n$ starting at $t_n$ is thus

$$\psi_n = \frac{1}{\Delta t_{\mathrm{e}}}\int_{t_n}^{t_n+\Delta t_{\mathrm{e}}} \tilde{\psi}(t)\,\mathrm{d}t.$$

In our experiments, the bias strength $\kappa$ is inferred from the width of the angular distributions measured for the calibration rotors. While the underlying stiffness is determined by the variance of the instantaneous angular displacement $\langle\tilde{\psi}^2\rangle = k_{\mathrm{B}}T/\tilde{\kappa}$, the camera only provides the exposure-averaged coordinate $\psi_n$ and thus its variance $\langle\psi_n^2\rangle$. The effect of finite exposure can therefore be determined via

$$\langle\psi_n^2\rangle = \frac{1}{\Delta t_{\mathrm{e}}^2}\int_{t_n}^{t_n+\Delta t_{\mathrm{e}}}\int_{t_n}^{t_n+\Delta t_{\mathrm{e}}} \langle\tilde{\psi}(t)\tilde{\psi}(t')\rangle\;\mathrm{d}t\,\mathrm{d}t',$$

since the relation between the measured and instantaneous variances depends on the temporal correlations of the underlying rotor motion. For a stationary Ornstein–Uhlenbeck process, the covariance between displacements separated by a time lag $\Delta t_{\mathrm{e}}$ is decaying exponentially

$$\langle\tilde{\psi}(t)\,\tilde{\psi}(t+s)\rangle = \frac{k_{\mathrm{B}}T}{\tilde{\kappa}}\exp\left(-\frac{|s|}{\tau}\right).$$

Substituting this covariance into the integral above and evaluating it for the camera exposure $\Delta t_{\mathrm{e}}$ gives

$$\langle\psi_n^2\rangle = \frac{k_{\mathrm{B}}T}{\tilde{\kappa}}\,\frac{2\tau}{\Delta t_{\mathrm{e}}^2}\left[\Delta t_{\mathrm{e}} - \tau\left(1 - e^{-\Delta t_{\mathrm{e}}/\tau}\right)\right] = \frac{k_{\mathrm{B}}T}{\tilde{\kappa}}\,B(\Delta t_{\mathrm{e}},\tau),$$

where $B(\Delta t_{\mathrm{e}},\tau)$ is the exposure-averaging factor. Finite exposure times $\Delta t_{\mathrm{e}} > 0$, lead to $B(\Delta t_{\mathrm{e}},\tau) < 1$. Hence, temporal averaging reduces the measured angular variance and narrows the recorded angular distribution. The apparent stiffness determined via $\kappa = k_{\mathrm{B}}T/\langle\psi_n^2\rangle$ is connected to the underlying stiffness via

$$\kappa(\Delta t_{\mathrm{e}}) = \frac{\tilde{\kappa}}{B(\Delta t_{\mathrm{e}},\tau)}.$$

For exposure times $\Delta t_{\mathrm{e}} \ll \tau$, $\kappa \longrightarrow \tilde{\kappa}$, whereas for $\Delta t_{\mathrm{e}} \sim \tau$, temporal averaging increasingly suppresses the measured angular fluctuations, resulting in an apparent increase in stiffness.

#### 7.2 Extracting the underlying bias strength and relaxation time

To determine the underlying bias strength experimentally, the same calibration rotors were recorded at different exposure times $\Delta t_\mathrm{e} \in \{5, 7, 10, 20\}\ \mathrm{ms}$. For each rotor and exposure time, the measured stiffness $\kappa$ was determined from the angular variance in each umbrella window and subsequently averaged across windows (Fig. S30). The resulting exposure-dependent stiffnesses were fitted separately for each rotor, with $\tilde{\kappa}$ and $\tau$ as fit parameters, by minimizing the logarithmic residuals

$$r(\Delta t_\mathrm{e}) = \log\left[\kappa(\Delta t_\mathrm{e})\right] - \log\left[\frac{\tilde{\kappa}}{B(\Delta t_\mathrm{e}, \tau)}\right].$$

The fitted $\tilde{\kappa}$ therefore represents the underlying bias strength in the absence of exposure averaging, whereas $\tau$ describes the characteristic relaxation timescale of the rotor within the restraint potential (Fig. S30c). Finite camera exposure additionally reduces the temporal information contained in the recorded trajectories. As $\Delta t_\mathrm{e}$ approaches the relaxation time $\tau$, an increasing fraction of the rotor relaxation occurs within a single exposure and is therefore temporally averaged, reducing the precision with which the underlying relaxation dynamics can be resolved.

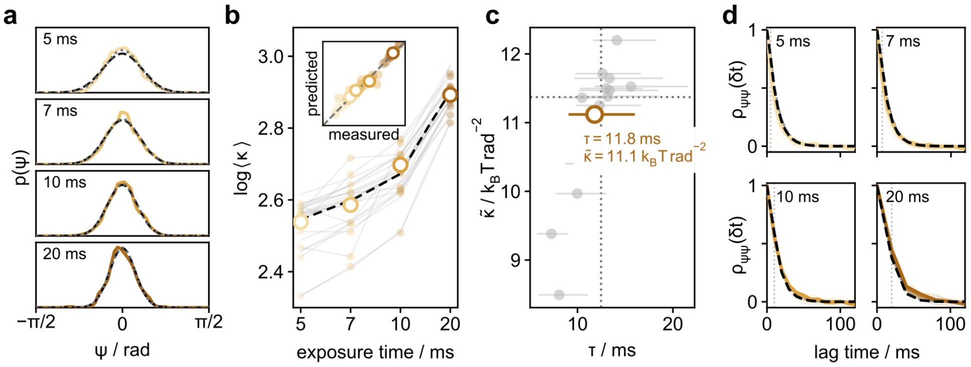

**Figure S30: Exposure-time dependence of the measured bias strength and Ornstein–Uhlenbeck analysis. a** Exemplary angular probability distributions $p(\psi)$ of a calibration rotor recorded at exposure times $\Delta t_\mathrm{e} = 5$, 7, 10 and 20 ms (colors) and corresponding Gaussian fits. **b** Mean measured stiffness $\langle\kappa\rangle$ as a function of exposure time. The dashed black line shows the median prediction from the Ornstein–Uhlenbeck model. Inset, measured versus model-predicted stiffnesses. **c** Underlying bias strength $\tilde{\kappa}$ and relaxation time $\tau$ obtained from the Ornstein–Uhlenbeck fit for individual calibration rotors. Dotted lines indicate the ensemble medians. **d** Autocorrelation functions of the angular displacement for the four exposure times. Coloured lines show individual calibration rotors and the dashed black lines the autocorrelation predicted Ornstein–Uhlenbeck model.

#### 7.3 Reconstruction of energy landscapes

Increasing the exposure time not only enhances temporal averaging but also reduces the number of recorded positions within each fixed-duration umbrella window of 5 s. To distinguish these effects, we compared reconstructions (i) using all recorded positions, (ii) using an equal number $N$ of consecutive positions from the beginning of each window, and (iii) using the same number $N$ of positions evenly spaced across the full window duration (Fig. S32). This allowed us to test whether differences in reconstruction quality arise from the reduced number of recorded positions or from the exposure time itself.

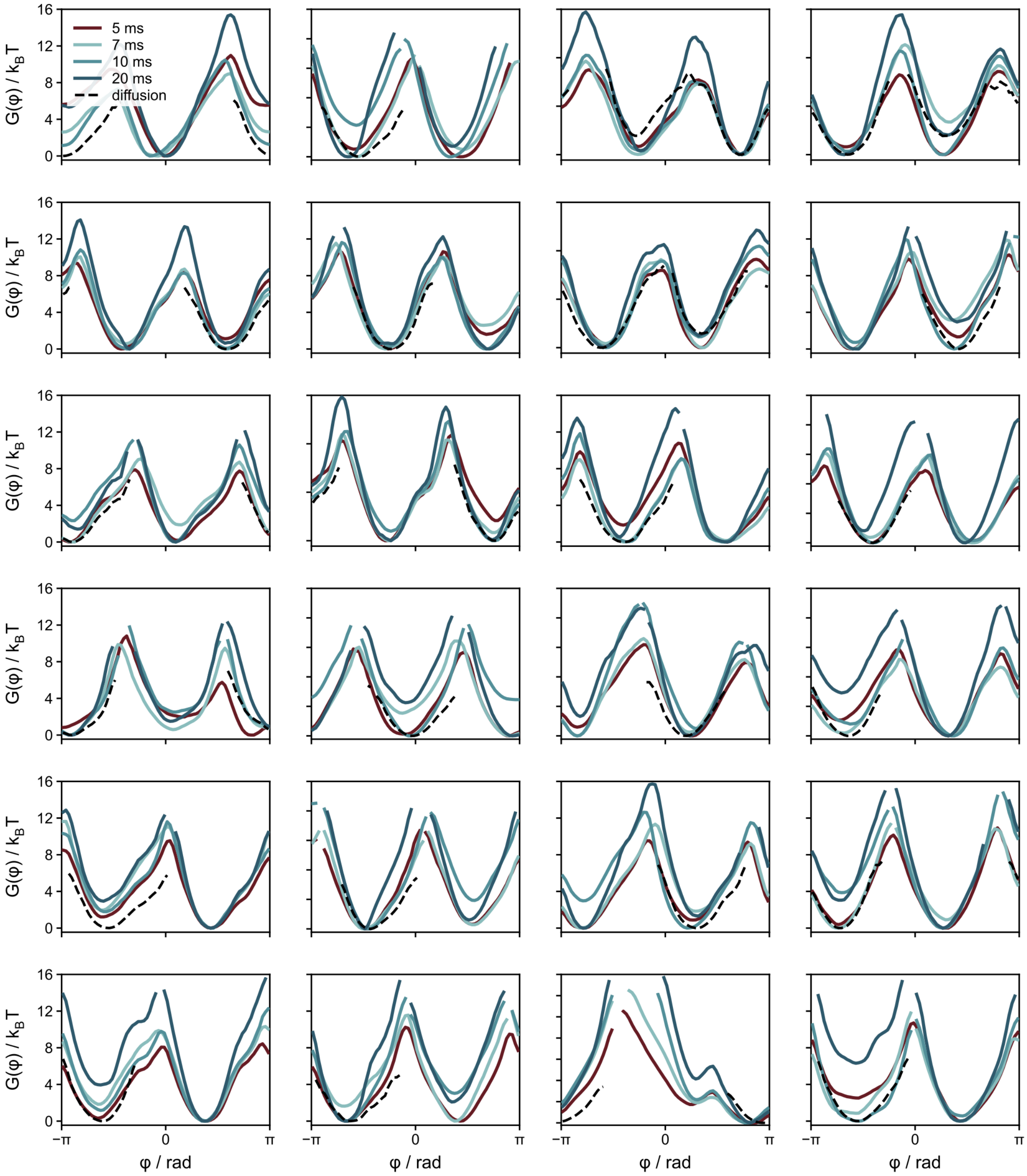

**Figure S31: Free-energy landscapes reconstructed at different camera exposure times using a fixed window duration.** Free-energy landscapes of individual target rotors reconstructed from umbrella-sampling measurements at exposure times of 5, 7, 10 and 20 ms using a fixed duration of 5 s per umbrella window. Dashed black lines show the corresponding landscapes obtained from bias-free diffusion measurements.

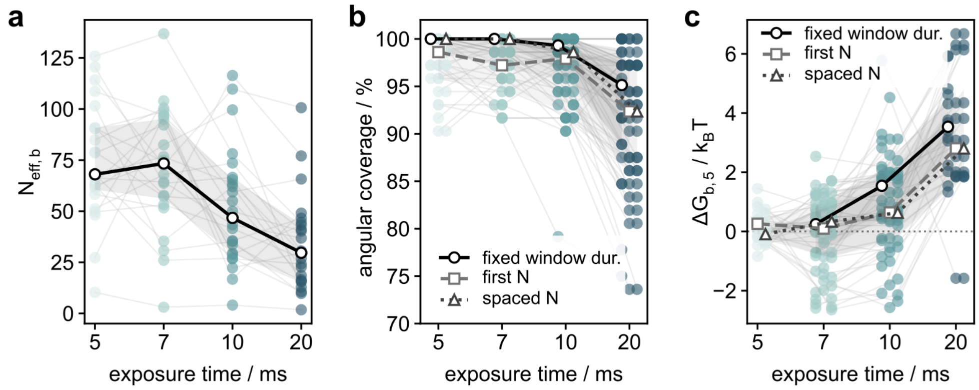

**Figure S32: Effect of camera exposure time on reconstruction quality. a** Effective number of samples in the barrier region, $N_{\mathrm{eff},b}$, as a function of exposure time evaluated in case of a fixed umbrella-window duration of 5 s. **b** Angular coverage of the reconstructed free-energy landscapes using all positions recorded within each 5-s umbrella window, the first $N$ positions (dashed), or $N$ positions evenly spaced across the full window (dotted). For the equal-sample-number reconstructions, $N = 250$ corresponds to the number of positions recorded during 5 s at the longest exposure time of 20 ms. **c** Difference in reconstructed barrier height, $\Delta G_{b,5}$, relative to the corresponding 5-ms reconstruction for the different exposure times and sampling strategies.

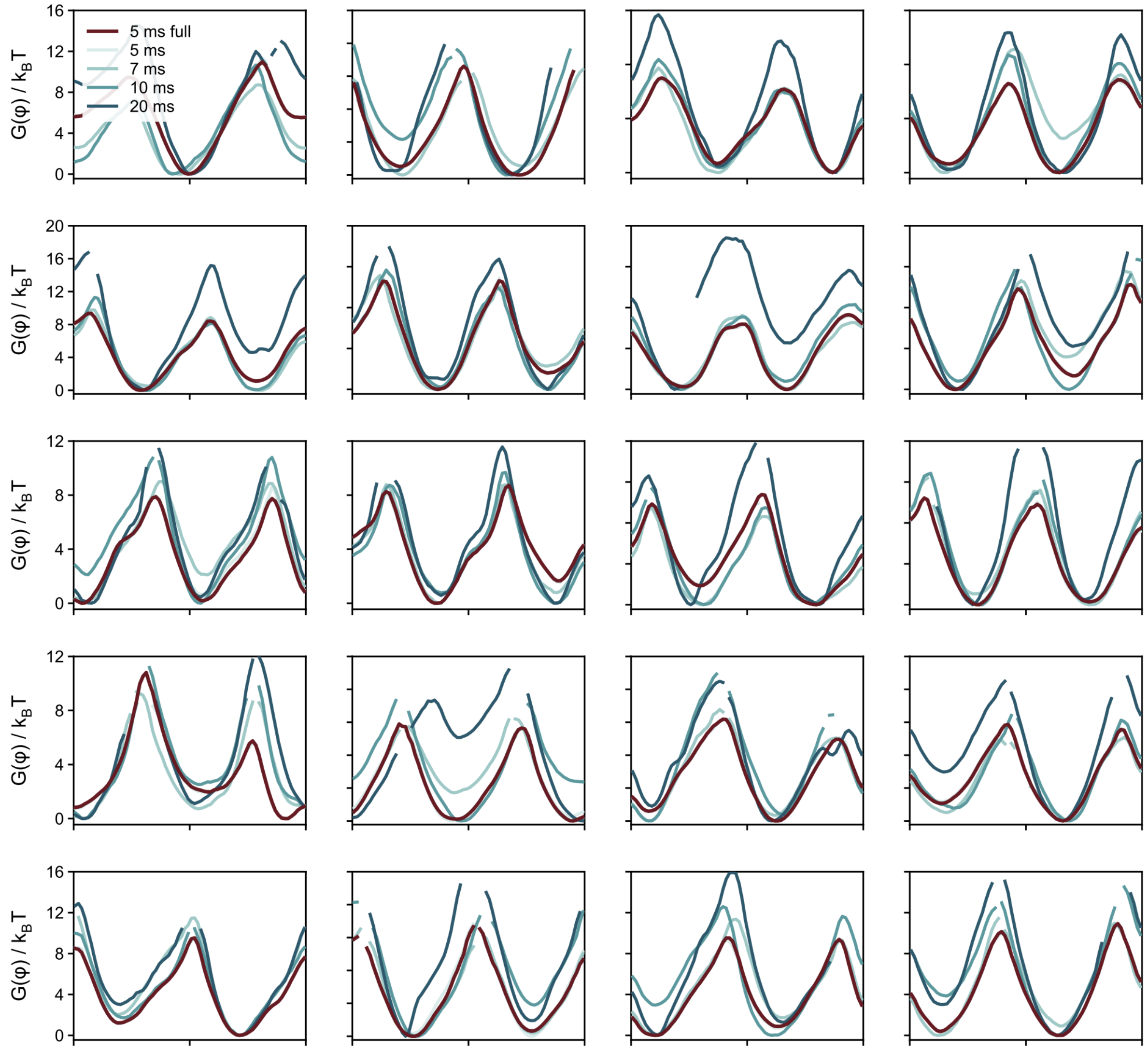

**Figure S33: Free-energy landscapes reconstructed using equal sample numbers.** Free-energy landscapes of individual target rotors reconstructed at exposure times of 5, 7, 10 and 20 ms using $N = 250$ positions evenly spaced across each 5-s umbrella window. Dashed black lines show the corresponding reconstructions obtained using all positions recorded at 5 ms exposure time.

### 8 Stator geometry shapes the rotary free-energy landscapes

#### 8.1 Outer-anchor stator variant

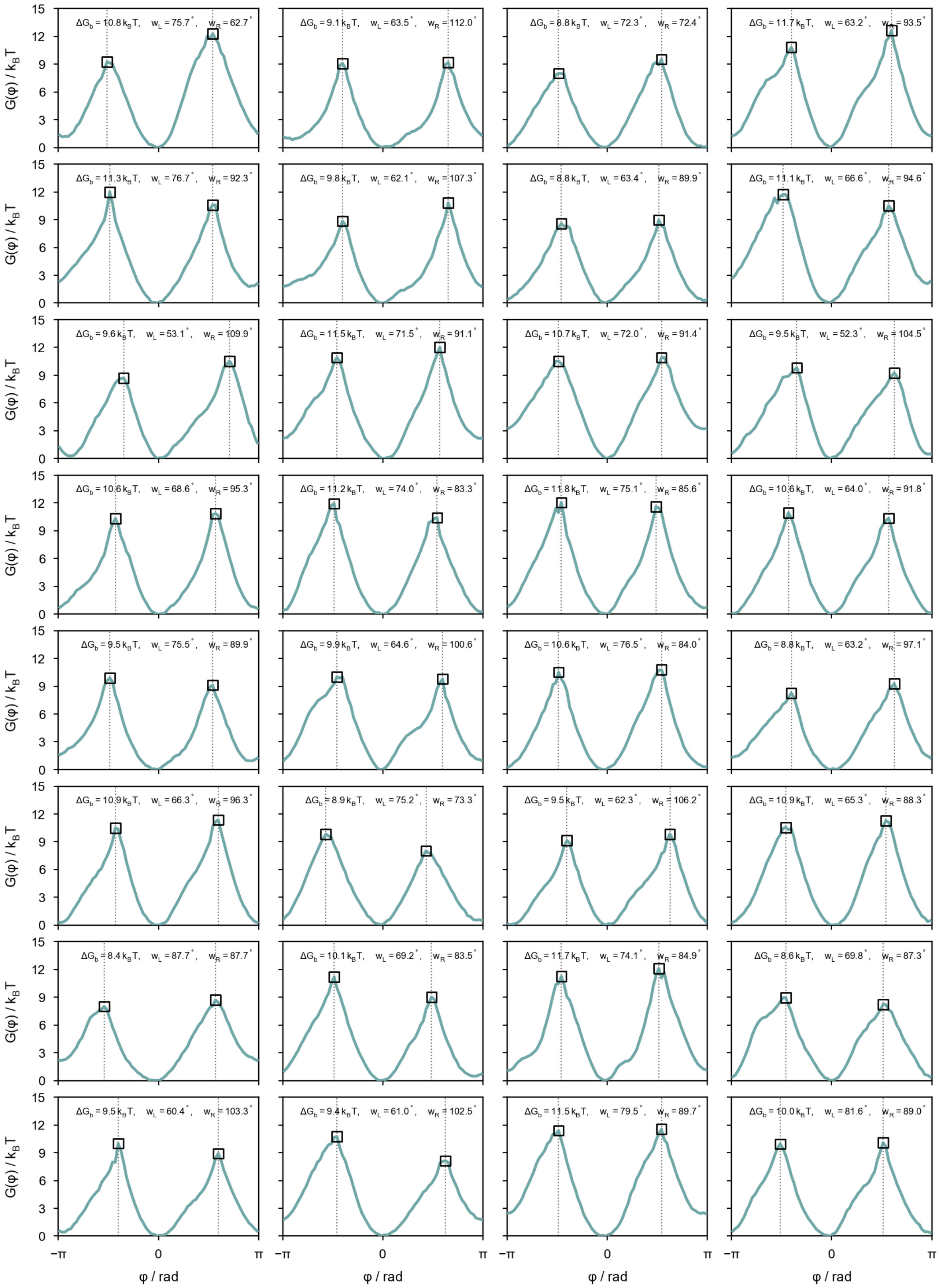

**Figure S34: Reconstructed free-energy landscapes for individual outer-anchor stator rotors.** Free-energy landscapes $G(\varphi)$ of individual rotors. Squares indicate the left and right barrier maxima, and dotted lines their angular positions. The mean barrier height $\Delta G_b$ is calculated from the two barrier maxima. The left and right barrier widths $w_L$ and $w_R$ denote the angular distances from the minimum to the respective crossings at 90% of the lower barrier height.

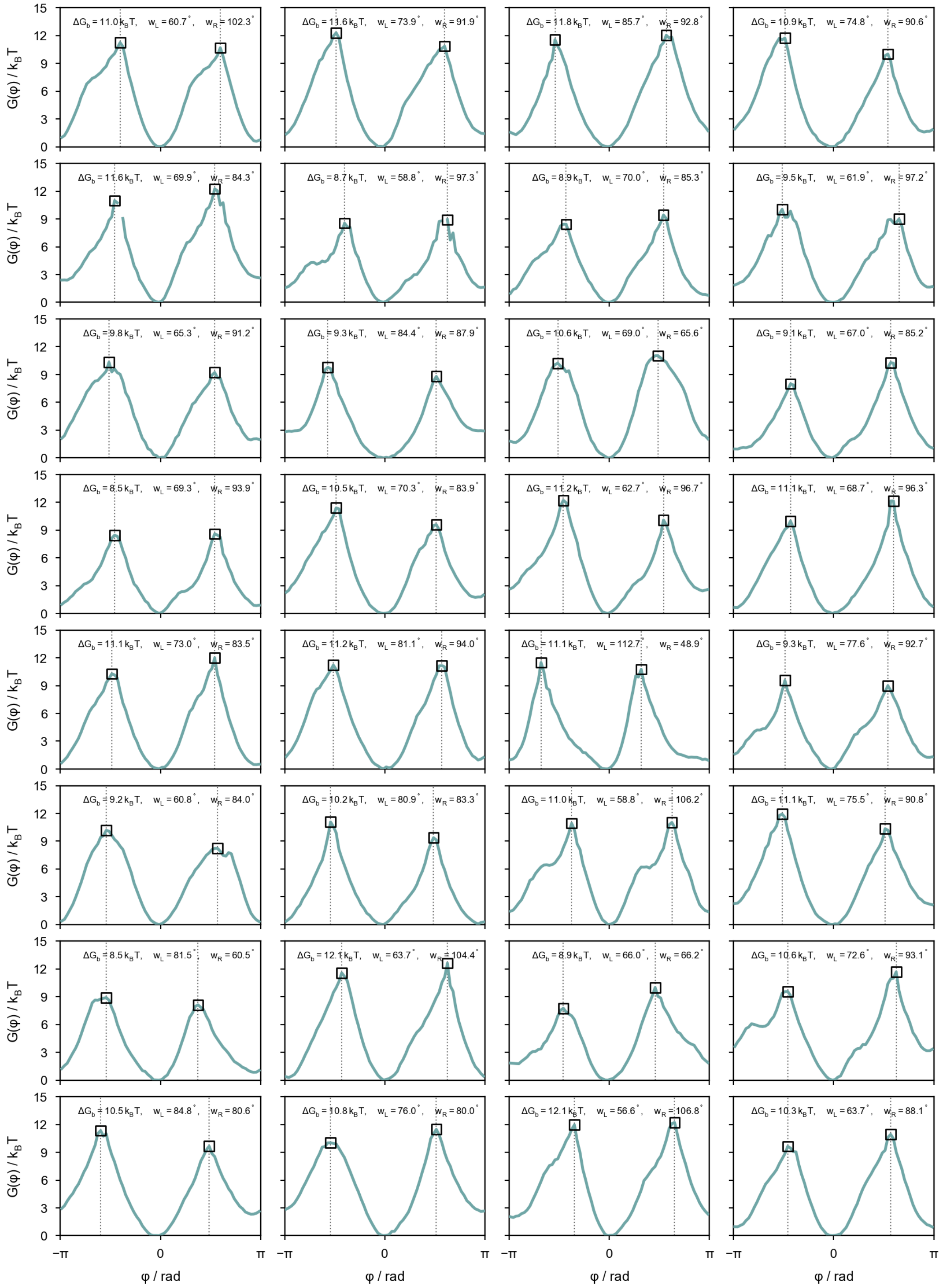

**Figure S35: Reconstructed free-energy landscapes for individual outer-anchor stator rotors (continued).**

#### 8.2 Dual-anchor stator variant

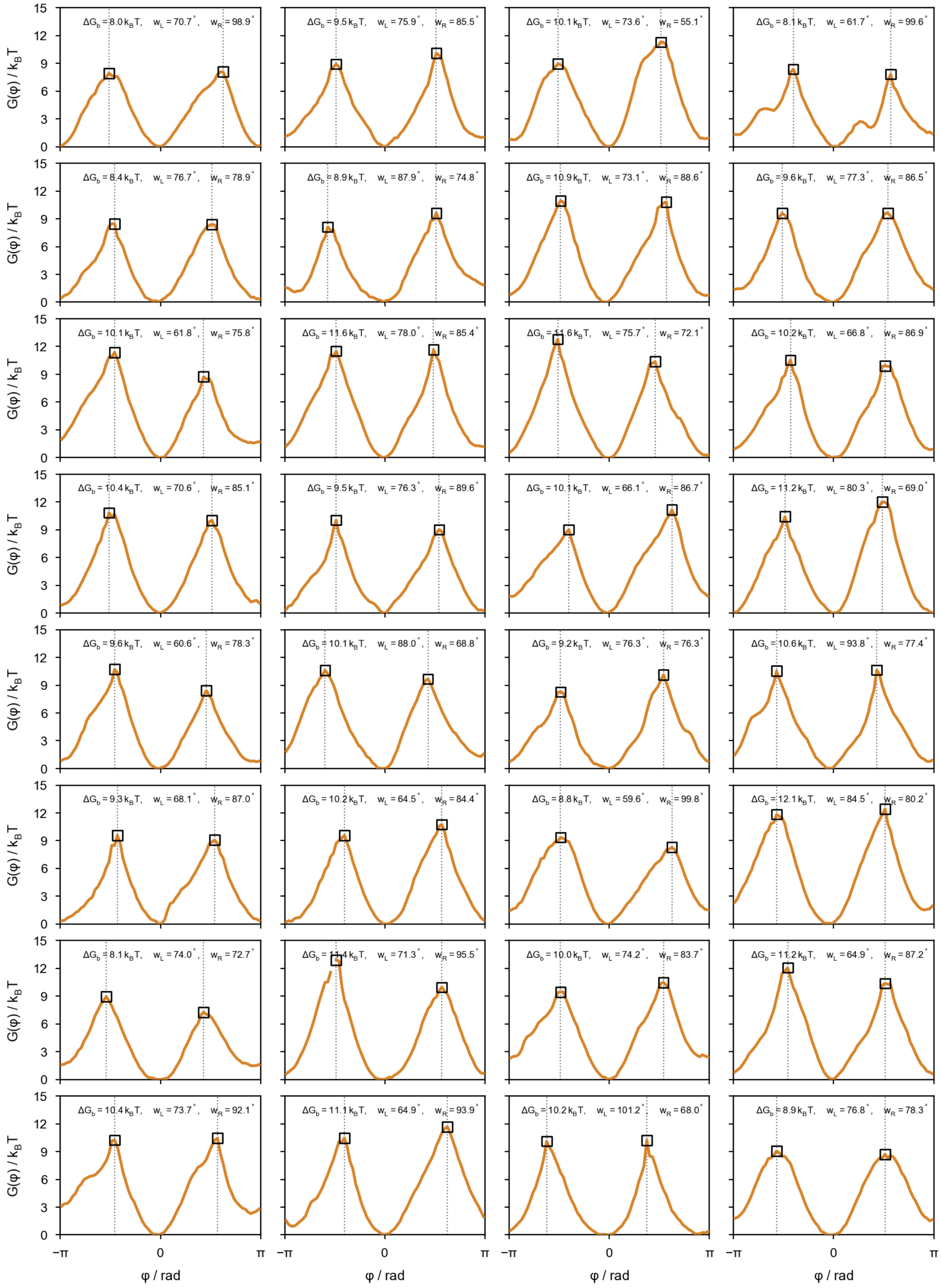

**Figure S36: Reconstructed free-energy landscapes for individual dual-anchor stator rotors.**

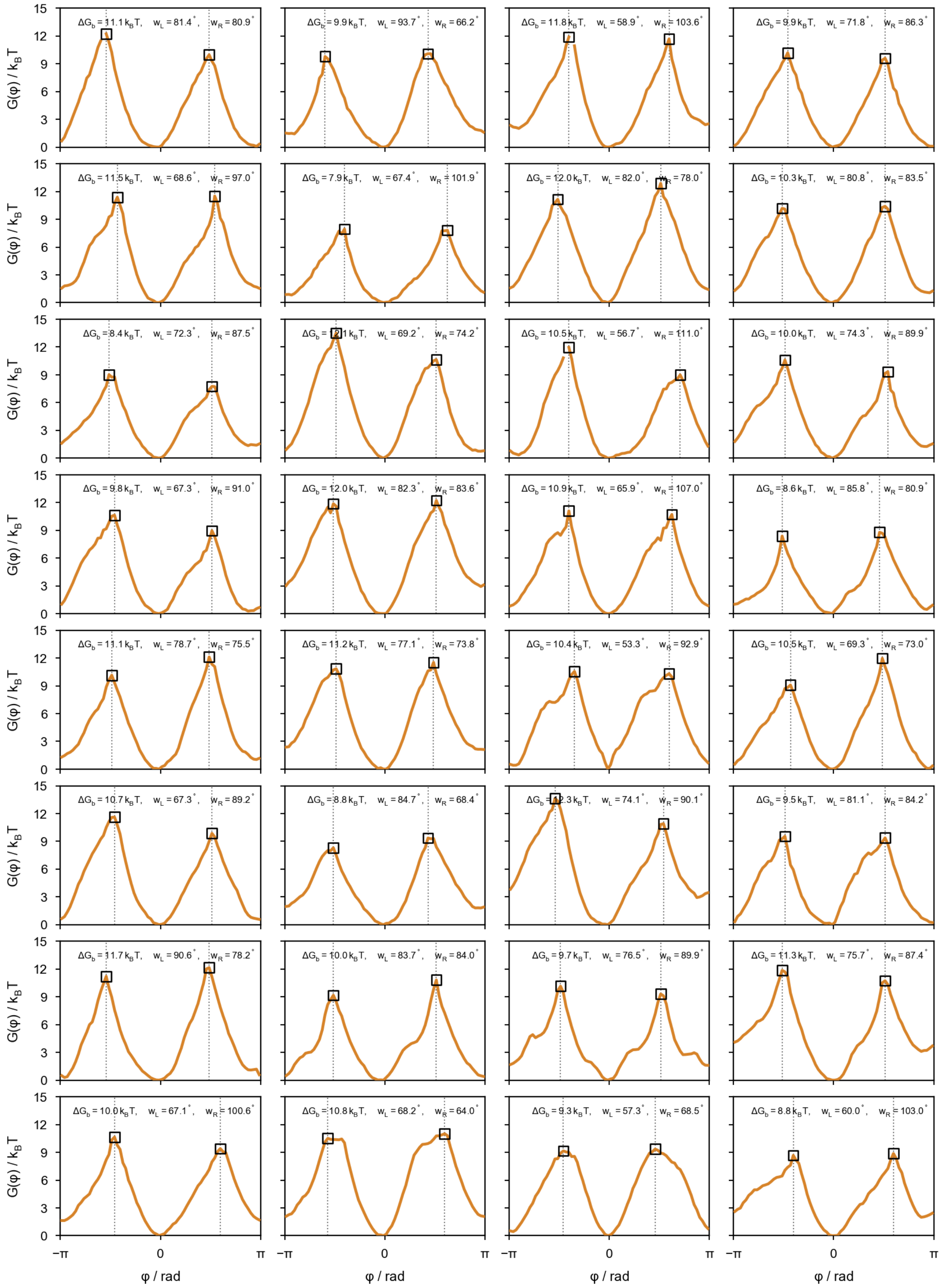

**Figure S37: Reconstructed free-energy landscapes for individual dual-anchor stator rotors (continued).**

#### 8.3 Inner-anchor stator variant

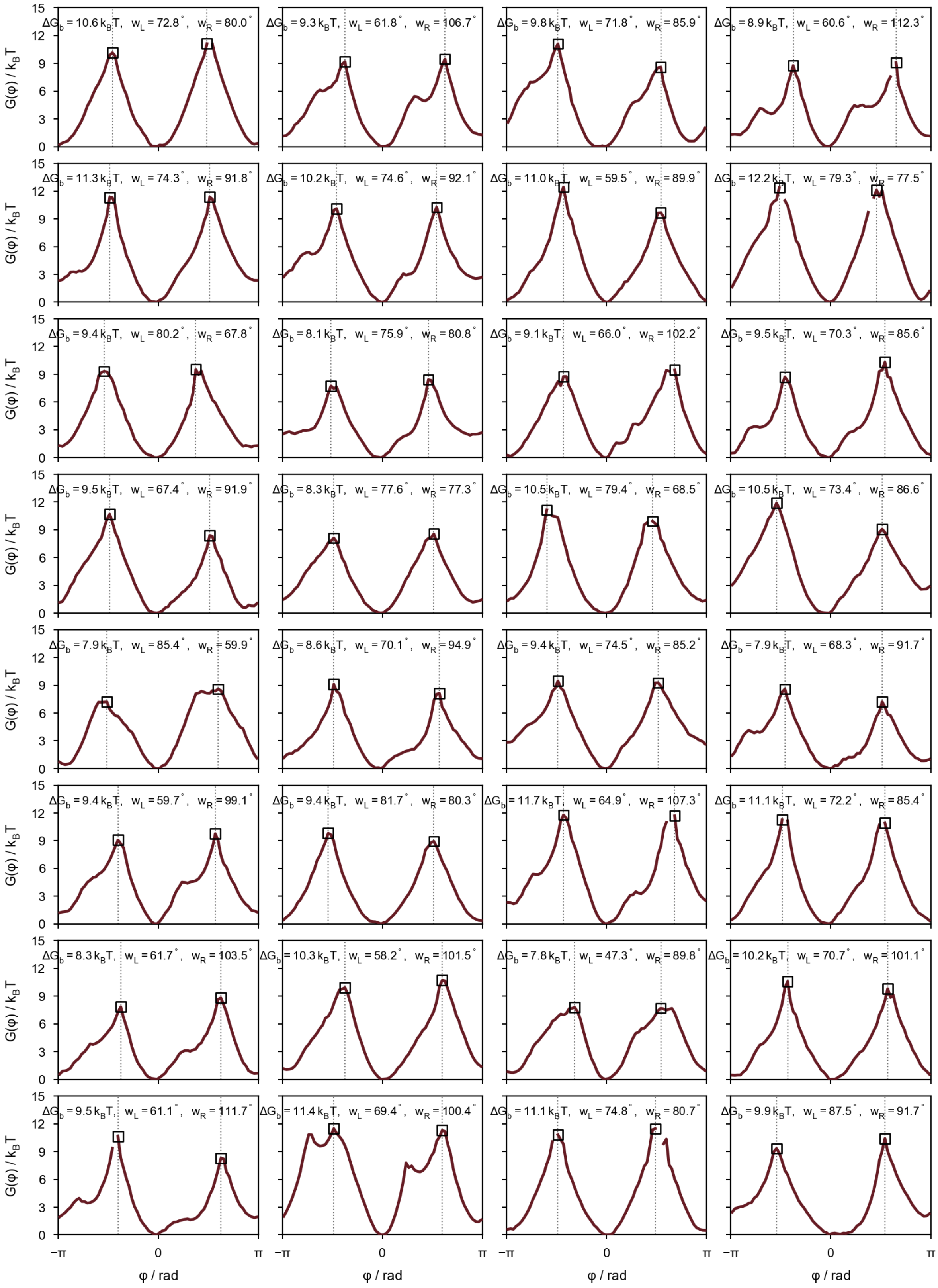

**Figure S38: Reconstructed free-energy landscapes for individual inner-anchor stator rotors.**

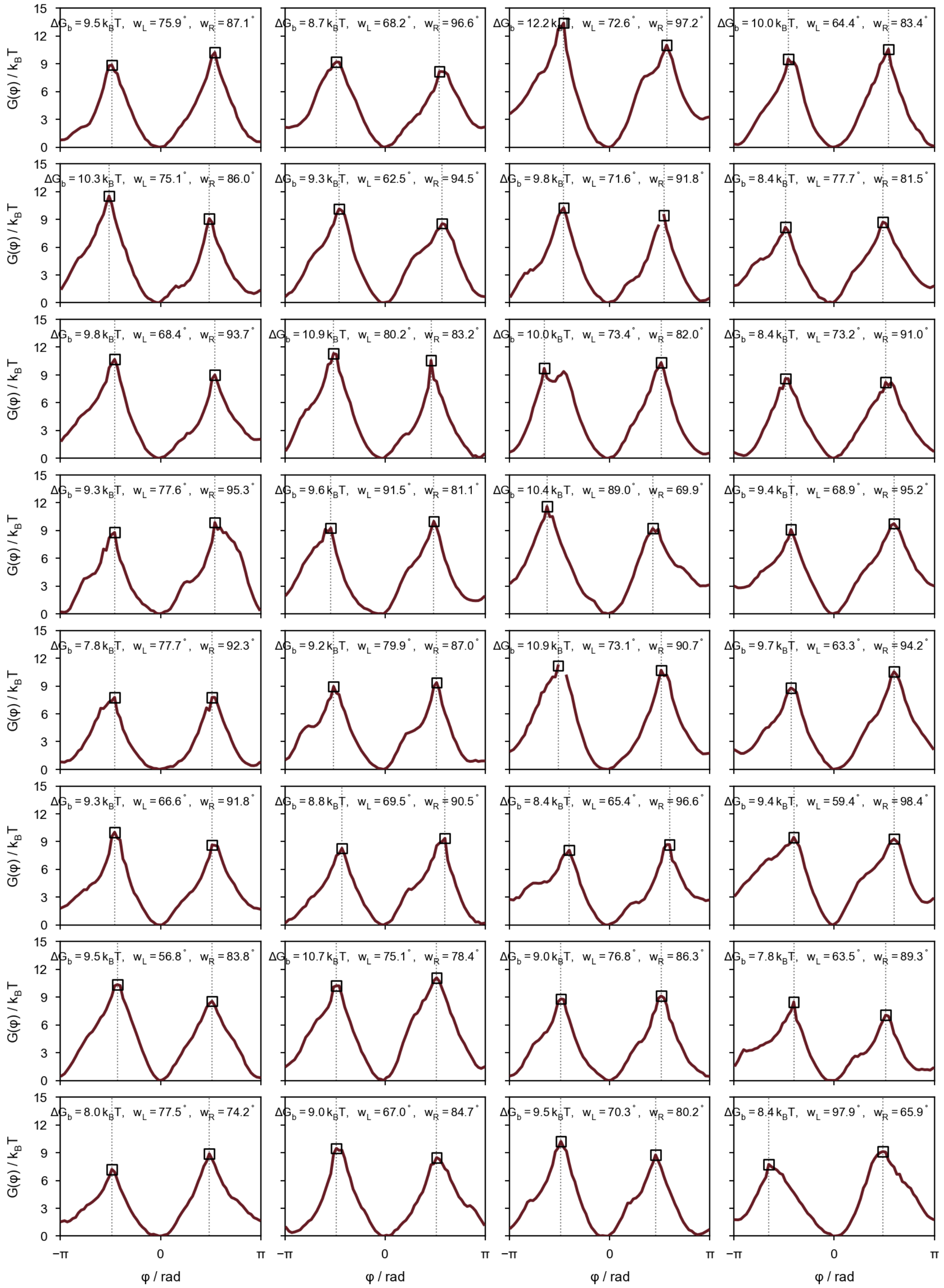

**Figure S39: Reconstructed free-energy landscapes for individual inner-anchor stator rotors (continued).**

### 9 Rotor dynamics in response to alternating rotating electric fields

We characterized the rotational dynamics of the stator variant with outer tether positions under an increasing electric field rotating at 1 Hz, with alternating directions evers 5 s. The majority of rotors preferentially followed counterclockwise rotation, consistent with the reduced restoring torque associated with the shallower right flank $G(\varphi)$.

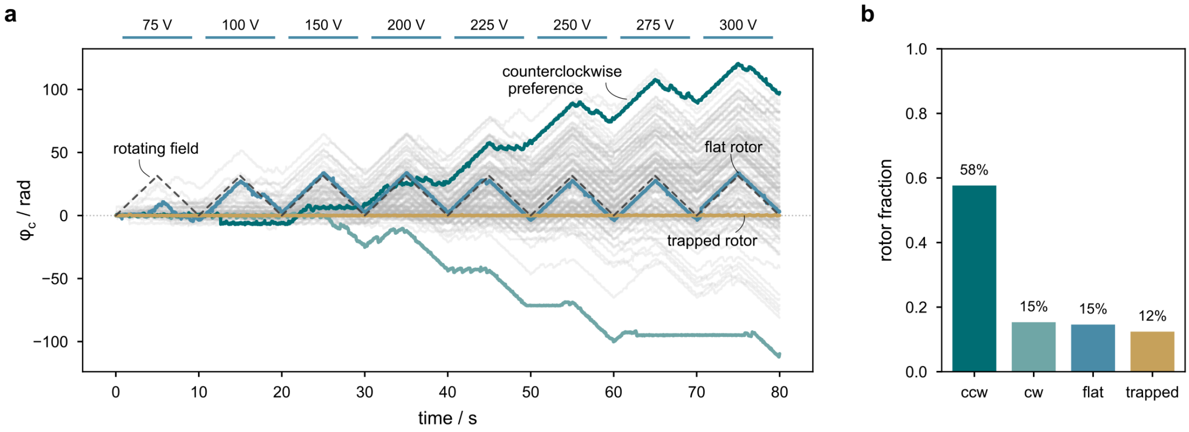

**Figure S40: Rotor dynamics under rotating fields. (a)** Angular trajectories of individual rotors under a rotating electric field alternating between counterclockwise and clockwise rotation at 1 Hz for increasing voltages. Gray lines represent individual rotor trajectories, while colored lines highlight representative counterclockwise (green), clockwise (light green), flat energy landscape (blue), and trapped (gold) rotors. The dashed line indicates the angular displacement of the rotating field. **(b)** Fraction of rotors classified according to their rotational response. The majority of rotors exhibit a preference for counterclockwise rotation.

### 10 DNA scaffold sequences and staples

**Table S1:** DNA sequences of the basic stator module without tether strands. The sequences are based on the 8064 nt-scaffold.

| function | position | sequence |
|---|---|---|
| fluoroescent marker | 30[87]28[88] | Atto488–GAAATAGCTATCCCAATCCAAATATGCGGGAG |
| biotin anchor | 24[55]22[40] | biotin-ttAAATCCAAGAAAACAAAATTAATTCCAAGTTACAAAATCGCGCAGAGG |
| biotin anchor | 24[151]22[136] | biotin-ttCAATAGTGCTTCTGTAAATCGTCGCGTAGATTTTCAGGTTTAACGTCA |
| biotin anchor | 32[87]30[88] | biotin-ttTTATCACCCCTTATTACGCAGTATGAAACAAT |
| biotin anchor | 32[119]30[120] | biotin-ttAATATTGAAATACATACATAAAGAGAATTG |
| biotin anchor | 40[151]38[152] | biotin-ttGGACTAAAGTTAAAGGCCGCTTTTCTGTATGG |
| biotin anchor | 40[55]38[56] | biotin-ttGAATACACTCTTAAACAGCTTGATGTTGAAAA |
| edge | 1[8]2[40] | TTTTTTTACCTTATGCGATTTTAAGAACTGGCTCAAATACTGC |
| edge | 3[12]2[12] | TTTTAATCCCCCTCAAATGCTCATAAATATTCATTGTTTT |
| edge | 5[12]4[12] | TTTTATAACAGTTGATTCCCGGAAGTTTCATTCCATTTTT |
| edge | 7[12]6[12] | TTTTTGTAGGTAAAGATTCACAATGCCTGAGTAATGTTTT |
| edge | 9[12]8[12] | TTTTAACGCCATCAAAAATATTTTTTAACCAATAGGTTTT |
| edge | 11[12]10[12] | TTTTGGGGATGTGCTGCAAGCGCCAGCTGGCGAAAGTTTT |
| edge | 13[12]12[12] | TTTTATTGGGCGCCAGGGTGGAGAGGCGGTTTGCGTTTTT |
| edge | 15[12]14[12] | TTTTAAATCGGAACCCTAAAGGTGCCGTAAAGCACTTTTT |
| edge | 17[12]16[12] | TTTTTCACGCAAATTAACCGTAAAAGAGTCTGTCCATTTT |
| edge | 19[12]18[12] | TTTTAACATCGCCATTAAAAGAACTGATAGCCCTAATTTT |
| edge | 21[12]20[12] | TTTTTATTAATTTTAAAAGTTCCTTTGCCCGAACGTTTTT |
| edge | 23[12]22[12] | TTTTGAGCAAAAGAAGATGTCATTTCAATTACCTTTTT |
| edge | 25[12]24[12] | TTTTCAAATATATTTTAGTGCGAGAAAACTTTTTTTTT |
| edge | 27[12]26[12] | TTTTGTTTATCAACAATAGATGCAGAACGCGCCTTTTT |
| edge | 29[12]28[12] | TTTTACGCTAACGAGCGTCTCCTGAATCTTACCATTTT |
| edge | 31[12]30[12] | TTTTGAAGGAAACCGAGGACGAACAAAGTTACCATTTT |
| edge | 33[12]32[12] | TTTTTAGCAAGGCCGGAAATAGCACCATTACCATTTTT |
| edge | 35[12]34[12] | TTTTTCATTAAAGCCAGAACAAACAAATAAATCCTTTT |
| edge | 37[12]36[12] | TTTTACCGCCACCCTCAGAACCGCCACCCTCAGATTTT |
| edge | 39[12]38[12] | TTTTTCGGTTTATCAGCTTGAGCCTTTAATTGTATTTT |
| edge | 41[12]40[12] | TTTTTACCAAGCGCGAAACGACCCCCAGCGATTATTTT |
| edge | 42[39]42[8] | GGTTTAATTTCAACTTTAATCATTTTTT |
| edge | 20[167]21[198] | AAGGAATTGAATAATGGAAGGGTTAGAACCTACCATATCTTTT |
| edge | 2[195]1[195] | TTTTGAATTACGAGGCATAACATAACGCCAAAAGTTTT |
| edge | 4[195]3[195] | TTTTCAGACCGGAAGCAAAAGCTTCAAAGCGAACTTTT |
| edge | 6[195]5[195] | TTTTAAGGCAAAGAATTAGATAAATCATACAGGCTTTT |
| edge | 8[195]7[195] | TTTTATGAACGGTAATCGTCAAACAAGAGAATCGTTTT |
| edge | 10[195]9[195] | TTTTTTTGAGGGGACGACGCCGTGCATCTGCCAGTTTT |
| edge | 12[195]11[195] | TTTTAATTCCACACAACATTTGTTATCCGCTCACTTTT |
| edge | 14[195]13[195] | TTTTAAGAATAGCCCGAGATCCCTTATAAATCAATTTT |
| edge | 16[195]15[195] | TTTTGGCGCGTACTATGGTAATGCGCCGCTACAGTTTT |
| edge | 18[195]17[195] | TTTTTCTGAAATGGATTATTTTGACGCTCAATCGTTTT |
| edge | 20[195]19[195] | TTTTTCAGTTGGCAAATCATCAATCAATATCTGGTTTT |
| edge | 22[198]22[168] | TTTTTATTTGCACGTAAAACAGAAATAA |
| edge | 24[195]23[195] | TTTTAAACATAGCGATAGCTCCCTTAGAATCCTTGATTTT |
| edge | 26[195]25[195] | TTTTTCAACAGTAGGGCTTAAGTATAAAGCCAACGCTTTT |
| edge | 28[195]27[195] | TTTTGGAATCATTACCGCGCTTTTATTTTCATCGTATTTT |
| edge | 30[195]29[195] | TTTTAACTGAACACCCTGAAATTAGACGGGAGAATTTTTT |
| edge | 32[195]31[195] | TTTTGAAAATTCATATGGTTTTTGTCACAATCAATATTTT |
| edge | 34[195]33[195] | TTTTAGAGCCACCACCGGAACAAAATCACCGGAACCTTTT |

| function | position | sequence |
|---|---|---|
| edge | 36[195]35[195] | TTTTTATTCTGAAACATGAACTATTTCGGAACCTATTTTT |
| edge | 38[195]37[195] | TTTTTCGTCTTTCCAGACGTACGATCTAAAGTTTTGTTTT |
| edge | 40[195]39[195] | TTTTGCATCGGAACGAGGGTTCAGCAGCGAAAGACATTTT |
| edge | 42[195]41[195] | TTTTGCGCATAGGCTGGCTGACGGTGTACAGACCAGTTTT |
| basic | 1[56]2[72] | GTCAGGACGTTGGGAAGAAAAATCTAATAGTA |
| basic | 1[88]2[104] | ATAAAACGAACTAACGGAACAACAGCTTTTGC |
| basic | 1[152]2[168] | ATACCACATTCAACTAATGCAGATGTAAGAGC |
| basic | 1[120]2[136] | GGTAGAAAGATTCATCAGTTGAGCAGACGAC |
| basic | 2[39]42[40] | GGAATCGTTTAAACTCATCTTTAAAGTACAACGGAGATCTTGAGAT |
| basic | 2[71]38[40] | AAATGTTTAAATCAAAGGTTTAGTACCGCCACCCTCAGAGAAAAAGGC |
| basic | 2[103]34[40] | AAAAGAAGTATAGTCAATCACCAGCGTCACCAATGAAACCTTGGCCTT |
| basic | 3[88]1[87] | CTGACTATTTTTGCCAGAGGGGGTACGTTA |
| basic | 3[56]1[55] | ATGACCATAGACTGGATAGCGTCCTTATACCA |
| basic | 3[152]1[151] | CTTCAAATCATAACCCTCGTTTACATTTAGGA |
| basic | 3[120]1[119] | CATCAAAAACCAAAATAGCGAGAGTTATTACA |
| basic | 4[39]6[40] | GGTGTCTAATTCTGCGAACGAGCATATATT |
| basic | 4[103]6[104] | TAGAGCTTTGTTTAGCTATATTTTCCTGTAAT |
| basic | 4[135]6[136] | TTTTGATTGAAAAGGTGGCATCGCTAAATC |
| basic | 4[71]6[72] | CTCAACATATTAGATACATTTCGCTTCAACGC |
| basic | 4[167]6[168] | GGTCAGGAGTAGCATTAACATCCACAAAATTA |
| basic | 5[88]3[87] | CAATAACCAATTGCTGAATATAACTTTACC |
| basic | 5[56]3[55] | GTTTGACCGTTTTAAATATGCAACAAACGAGA |
| basic | 5[120]3[119] | GCGCGAGCAAGAGGTCATTTTTGCGCGGATTG |
| basic | 5[152]3[151] | TAATAGTATTAGAGAGTACCTTTACCGAAAGA |
| basic | 6[71]38[72] | AAGGATAACACCATCAAGCCCGGACAGGGATAGCAAGCCTAATAAT |
| basic | 6[39]42[72] | TTAAATGAAAGGGTAACGAAAGTAAATTGTGTCGAAACGAGAAA |
| basic | 6[103]34[72] | ACTTTTGCTGATAAATACTTGAGCTAATCAGTAGCGACACATTGAC |
| basic | 7[120]5[119] | AGCTATTTCCAAAAACATTATGACCATTTGGG |
| basic | 7[152]5[151] | AGGTCATTAGCCTCAGAGCATAAAAATTCTAC |
| basic | 7[56]5[55] | AGTCAAATAAATTTTTAGAACCCTTAGATTTA |
| basic | 7[88]5[87] | GTTCTAGCGGGAGAAGCCTTTATAAATGGT |
| basic | 8[103]10[104] | AGCAAATACGTCGGATTCTCCGTGAACCAGGC |
| basic | 8[71]10[72] | TTGTTAAACATCAACATTAAATGTTGCGCAAC |
| basic | 8[39]10[40] | CAGCTCAATTCGCGTCTGGCCTGCCTCTTC |
| basic | 9[88]7[87] | TAACAACCTTTAAATTGTAAACGTTCAACC |
| basic | 9[56]7[55] | CCAGCTTTATTCGCATTAAATTTTCCGGAGAC |
| basic | 10[167]26[104] | GGCCTCAGTGTTTCCTACCTCCGGTAAATAAGAATAAACAGTAATAAG |
| basic | 10[39]42[104] | GCTATTAGCGATTAACGTAATGTGTTACTTAGCCGGAAGCTGCTCA |
| basic | 10[135]30[104] | CCGGCACCCGAGCTTCCCGACTAGAAACGATTTTTTGTCCAATAAT |
| basic | 10[71]38[104] | TGTTGGGACAGTCACGTAAGTGCCACCGTAACACTGAGTTGAATAGAA |
| basic | 10[103]34[104] | AAAGCGCCCCAAGCTTATTCATTATAGCGTCAGACTGTAGGCCACCAG |
| basic | 11[120]9[119] | CCCGGGTACGCTTCTGGTGCCGGAGGAACAAA |
| basic | 11[56]9[55] | GGTTTTCCAGGGCGATCGGTGCGGTCCTGTAG |
| basic | 11[152]9[151] | GTCATAGCGAAGATCGCACTCCAGAGGTCACG |
| basic | 11[88]9[87] | GGCCAGTGATTCGCCATTCAGGCGAGCGAG |
| basic | 12[135]14[136] | TGAGTGACCAGCAGGCGAAAATAGTCCACT |
| basic | 12[103]14[104] | GCTCACTGGAGTTGCAGCAAGCGGTCAAAGGG |
| basic | 12[167]14[168] | GAAGCATATCCGAAATCGGCAAAATAGGGTTG |
| basic | 13[88]11[87] | CCCTGAGACCCGCTTTCCAGTCGAAACGAC |
| basic | 13[152]11[151] | ATGGTGGTAAGTGTAAAGCCTGGGTAATCATG |
| basic | 13[56]11[55] | GGGCAACAAGCTGCATTAATGAATAACGCCAG |
| basic | 13[120]11[119] | GGTTTGCCGCTAACTCACATTAATTGAGGATC |
| basic | 14[135]30[136] | ATTAAAGCAAGTGTATTCTAAGAAAATAGCAGCCTTTAAGAGAGAT |

| function | position | sequence |
|---|---|---|
| basic | 14[103]34[136] | CGAAAAACGAAAGCGAAGAGGGAGGATTTTCGGTCATAGCCCGCCACCC |
| basic | 14[167]26[136] | AGTGTTGTCACACCCGAAAATCATCTAGAAAAAGCCTGTTATGTAATT |
| basic | 15[120]13[119] | GGCGCTGGAACGTGGACTCCAACGTCCACGCT |
| basic | 15[152]13[151] | GTAACCACTCCAGTTTGGAACAAGCCTGTTTG |
| basic | 16[167]18[168] | CGAGCACGATGGAAATACCTACATTTACATTG |
| basic | 16[103]18[104] | AAGGGATTTCGGCCTTGCTGGTAATAGAACCC |
| basic | 16[135]18[136] | TCAGAGCACCGCCAGCCATTGCGTAATAAA |
| basic | 16[71]18[72] | ATCCTGAGATCACTTGCCTGAGTAACGTGGCA |
| basic | 16[39]18[40] | CACCGAGTTGTAGCAATACTTCTAGTCTTT |
| basic | 17[56]15[55] | GTAATAACAAGTGTTTTTATAATCAGAGCTTG |
| basic | 17[88]15[87] | TCAAACTATTAGACAGGAACGGTGAGAAAG |
| basic | 17[120]15[119] | ACAATATTGGGAGCTAAACAGGAGGGCGCTAG |
| basic | 17[152]15[151] | AAACGCTCTATAACGTGCTTTCCTCGCTGCGC |
| basic | 18[135]30[168] | AGGGACAATGAAAACAAGCAAAAAAACAGGGAAGCGCCAAAGTC |
| basic | 18[103]34[168] | TTCTGACCTGCAACAGGCCAAAGACCATCTTTTCATAATCCGCCTC |
| basic | 18[167]26[168] | GCAGATTCCTCAAATAAGACGCTGATACAAATTCTTACCATTGAGA |
| basic | 19[152]17[151] | TGCTGAACACCAGTCACACGACCAAACAGGAA |
| basic | 19[56]17[55] | AGATAAAAATTTTTGAATGGCTATTTTGATTA |
| basic | 19[120]17[119] | AGCAGCAATTCTGGCCAACAGAGATATCCAGA |
| basic | 19[88]17[87] | ACACCGCCTGAAAGCGTAAGAATGAAGAAC |
| basic | 20[135]21[151] | TAGGAGCATATAATCCTGATTGTTTGGATT |
| basic | 20[71]21[87] | GTATTAGAAAACCACCAGAAGGAGCGGAATT |
| basic | 20[103]21[119] | GTCAATAGATTCCTGATTATCAGATGATGGCA |
| basic | 20[39]21[55] | TATTAAATTGAGTAACATTATCATTTTGCG |
| basic | 21[56]19[55] | GAACAAAGCTTTACAAACAATTCGCCAGCAGA |
| basic | 21[152]19[151] | ATACTTCTGAGGAAGGTTATCTAACATCACCT |
| basic | 21[120]19[119] | ATTCATCAACTAACAACTAATAGATGAGAGCC |
| basic | 21[88]19[87] | ATCATCATATAATACATTTGAGGAGTATTA |
| basic | 22[167]20[168] | AGAAATTGCTATTAATTAATTTTTAGATTATCAAACCCACAGTTGA |
| basic | 22[103]12[168] | CGGGAGAATTAATGGAAACAGTACCCTTTTTAGTGTGAAAACGAGCCG |
| basic | 22[135]16[168] | GATGAATATGAGTGAATAACCTTGAATTTATCCCGCGCTTTGCTTTGA |
| basic | 24[87]22[72] | GGGTTATAATTTGAATTACCTTTTACAATAACGGATTCGCCTGATTGC |
| basic | 24[119]22[104] | AGAGACTAATAAATCAATATATGTACAGTAACAGTACCTTTTACAT |
| basic | 26[55]24[56] | GACAATAACCTAAATTTAATGGTTGCTGATGC |
| basic | 26[119]24[120] | TCGAGCCACCGGAATCATAATTAAGGTCTG |
| basic | 26[151]24[152] | ACGCCAACTAGTATCATATGCGTTAGAAGAGT |
| basic | 26[167]20[136] | ATCGCCATGAACAAGCAAGCCGTCCAATAGATCTAAAGAATATCTT |
| basic | 26[87]24[88] | ACAAAAGGGTGATAAATAAGGCGTCTTAGGTT |
| basic | 26[103]12[136] | AGAATATATAATCGGCTGTCTTTCAGCGAACCCGAATTCGGTGCCTAA |
| basic | 26[135]16[136] | TAGGCAGAACGGGTATTAAACCAATATCCGGTAGCGGTCACGTTAGAA |
| basic | 28[87]26[88] | GTTTTGAATGTAGAAACCAATCAAAAGTACCG |
| basic | 28[55]26[56] | ATTTTGCAAAAAATAATATCCCATCAGACGAC |
| basic | 28[151]26[152] | AGAAGGCTGTACCGCACTCATCGAATTTAACA |
| basic | 28[119]26[120] | GGCGTTTTCTTATCATTCCAAGAGGCATTT |
| basic | 30[71]8[104] | ATCTTACCGGCATGATTAAGACTGTCACCGTAATGCCGATTGTATA |
| basic | 30[135]16[104] | AACCCACAGTGGCAACATATAAAAAACCGATTAGGAGCGGCCGATTA |
| basic | 30[119]28[120] | AGTTAAGCTTAACGTCAAAAATGAACGCGA |
| basic | 30[39]4[104] | CAGATAGCAACGCAATAATAACGGCCAGCAAAGAAGCAAAGGATGGCT |
| basic | 30[55]28[56] | TTTTAAGATGCCAGTTACAAAATATAGTTGCT |
| basic | 30[167]20[104] | AGAGGGTAACGGAATAAGTTTATTACCAGCTGCCACGCTTAGAGCC |
| basic | 30[151]28[152] | CTAATATCCAGAGAGAATAACATATCAGATAT |
| basic | 30[103]12[104] | AAGAGCAAGTTAGCAAACGTAGAACGGAAATTCTCACAGTTGCGTTGC |
| basic | 32[55]30[56] | AATTAGAGAATACCCAAAAGAACTGAAGCCCT |

| function | position | sequence |
|---|---|---|
| basic | 32[151]30[152] | CGACATTCGAAACGCAAAGACACCATTGAGCG |
| basic | 34[87]32[88] | GCCGCCAGGAATCAAGTTTGCCTTAAGGTGAA |
| basic | 34[103]12[72] | AACCACCAGGAGTGTACTGGTAATCAGGCGGAACGTTGTAGGAAACCT |
| basic | 34[151]32[152] | CTCAGAACCCCTTATTAGCGTTTGCAAAAGGG |
| basic | 34[39]4[72] | GATATTCATGGAAAGCGCAGTCTCACTCAGGAAATCAGGTTGCTGTAG |
| basic | 34[55]32[56] | TCAGACGAATCGATAGCAGCACCGCATTTGGG |
| basic | 34[119]32[120] | TCAGAGCCCGCGTTTTCATCGGCGAAGGTA |
| basic | 34[71]8[72] | AGGAGGTTGTAAGCGTCATACATTAAGTATATATGATATTAATATT |
| basic | 36[55]34[56] | ATCACCGTTGAATTTACCGTTCCAGAGGCAGG |
| basic | 36[87]34[88] | GGTTGATAGGCTTTTGATGATACACCAGAGCC |
| basic | 36[151]34[152] | CAAGAGAAAGTGCCCGTATAAACACCGCCACC |
| basic | 36[119]34[120] | CTCAGTACAAGTTTTAACGGGGTACCACCC |
| basic | 38[71]8[40] | TTTTTCACACCGATAGTTGCGCCAACCTAAGAGAAAGGTGTTAAAT |
| basic | 38[151]36[152] | GATTTTGCCATTCCACAGACAGCCAGACTCCT |
| basic | 38[119]36[120] | CGGAGTGATCGTCACCAGTACAAGGTTTTG |
| basic | 38[39]4[40] | TCCAAAAGGCTTTCGAGGTGAATTTAAAACACAGTTCAGATAAAGTAC |
| basic | 38[87]36[88] | ATTGCGAACAATAGGAACCCATGTGTCGAGAG |
| basic | 38[55]36[56] | TCTCCAAACCACCACCCTCATTTTATAGGTGT |
| basic | 38[103]12[40] | AGGAACAAATCGCCCACGCATAACGGTAAAATAGTTGGGTCGGCCAAC |
| basic | 40[119]38[120] | ATTAAACGCGATATATTCGGTCGGTTTCAG |
| basic | 40[87]38[88] | AAGGCACCGACAATGACAACAACCCTAAAGGA |
| basic | 42[71]40[56] | CACCAGAACGAGTAGTAAATTGGGTTGTATCATCGCCTGAAGGCAAAA |
| basic | 42[103]40[88] | TTCAGTGAATAAGGCTTGCCCTGATCCGCGACCTGCTCCACCACTACG |
| basic | 42[135]40[120] | ATTACCCAAATCAACGTAACAAACGAGGCGCAGACGGTAGTTTCC |
| basic | 42[167]40[152] | TCAAGAGTAATCTTGACAAGAACCGAACTGACCAACTTTGGGCTTTGA |

**Table S2:** Specific DNA sequences for different stator variants.

| function | position | sequence |
|---|---|---|
| **all variants** | | |
| inner, outer, dual | 12[39]13[55] | GCGCGGGGTTTTTCTTTTCACCAGTGAGAC |
| inner, outer, dual | 15[56]14[40] | ACGGGGAACCATCACCCAAATCAAGTTTTTTG |
| inner, outer, dual | 12[71]13[87] | GTCGTGCCGCTGATTGCCCTTCACCGCCTGG |
| inner, outer, dual | 15[88]14[72] | GAAGGGAACGTCTATCAGGGCGATGGCCCAC |
| inner, outer, dual | 9[120]10[136] | CGGCGGATTGACCGTAATGGGATCCAGCTTT |
| inner, outer, dual | 8[135]7[119] | AGAAAAGCCCCAAAAACAGGAAGGAGAGGGT |
| inner, outer, dual | 9[152]10[168] | TTGGTGTAGATGGGCGCATCGTAAACAGTATC |
| inner, outer, dual | 8[167]7[151] | CATGTCAATCATATGTACCCCGGTAGGCTATC |
| inner, outer, dual | 26[39]28[40] | TTCAGCTAATAAGTCCTGAACAAGCCCAGCTA |
| inner, outer, dual | 26[71]28[72] | ATTCTGTCCCTAATTTACGAGCAGCCTTAA |
| inner, outer, dual | 34[135]36[136] | TCAGAGCCCAGTGCCTTGAGTAACGGATTAGG |
| inner, outer, dual | 3[136]4[136] | GAGGAAGCATTGCTCC |
| inner, outer, dual | 34[167]36[168] | CCTCAGAGGTTAATGCCCCCTGCAGTATTA |
| **inner anchor** | | |
| inner anchor | 14[39]15[39] | GGGTCGAGGGAGCC |
| inner anchor | 18[39]19[39] | AATGCGCATACCGA |
| inner anchor | 22[39]24[40] | CGAATTATATGAAACAAACATCAATCGCAAGA |
| inner anchor | 24[39]26[40] | CAAAGAACTAATTTCATCTTCTGAACAACATG |
| inner anchor | 19[40]20[40] | ACGAACCAACAACTCG |
| inner anchor | 22[71]24[72] | TTTGAATAACATTTAACAATTTCTAACTAT |
| inner anchor | 24[71]26[72] | ATGTAAATTGAAATACCGACCGTTAAAGTA |

| function | position | sequence |
|---|---|---|
| inner anchor | 38[135]40[136] | TTTCAACACTGAGGCTTGCAGGGAGACTTTTT |
| inner anchor | 40[135]42[136] | CATGAGGACAATCATAAGGGAACCGGATATTC |
| inner anchor | 2[167]3[167] | AACACTATATCGCGTT |
| inner anchor | 6[167]7[167] | AGCAATAAGCCTGAGA |
| inner anchor | 38[167]40[168] | TGAATTTTGCGGGATCGTCACCCAGCAACG |
| inner anchor | 40[167]42[168] | GCTACAGAAAAGAGGACAGATGAACCTTCA |
| inner anchor | 3[168]4[168] | TTAATTCGCTCCAACA |
| inner anchor | 7[168]8[168] | GTCTGGAGAAAACTAG |
| inner anchor | 14[71]38[136] | TACGTGAAAGCCGGCGtttttcagcaccttttATTAGCGGACTACAACGCCTGTAGTAAACAAC |
| inner anchor | 18[71]38[168] | CAGACAATCAGAGGTGtttttcagcaccttttAGAGGCTGCTCATAGTTAGCGTATAGTAAA |
| inner anchor | 2[135]30[40] | GATAAAAAGATTAAtttttcagcaccttttCAATTTTATTTCCAGAGCCTAATTAAAGTAAG |
| inner anchor | 6[135]30[72] | GGTTGTATTGAGAGtttttcagcaccttttATCAAGATAACAGCCATATTATTAATAGCT |

**outer anchor**

| function | position | sequence |
|---|---|---|
| outer anchor | 28[39]30[40] | CAATTTTATTTCCAGAGCCTAATTAAAGTAAG |
| outer anchor | 14[71]15[71] | TACGTGAAAGCCGGCG |
| outer anchor | 18[71]19[71] | CAGACAATCAGAGGTG |
| outer anchor | 28[71]30[72] | ATCAAGATAACAGCCATATTATTAATAGCT |
| outer anchor | 19[72]20[72] | AGGCGGTCATTTAGAA |
| outer anchor | 2[135]3[135] | GATAAAAAGATTAA |
| outer anchor | 6[135]7[135] | GGTTGTATTGAGAG |
| outer anchor | 36[135]38[136] | ATTAGCGGACTACAACGCCTGTAGTAAACAAC |
| outer anchor | 7[136]8[136] | ATCTACAATGATAATC |
| outer anchor | 36[167]38[168] | AGAGGCTGCTCATAGTTAGCGTATAGTAAA |
| outer anchor | 22[39]24[40] | CGAATTATATGAAACAAACATCAATCGCAAGA |
| outer anchor | 38[135]40[136] | TTTCAACACTGAGGCTTGCAGGGAGACTTTTT |
| outer anchor | 38[167]40[168] | TGAATTTTGCGGGATCGTCACCCAGCAACG |
| outer anchor | 15[72]16[72] | AACGTGGCACGCCAGA |
| outer anchor | 15[40]16[40] | CCCGATTTAGTGAGGC |
| outer anchor | 19[40]20[40] | ACGAACCAACAACTCG |
| outer anchor | 7[168]8[168] | GTCTGGAGAAAACTAG |
| outer anchor | 3[168]4[168] | TTAATTCGCTCCAACA |
| outer anchor | 14[39]42[136] | GGGTCGAGGGAGCCtttgtatggtgtctagtctgatgctgatttCATGAGGACAATCATAAGGGAACCGGATATTC |
| outer anchor | 18[39]42[168] | AATGCGCATACCGAtttgtatggtgtctagtctgatgctgatttGCTACAGAAAAGAGGACAGATGAACCTTCA |
| outer anchor | 2[167]26[40] | AACACTATATCGCGTTtttgtatggtgtctagtctgatgctgatttCAAAGAACTAATTTCATCTTCTGAACAACATG |
| outer anchor | 6[167]26[72] | AGCAATAAGCCTGAGAtttgtatggtgtctagtctgatgctgatttATGTAAATTGAAATACCGACCGTTAAAGTA |

**dual anchor**

| function | position | sequence |
|---|---|---|
| dual anchor | 22[39]24[40] | CGAATTATATGAAACAAACATCAATCGCAAGA |
| dual anchor | 15[40]16[40] | CCCGATTTAGTGAGGC |
| dual anchor | 15[72]16[72] | AACGTGGCACGCCAGA |
| dual anchor | 38[135]40[136] | TTTCAACACTGAGGCTTGCAGGGAGACTTTTT |
| dual anchor | 38[167]40[168] | TGAATTTTGCGGGATCGTCACCCAGCAACG |
| dual anchor | 7[168]8[168] | GTCTGGAGAAAACTAG |
| dual anchor | 14[39]42[136] | GGGTCGAGGGAGCCtttgtatggtgtctagtctgatgctgatttCATGAGGACAATCATAAGGGAACCGGATATTC |
| dual anchor | 18[39]42[168] | AATGCGCATACCGAtttgtatggtgtctagtctgatgctgatttGCTACAGAAAAGAGGACAGATGAACCTTCA |
| dual anchor | 2[167]26[40] | AACACTATATCGCGTTtttgtatggtgtctagtctgatgctgatttCAAAGAACTAATTTCATCTTCTGAACAACATG |
| dual anchor | 6[167]26[72] | AGCAATAAGCCTGAGAtttgtatggtgtctagtctgatgctgatttATGTAAATTGAAATACCGACCGTTAAAGTA |
| dual anchor | 14[71]38[136] | TACGTGAAAGCCGGCGtttttcagcaccttttATTAGCGGACTACAACGCCTGTAGTAAACAAC |
| dual anchor | 18[71]38[168] | CAGACAATCAGAGGTGtttttcagcaccttttAGAGGCTGCTCATAGTTAGCGTATAGTAAA |
| dual anchor | 2[135]30[40] | GATAAAAAGATTAAtttttcagcaccttttCAATTTTATTTCCAGAGCCTAATTAAAGTAAG |
| dual anchor | 6[135]30[72] | GGTTGTATTGAGAGtttttcagcaccttttATCAAGATAACAGCCATATTATTAATAGCT |

**Table S3:** DNA sequences for the 6-helix bundle rotor module. The sequences are based on remaining 828 nt of the the 8064 nt scaffold.

| function | position | sequence |
|---|---|---|
| rotor | [146]1[153] | CTCATTTTGATGAAGGGTAAAACGGCAGATCAGCG |
| rotor | [154]3[167] | GG GTCATGGTGCCCCCTGTTCTTCGCGT |
| rotor | [133]3[146] | TGGAGGTTCCAGCGGGGTTTCTGCCAGC |
| rotor | [153]0[140] | CCTGCATCAGACGAGTCCAGCCACCGTC |
| rotor | [139]3[132] | GGTGGTGCCTTTAGGCCGCCAGCAGTTGATACCGG |
| rotor | [216]1[217] | TGTTCAGCAAATCGATGAGCCGGGTCACT |
| rotor | [216]3[195] | GTGCCGGACTTGTAGAACGTCTTTTCGTGGATAGCTCTCACGCCACGGG |
| rotor | [132]3[125] | CAGTGTCACTGCGCGCCTGTGCCGTTTTCACGGTC |
| rotor | [167]2[175] | CGGATCAGATGCTGATTGCCGGCAGCAAGCTTTCGCACTCAATTACCTG |
| rotor | [160]3[153] | AATGCCAGTTAAACAACTTAAATTTCTGACGCGTG |
| rotor | [174]0[161] | CAGCCAGCGGTGCCTGCAGGCCCGCAAG |
| rotor | [181]3[188] | CTGGTCATTCCGGCAGACGCAGAAACAGCCGTGAGCCGAGGTGGAGCCG |
| rotor | [196]3[209] | TTGCGGTTTAACGGAACGGATAACCTCA |
| rotor | [188]1[195] | GAAAAAGAAACGCGGTCCGTTAGCGTGGGGCGCGG |
| rotor | [195]0[182] | CATCAGATGCCGGGTCCGCCGTGCTGGT |
| rotor | [209]5[216] | GGTGAAGCTCGTCGCTGGCAG |
| rotor | [118]5[118] | GTGTACATCGACATAATTTTAAAATCCCGTAAAAAA |
| rotor | [111]3[111] | CACTCTGTGGTGCTGCTTTTGCCAGAATGCGGCGGG |
| rotor | [125]1[132] | GGCGGTTAGCCGCACAGGCGGCCATCCCACGCATTTTCCAGCTTACGGC |
| interface extension | [218]5[244] | GTTGCCACTGGTGCAATCGGCGAAACGTGAGAGACCAGAGCACAGAGTCTGGAGC |
| interface extension | [258]3[237] | GTAAGCGTCATACCAGTCCCGGAATTTGTACAGCG |
| interface extension | [237]1[261] | TAAACATCCCTTACCTGCGGCTGGTAATGGGTAAAGGTTTTTGCGGGATCG |
| interface extension | [244]3[216] | GGTCATTGCCTGATCCTCATAACGGAACCCTCCGGTAGACTTTCTCCGTCCGGAAA |
| interface extension | [261]6[224] | TTAAAGGCCGCTCTTTGCTCGTCACCATGTTTA |

**Table S4:** DNA sequences for the 6-helix bundle used as rotor extension[3]. The sequences are based on the 7249 nt scaffold (M13mp18).

| function | position | sequence |
|---|---|---|
| interface rotor | 3[100]3[125] | CATGGCTTTTGATGATACAGGAGTGT |
| interface rotor | 0[118]4[112] | AGAGATCATGAACGCGTTCCA |
| interface rotor | 2[132]2[115] | GCTGAGGCTTGCAGGGAG |
| interface rotor | 1[115]0[119] | TCACCCTCAGCATTTTTG |
| interface rotor | 3[110]4[119] | GATGATACAGGAGTGTAATTTAC |
| interface rotor | 0[139]4[126] | CGGAGAGAGCATGTAAGCGCAGTCTCTG |
| interface rotor | 3[126]2[115] | ACTGGTAGCTGAGGCTTGCAGGGAG |
| interface rotor | 1[115]4[112] | TCACCCTCAGCATTTTTGAGAGATCATGAACGCGTTCCA |
| interface rotor | 5[119]1[132] | GTAATCGTAAAACTGGTAGCTAGCGAAA |
| interface rotor | 5[98]0[98] | AAACAAGAGAATCGTACAAAGGCTATCA |
| fluorophore strand | all anchors | Atto655–atatgtacgggtctcactta |
| fluorophore anchor | 1[1183]3[1196] | taagtgagacccgtacatatAATGGATAGTGCCACAAGAACGGGTATTtaagtgagacccgtacatat |
| fluorophore anchor | 0[1189]3[1182] | taagtgagacccgtacatatGAAGGGAGTTCCAGCGCTAACGAGCGTCATCATTCtaagtgagacccgtacatat |
| fluorophore anchor | 0[1168]3[1161] | taagtgagacccgtacatatGCGGGCGCCCGAGAAGCCTAATTTGCCAAATAATCtaagtgagacccgtacatat |
| fluorophore anchor | 0[1126]3[1119] | taagtgagacccgtacatatGTAACCAGGTTCCGTCCCAATCCAAATAAATAATAtaagtgagacccgtacatat |
| fluorophore anchor | 1[1120]3[1133] | taagtgagacccgtacatatCCAGCCATATCAAATCCCATCCTAATTTtaagtgagacccgtacatat |
| fluorophore anchor | 1[1162]3[1175] | taagtgagacccgtacatatTTTTGACAGCAAATGGCTGTCTTTCCTTtaagtgagacccgtacatat |
| fluorophore anchor | 0[1210]3[1203] | taagtgagacccgtacatatCCGGCGACCACTATTTTTATCCTGAATCAAACCAAtaagtgagacccgtacatat |
| fluorophore anchor | 1[1141]3[1154] | taagtgagacccgtacatatACGCTCAATCACCTTGTAGAAACCAATCtaagtgagacccgtacatat |
| fluorophore anchor | 0[1147]3[1140] | taagtgagacccgtacatatAGTGTAGCCTTATAAATAAACAGCCATAACGAGCAtaagtgagacccgtacatat |

| function | position | sequence |
|---|---|---|
| fluorophore anchor | 1[1267]3[1280] | taagtgagacccgtacatatTGACCTGAACATCGCCAATAGCAAGCAAtaagtgagacccgtacatat |
| fluorophore anchor | 1[1204]3[1217] | taagtgagacccgtacatatGATTCACAGTATTAGTACCGCACTCATCtaagtgagacccgtacatat |
| fluorophore anchor | 1[1246]3[1259] | taagtgagacccgtacatatCCAACAGGAACGAATCGTAGGAATCATTtaagtgagacccgtacatat |
| fluorophore anchor | 0[1273]3[1266] | taagtgagacccgtacatatTGCCGTACCACTACCGACTTGCGGGAGGACCGCGCtaagtgagacccgtacatat |
| fluorophore anchor | 1[1225]3[1238] | taagtgagacccgtacatatGTAATAATAAAACAAGCAAGCCGTTTTTtaagtgagacccgtacatat |
| fluorophore anchor | 0[1231]3[1224] | taagtgagacccgtacatatTTTAGAGTCCAACGCTATTTTGCACCCAGAGAACAtaagtgagacccgtacatat |
| fluorophore anchor | 0[1252]3[1245] | taagtgagacccgtacatatAACCCTAACCGTCTGCCTTAAATCAAGAATTTTCAtaagtgagacccgtacatat |
| fluorophore anchor | 1[1288]3[1310] | taagtgagacccgtacatatCGTGGCATGCGCGAATAGAAGGCTTATCCGGTATTTTtaagtgagacccgtacatat |
| fluorophore anchor | 0[1296]1[1287] | taagtgagacccgtacatatTTAAGTTTTTTGGGAAGAATAtaagtgagacccgtacatat |
| fluorophore anchor | 4[1310]3[1287] | taagtgagacccgtacatatTTTTCTAAGAACGCGAGGCGTTTTAGCGATCAGATtaagtgagacccgtacatat |
| fluorophore anchor | 4[1280]5[1296] | taagtgagacccgtacatatAACCTCCGTGAACCATCACCCAAATCTTtaagtgagacccgtacatat |
| fluorophore anchor | 2[1315]1[1315] | taagtgagacccgtacatatTTTGGCTATTAGTCTTTAACAGACAATATTTTTGAATTtaagtgagacccgtacatat |
| basic | 1[154]3[167] | GCAACGGCGCCCACGCCTTGAGTAACAG |
| basic | 0[139]3[132] | CGGAGAGAGCATGTAAGCGCAGTCTCTGACTGGTA |
| basic | 0[832]3[825] | AAGCAAACGAGCCGAAACGCAATAATAAAGAATAA |
| basic | 1[553]3[566] | AACAAAGAACAAACGAATCCTTGAAAAC |
| basic | 1[1036]3[1049] | CTTTGATAGCACTACGACGACAATAAAC |
| basic | 0[643]3[636] | TATAATGCTGCAAGCAAAAGGGCGACATCTGAGAG |
| basic | 1[280]3[293] | AAACACTTTGAAAATTGCTCAGTACCAG |
| basic | 1[343]3[356] | TATCATCGTTTCAGTGTATCACCGTACT |
| basic | 1[1015]3[1028] | TTAACCGAGAGCCGGGTAAAGTAATTCT |
| basic | 0[412]3[405] | CATACAGGATAGGTGTAGCGCGTTTTCAAACCGCC |
| basic | 0[286]3[279] | AAGGATATAGGAACAACCGCCACCCTCACGGGGTT |
| basic | 0[622]3[615] | GTTTTAAAGCTGGCATTGAGGGAGGGAAGAATTTA |
| basic | 1[322]3[335] | ACAAAGTAAAGGAAATAAGTATAGCCCG |
| basic | 1[448]3[461] | TGAAAGACACAGACAATAGGAACCCATG |
| basic | 0[370]3[363] | CCTCAGATCGGATTTAGCGTTTGCCATCCAGGAGG |
| basic | 0[580]3[573] | TCCATATGTTGGGAAAGGTGAATTATCAATAGCGA |
| basic | 1[133]3[146] | GACAGCATTCGGTCATAAGTTTTAACGG |
| basic | 1[511]3[524] | TAATCTTCATTTGACCTTGCTTCTGTAA |
| basic | 1[532]3[545] | ATTCATTATTAATTCTATTAATTAATTT |
| basic | 1[427]3[440] | GGGAACCGCGTAACATTTTCAGGGATAG |
| basic | 0[244]3[237] | TGCCTGATTCGCATAACCACCACCAGAGTATTAAG |
| basic | 1[1099]3[1112] | ATATCCACTGGTCATCCTGAACAAGAAA |
| basic | 1[889]3[902] | AAAACCATTGCGGACCAGTATAAAGCCA |
| basic | 1[700]3[713] | AGGACGTTTTAACGCCAATCGCAAGACA |
| basic | 0[748]3[741] | CCAGACCCGGGTACCATACATAAAGGTGCAAATAT |
| basic | 1[196]3[209] | AGGAAGTCAGCTTGCCTATTTCGGAACC |
| basic | 1[175]3[188] | AGGACTAGCCGACAATAAACAGTTAATG |
| basic | 0[538]3[531] | TAGTTTGGAAACCATAGAGCCAGCAAAAATCGTCG |
| basic | 0[979]3[972] | TACGCCAGGTTTGCCTAATATCAGAGAGCAGAGGC |
| basic | 0[874]3[867] | ATAAATCGTGAGCTGTAAGCAGATAGCCAGTATCA |
| basic | 0[790]3[783] | AAGACTTCCTGTGTAAGACTCCTTATTATGGTTTG |
| basic | 1[385]3[398] | CATGTTAAATGAATACCGCCACCCTCAG |
| basic | 0[391]3[384] | TAGCAAAACGGCGGTTTCGGTCATAGCCCCTCAGA |
| basic | 0[223]3[216] | AAGATTCAACGTTACAGCATTGACAGGATATTATT |
| basic | 0[916]3[909] | CCCCCTCTTCCAGTGAAATAGCAATAGCACGCTCA |
| basic | 0[853]3[846] | TACCCTGAAAGCCTGTTACCAGAAGGAAATTACTA |
| basic | 1[1057]3[1070] | CTTGCCTGTTATCTTCAGCTAATGCAGA |
| basic | 1[259]3[272] | AAGAGGCGCTCCAAAGGATTAGGATTAG |
| basic | 4[125]1[132] | AATTTACGTAATCGTAAAACTGGTAGCTAGCGAAA |
| basic | 0[181]3[174] | CATCAATCCCAAAAGATATTCACAAACATGCCCGT |
| basic | 0[265]3[258] | CCTCATAATCAGCTCACCCTCAGAGCCGCAAGAGA |

| function | position | sequence |
|---|---|---|
| basic | 0[1063]3[1056] | GAGCACGAGAGAGTGAGAGAATAACATAAACATGT |
| basic | 1[574]3[587] | AATAAGGCTGAGCATAGCTTAGATTAAG |
| basic | 1[595]3[608] | AAACACCCGAATTAGAAGAGTCAATAGT |
| basic | 0[328]3[321] | GACCCTGGCCAGCTAGCCACCACCGGAAGGTTGAT |
| basic | 0[349]3[342] | CGGTTGTATGTGAGAATCAAAATCACCGGAATAGG |
| basic | 1[973]3[986] | TCAGTGAAGACTTTATTTTCGAGCCAGT |
| basic | 0[1105]3[1098] | CTTAATGGCGAAAAATTTTTTGTTTAACAGATAAG |
| basic | 1[301]3[314] | AGCGATTGCGAATAAGTGCCGTCGAGAG |
| basic | 1[826]3[839] | AAAGGAATGGCAATACACCGGAATCATA |
| basic | 1[616]3[629] | ATTGGGCAGTTACATCAAAATCATAGGT |
| basic | 0[664]3[657] | GCTTAGATAACGCCATGGTTTACCAGCGCTCCGGC |
| basic | 1[637]3[650] | TTTCAACGCCTGATACTACCTTTTTAAC |
| basic | 1[994]3[1007] | AGAGTCTATTTGAGGAATATAAAGTACC |
| basic | 0[475]3[468] | GGGCGCGCGACAGTAACCATCGATAGCATACCGTA |
| basic | 1[217]3[230] | TAAAATATTTCGAGCTGAAACATGAAAG |
| basic | 1[1078]3[1091] | AAACTATAGTTGAATGTTTATCAACAAT |
| basic | 0[1042]3[1035] | CTCGTTAGCCCTTCGGAAGCGCATTAGAGTCCAGA |
| basic | 1[931]3[944] | AGAGGGGCCGAACGTCGCCATATTTAAC |
| basic | 0[433]3[426] | TAGCATTTGGGCGCGTTTGCCTTTAGCGCACCCTC |
| basic | 0[1000]3[993] | AGGGATTGGTGGTTAAGTCAGAGGGTAAAATAAGA |
| basic | 0[727]3[720] | ACAGGTCAGGTCGAATAAAAGAAACGCAAAGAACG |
| basic | 1[679]3[692] | AACTGGCAGTAACATATGTAAATGCTGA |
| basic | 0[811]3[804] | AAAGATTCTCACAACCCAAAAGAACTGGTGATAAA |
| basic | 0[307]3[300] | GAGAAGCTTCGCGTCCTCAGAGCCGCCAGCGGATA |
| basic | 1[364]3[377] | GTCGAAATTTGCTATTTAGTACCGCCAC |
| basic | 1[742]3[755] | CGGAACATGCACGTATTTTAGTTAATTT |
| basic | 0[1084]3[1077] | GCGTACTCACGCTGATGAAAATAGCAGCACGCGCC |
| basic | 0[958]3[951] | TAGCGTCGGCCAACACAAGAATTGAGTTAACGCCA |
| basic | 0[937]3[930] | CGTCATATGCCAGCATAATAAGAGCAAGTTGAGAA |
| basic | 1[805]3[818] | TAATGCACTGATTGTAAGGCGTTAAATA |
| basic | 1[910]3[923] | TTTTGCAAGTTTGAACAGTAGGGCTTAA |
| basic | 0[706]3[699] | CCTTTAACCAGTGCCACGGAATAAGTTTTGCAAAT |
| basic | 0[160]3[153] | TTCTAGCCCGGTTGCCTCATTAAAGCCAGGTCAGT |
| basic | 1[406]3[419] | GGCGCAGTCGTCTTACCCTCAGAGCCAC |
| basic | 0[1021]3[1014] | GCTAAACTGAGACGATTAACTGAACACCGACAAAA |
| basic | 1[658]3[671] | ATTACCTGGGAGAATTAGGTTGGGTTAT |
| basic | 1[469]3[503] | TGTACAGACTACAAACACTGAGTTTCGTCAATAAATCAATATATGTGAG |
| basic | 1[721]3[734] | TACGTTAGAAATTGCGAGAAAACTTTTT |
| basic | 1[868]3[881] | CTCGTTTAGAAGGATATGCGTTATACAA |
| basic | 0[517]3[510] | TTCGCAACCGGCACTAGCACCATTACCATGAATAA |
| basic | 0[601]3[594] | TACGGTGGGGCCTCATTGACGGAAATTAACGCTGA |
| basic | 0[685]3[678] | AAGAGGTCACGACGCACAATCAATAGAAATAACTA |
| basic | 0[769]3[762] | TAATTCGTAATCATTGTTAGCAAACGTACATCTTC |
| basic | 1[763]3[776] | AGAAAGAACCTACCTGACCTAAATTTAA |
| basic | 1[847]3[860] | AAGAGCAATATTCCGAAAAAGCCTGTTT |
| basic | 0[559]3[552] | AATTCTGCGCCATTCGACTTGAGCCATTTCCCTTA |
| basic | 0[454]3[447] | CATCAATTCTGCCAAATCAGTAGCGACACAAGCCC |
| basic | 0[202]3[195] | GGCCGGATAAGCAAGCAGGTCAGACGATCCCCCTG |
| basic | 1[784]3[797] | TTTAGGACTGAATAAAATACCGACCGTG |
| basic | 0[496]4[483] | GTTTAGCGATCGCAGGCCGGAAACGTCA |
| basic | 0[895]3[888] | GTTCAGAGCGTTGCCCGAAGCCCTTTTTATTCTTA |
| basic | 1[238]3[251] | GAAGGCATGTATCGAGGCTGAGACTCCT |
| basic | 2[1266]0[1253] | CCATTAAAAATACCAGATAGAAAATCGG |

| function | position | sequence |
|---|---|---|
| basic | 2[1287]0[1274] | ACTGATAGCCCTAAAAAGCGTGTCGAGG |
| basic | 2[153]0[140] | GCATAACCGATATATCGGAACTTAATGC |
| basic | 4[146]1[153] | GAATGGACAATCATATGTACCTGATAAAGAGGGTA |
| basic | 4[1238]1[1245] | TTAGTTGTCAAAGGGCGAAAAAAGGGAGATTCTGG |
| basic | 4[1259]1[1266] | TTTTGAAATCAGGGCGATGGCAAGCACTACCCTTC |
| basic | 4[1217]1[1224] | GCTACAATAAAGAACGTGGACCTTGACGACGACCA |
| basic | 4[167]1[174] | AATAAATATAATCAGAAAAGCATGATATGGCTTTG |
| basic | 2[1245]0[1232] | CCACCAGCAGAAGAAAGGGACCCCCCGA |
| basic | 2[174]0[161] | ATGACAACAACCATCTACAGATCAACCG |
| basic | 2[1224]0[1211] | GAGGTGAGGCGGTCCAGTCACGGGAAAG |
| basic | 2[1203]0[1190] | ACACCGCCTGCAACTATTTACGAGAAAG |
| basic | 4[1196]1[1203] | TTACCAATTTGGAACAAGAGTACGTGGCATTGGCA |
| basic | 2[195]0[182] | ATACCGATAGTTGCAAGACTTAAATCAC |
| basic | 4[188]1[195] | TGGCCTTACAGGAAGATTGTAGACAGTCTTTCATG |
| basic | 4[230]1[237] | CCGCCGCATATTTTGTTAAAAGTAATGTCCACTAC |
| basic | 2[951]0[938] | ATTAAATCCTTTGCGTAATAGGCGGAAT |
| basic | 2[237]0[224] | GTTTATCAGCTTGCCGTAATGGTAGGTA |
| basic | 4[1112]1[1119] | AGAAACGTCCTGTTTGATGGTCCACACCATTACCG |
| basic | 2[552]0[539] | ATCAAGAAAACAAAACCCAAAGTAGATT |
| basic | 4[944]1[951] | AAGCCCATGCATTAATGAATCCAATACTTAAAATG |
| basic | 4[629]1[636] | TCAACCGGAAAGGGGGATGTGCTGTAGCGGTTTAA |
| basic | 2[720]0[707] | CGTAGATTTTCAGGTGGGAAGGAGAGTA |
| basic | 4[587]1[594] | TTCATTAAGGGCGATCGGTGCTCTGGAATGACGAG |
| basic | 2[1140]0[1127] | TGCTGAACCTCAAATTGCAACGCTGCGC |
| basic | 2[1056]0[1043] | AAAATATCTTTAGGTAGTAATTGCTTTC |
| basic | 4[713]1[720] | AAGACACCAAGCTTTGCCTGCAGGATTAAAAAATC |
| basic | 4[1049]1[1056] | AAAACAGACCGCCTGGCCCTGTATAACGAACATCA |
| basic | 4[1091]1[1098] | GTCAAAAGTTTGCCCCAGCAGCGCCGCTGCTGGTA |
| basic | 4[608]1[615] | GGTAAATTTCGCTATTACGCCATATGCAGTAGTAA |
| basic | 2[636]0[623] | TGCTTTGAATACCATTGAGATTCAACAT |
| basic | 2[1077]0[1064] | AGGAATTGAGGAAGGAGTAGACTTTGAC |
| basic | 4[797]1[804] | CATGATTGAAATTGTTATCCGAAGAGGAATTCAAC |
| basic | 2[321]0[308] | CAACTAAAGGAATTATACCAATTTGCGG |
| basic | 2[762]0[749] | ATATCAAAATTATTACATTATAAGCGAA |
| basic | 4[860]1[867] | GAACAAAGGGGTGCCTAATGAAAAAATCCATAACC |
| basic | 4[251]1[258] | CCACCAGTAAATTTTTGTTAATATTTTAAAAACGA |
| basic | 4[671]1[678] | AATTCATAGGGTTTTCCCAGTCATTTTTTTTTAAG |
| basic | 4[818]1[825] | CGGAATATTCCACACAACATAGCGGATTAACGCCA |
| basic | 2[279]0[266] | TCTCCAAAAAAAAGAAAAGAATTAGAAC |
| basic | 4[419]1[426] | TCAGACTCACGTTGGTGTAGAAACATCCATCATAA |
| basic | 2[405]0[392] | TCCAGACGTTAGTACTTAGCCAAAGAAT |
| basic | 4[440]1[447] | GAATCAAATCGTAACCGTGCATCTACTACCAACTT |
| basic | 2[594]0[581] | TTCATTTCAATTACCTTGCCCGTTTCAT |
| basic | 5[959]0[959] | GCGCGGGGAGAGGCGAATCCTGAGAAGA |
| basic | 1[490]0[476] | CTGGCTGTGGAAACAGTACCCAGTACAAACCAGGCTCATTTG |
| basic | 4[545]1[552] | TGGGAATGGCAAAGCGCCATTCGAACGATCAACGT |
| basic | 4[1007]1[1014] | CTGAACATTTCTTTTCACCAGAGGAGGCACGCAAA |
| basic | 2[888]0[875] | ACAAAGAAACCACCACCAGACAATGACC |
| basic | 4[923]1[930] | AAACAATCGGGAAACCTGTCGAATATTCTTTTGCC |
| basic | 4[398]1[405] | TCGGCATATTGACCGTAATGGGCAAGGCGGAACGA |
| basic | 4[902]1[909] | TATCTTAGCTCACTGCCCGCTAAATGCTGAGAGGC |
| basic | 2[300]0[287] | ATAATTTTTTCACGCATCTTTTCAACGC |
| basic | 2[804]0[791] | TTTGGATTATACTTATACCACAGCCCGA |

| function | position | sequence |
|---|---|---|
| basic | 4[1175]1[1182] | TTTCCAGTAGGGTTGAGTGTTAGAAAGCCGTCTGA |
| basic | 2[1161]0[1148] | GAAAAATCTAAAGCTGGAAATGCTGGCA |
| basic | 2[930]0[917] | TTATTAATTTTAAAAAAGAAGATTGAAT |
| basic | 2[573]0[560] | AAAGAAGATGATGACTGCTCAGATTCCC |
| basic | 2[783]0[770] | ATGGAAGGGTTAGATTCATCACGCGTTT |
| basic | 2[867]0[854] | GCGGAATTATCATCACACTATAGGTCTT |
| basic | 2[426]0[413] | GATCTAAAGTTTTGACGGTCAAATAAAT |
| basic | 2[468]0[455] | CGCCTGTAGCATTCGGACAGAAAGGTGG |
| basic | 4[524]1[531] | TCACCAGCGCTTCTGGTGCCGACCATTAACCGGAT |
| basic | 4[209]1[216] | GGTTGAGATATTTAAATTGTAAAAAGGGAAACGGG |
| basic | 2[741]0[728] | AAAACAGAAATAAAATAAAACAACTCCA |
| basic | 4[566]1[573] | CCGTCACCAGGCTGCGCAACTAACAGTTTTCAGTG |
| basic | 4[734]1[741] | GCAACATCTCTAGAGGATCCCGGAAGCAGAACTAA |
| basic | 2[531]0[518] | ACATTTAACAATTTGACAAGAGATACAT |
| basic | 2[615]0[602] | AAATCGCGCAGAGGAGAACGAACTAAAG |
| basic | 2[216]0[203] | GTGAATTTCTTAAATTCCATTTGAGAAA |
| basic | 2[1182]0[1169] | CGCTGAGAGCCAGCGCTCAATGAAAGGA |
| basic | 4[776]1[783] | CGCAGTAGGTCATAGCTGTTTCAAATATGTTGAGA |
| basic | 4[461]1[468] | GCACCGTGTTTGAGGGGACGAAGCTGAATGAACGG |
| basic | 4[272]1[279] | GAGCCACCATTTTTTAACCAAAAAATTTTACACTA |
| basic | 2[846]0[833] | TGATTATCAGATGATTACGAGTAGTCAG |
| basic | 2[972]4[959] | ACAAACAATTCGACAACTCGTACATGTAATTTAGGATAACCC |
| basic | 2[1014]0[1001] | TCAATAGATAATACGTCCATCCGATTAA |
| basic | 4[839]1[846] | ACCGAGGGAAGCATAAAGTGTACTATTAGCATAGT |
| basic | 2[258]0[245] | AAGGAGCCTTTAATCCAACCTAATGCAA |
| basic | 4[314]1[321] | CCGCCTCCTGGCCTTCCTGTATAATACTGCGCGAA |
| basic | 4[1133]1[1140] | TTATTTAAAATCGGCAAAATCCGGTCACAGGAAAA |
| basic | 2[825]0[812] | TCATCAATATAATCGATACATGCATCAA |
| basic | 4[1154]1[1161] | GTTACAAAATCAAAAGAATAGCTAGGGCACCTACA |
| basic | 4[482]1[489] | CCAATGAATCGGCCTCAGGAATATATTTGCATAGG |
| basic | 2[678]0[665] | GTACCTTTTACATCTATGCGAGCGGATG |
| basic | 2[993]0[980] | GATTTAGAAGTATTGGCCACCGGAACGG |
| basic | 2[384]0[371] | TTTCTGTATGGGATTCCGCGAAATAAAG |
| basic | 2[363]0[350] | AACAACTTTCAACAGCCTGATGCTAAAT |
| basic | 2[447]0[434] | AGCCCTCATAGTTAGAACTGAATAGTAG |
| basic | 2[699]0[686] | TCAGATGAATATACTCATTATTTTTGAT |
| basic | 4[881]1[888] | AAGAAAAAAACTCACATTAATTAAACGAGGACGATA |
| basic | 4[986]1[993] | TTGAGCGGTATTGGGCGCCAGTTAGACAGAGTAAA |
| basic | 4[650]1[657] | CCAAAGAGCGATTAAGTTGGGGCTTAATATTGTGA |
| basic | 4[356]1[363] | TTTTCATCGAGTAACAACCCGGCATAAAAAATTGT |
| basic | 1[952]1[972] | TTTAGACTGGTGTTTTTATAA |
| basic | 2[657]0[644] | ACAATAACGGATTCTTTAATCTGCTGAA |
| basic | 2[909]0[896] | GTAACATTATCATTAAATAGCTTAAACA |
| basic | 2[1119]0[1106] | CCCTCAATCAATATGAACAATCGCCGCG |
| basic | 4[377]1[384] | CCCTTATCTCCGTGGGAACAAATTAAGCCCTGCTC |
| basic | 4[335]1[342] | GAACCAGTTCATCAACATTAAACCAAAAAGATTTG |
| basic | 4[1028]1[1035] | CGGGAGAGGCAACAGCTGATTGAATCAGAATACTT |
| basic | 4[503]1[510] | TTAGCAACTCCAGCCAGCTTTATGGTCATCAAGAG |
| basic | 2[342]0[329] | CGGAGTGAGAATAGACAACGGACATTAT |
| basic | 2[510]0[497] | ATTACCTTTTTTAAACCTTCAATAACCT |
| basic | 4[755]1[762] | GAAAATACGAGCTCGAATTCGAGCTTCATACAGGT |
| basic | 4[1070]1[1077] | CTTTACATGCAGCAAGCGGTCATGGTTGAGAACTC |
| basic | 2[1098]0[1085] | GTTGGCAAATCAACCGGCCTTACAGGGC |

| function | position | sequence |
|---|---|---|
| basic | 4[293]1[300] | CCCTCAGGCCATCAAAAATAACTTTATTGACCCCC |
| basic | 4[692]1[699] | ATTTTGTTTGTAAAACGACGGTTGCTCCACCAGTC |
| basic | 2[1035]0[1022] | ACAACTAATAGATTTTGTAGCAGCGGGA |
